# Mobile elements in human population-specific genome and phenotype divergence

**DOI:** 10.1101/2022.03.25.485726

**Authors:** Shohei Kojima, Satoshi Koyama, Mirei Ka, Yuka Saito, Erica H. Parrish, Mikiko Endo, Sadaaki Takata, Misaki Mizukoshi, Keiko Hikino, Atsushi Takeda, Asami F. Gelinas, Steven M. Heaton, Rie Koide, Anselmo J. Kamada, Michiya Noguchi, Michiaki Hamada, Biobank Japan Project Consortium, Yoichiro Kamatani, Yasuhiro Murakawa, Kazuyoshi Ishigaki, Yukio Nakamura, Kaoru Ito, Chikashi Terao, Yukihide Momozawa, Nicholas F. Parrish

## Abstract

Mobile genetic elements (MEs) are heritable mutagens that contribute to divergence between lineages by recursively generating structural variants. ME variants (MEVs) are difficult to genotype, obscuring their impact on recent genome and trait diversification. We developed a tool that uses short-read sequence data to accurately genotype MEVs, enabling us to study them using statistical genetics methods in global human genomes. We observe population-specific differences in the distribution of *Alu* insertions that distinguish Japanese from other populations. We integrated MEVs with epigenomic and expression quantitative trait loci (eQTL) maps to determine how they impact traits. This reveals coherent patterns by which specific MEs regulate tissue-specific gene expression, including creating or attenuating enhancers and recruiting post-transcriptional regulators. We pinpoint MEVs as genetic causes of disease risk, including a LINE-1 insertion linked to keloid and other diseases of fibroblast inflammation, by introducing MEVs into the genome-wide association study (GWAS) framework. In addition to nominating previously-hidden MEVs as causes of human diseases, this work highlights MEs as accelerators of human population divergence and begins to decipher the semantics of MEs.

## Introduction

Mobile genetic elements (MEs) characteristically insert copies of themselves to new genome locations. The evolutionary innovations of MEs are constrained within the linear descent of their host genomes, thus differences in the sequences, mobilization activity, or insertion preferences of the MEs in a particular lineage can increase the rate at which descendant genomes accumulate mutations characteristic of that lineage; in other words, MEs can accelerate genomic divergence. MEs account for a large part of species-specific genomic differentiation (*1*), but the degree to which MEs cause species-level phenotypic differences is difficult to dissect due to accumulation of other genetic variation. MEs may also be a force driving speciation, however direct evidence of within-species divergence driven by MEs is limited (*2*). Human genomes and phenotypes are well studied, offering a unique opportunity to dissect the consequences of MEs on recent and ongoing lineage divergence and phenotypic diversity.

Many of the phenotypes most closely linked to ME replication portend evolutionary dead ends. Mutation of ME-restrictive factors, such as those in the PIWI-interacting RNA pathway, often lead to sterility (*3*). For example, mutations affecting *PIWIL1*, which are reportedly present at higher frequency in East Asians, have been associated with azoospermia in humans (*4*). Insertions of all of the MEs actively replicating in human genomes, namely long interspersed nuclear element 1 (L1), SINE-VNTR-*Alu* (SVA) and *Alu* elements, have been implicated in severe Mendelian diseases (*5*). For example, an SVA insertion so far only reported in the Japanese population causes Fukuyama congenital muscular dystrophy (*6*). ME replication, occurring in a genotype-dependent and lineage-specific manner, can lead to phenotypes detrimental to organismal fitness, providing a powerful basis for the metaphor that MEs and their host genomes are engaged in an arms race.

MEs also influence the complex traits that differentiate humans and human populations, but our view of this landscape remains partial. For example, subjects carrying a *SLCO1B3* allele with exonic insertion of a proposed Japanese-specific highly-active L1 (*7*) develop a benign form of hyperbilirubinemia (*8*). Recent studies have identified ME polymorphisms associated with differential gene expression (*9*–*11*), and differential polygenic disease risk (*12*, *13*), but the global influence on human traits remains unclear. MEs make up a large fraction of DNase hypersensitive sites (*14*), which are enriched in complex trait heritability (*15*), and are also the main source of novel regulatory elements in primate genomes (*16*). Moreover, structural variants (SVs), about a quarter of which are MEVs (*17*, *18*), are frequently in tight linkage disequilibrium (LD) with expression quantitative trait loci (eQTL) and trait-associated variants (*18*, *19*). Actively-replicating MEs necessarily carry promoters and transcription-factor binding sites which drive their expression, and some MEs appear to have been coopted as lineage-specific gene regulatory elements (*20*, *21*). These observations provide a rationale to comprehensively assess the impact of ME polymorphisms on human biology, for example performing ME-oriented genotype-trait association studies using biobanks.

One barrier to ME-phenotype correlation is the low accuracy of current methods used to genotype MEVs, lower than those available for single-nucleotide variants (SNVs) and often too low to derive meaningful hypotheses from statistical genetics approaches. Long-read and strand-specific sequencing are ideal to resolve MEVs and other SVs (*18*, *22*), however, the number of genomes studied using these methods is low, and will remain orders of magnitude lower than those genotyped by short-reads until new enabling technologies emerge (*23*, *24*). Here we report a tool, MEGAnE, that precisely detects and genotypes MEVs from biobank-scale short-read whole-genome sequencing (WGS) datasets. By applying statistical genetics approaches to MEs, we showcase MEs as drivers of genomic and phenotypic innovation in humans and divergence of populations.

## Results

### Development and Benchmarking of MEGAnE

Accurate variant genotyping is required for statistical genetics, especially haplotype estimation and genotype imputation. For common SVs already discovered, sequence-resolved, and assigned to haplotypes using long-read sequencing, this has recently become possible using short-read WGS data (*18*, *25*). To enable both discovery and accurate MEV genotyping from genomes studied using short-reads, we developed a new bioinformatic tool, mobile element genotype analysis environment (MEGAnE). MEGAnE uses reads chimerically mapping to ME and non-ME sequences, hybrid read pairs that map to ME and non-ME sequences, and polyA or polyT containing reads to find MEV breakpoints. In addition to evaluating the number of breakpoint-supporting reads and reads evidencing the insertion-absent allele, it infers genotype using the local read depth at the target-site duplication (TSD) characteristic of ME transpositions (see Supplementary Note).

Compared to SVs resolved by long-reads, MEGAnE discovers ME insertions (MEIs) and ME absences with false-positive rates of 3% and 6%, respectively (Fig. 1A and S5). It discovers more than 80% of the target-primed reverse transcription (TPRT)-mediated insertions that can be found using long-reads, and more than 80% of MEVs are genotyped as accurately as using long-reads or a graph-based genotyper (Fig. S6). Less than 2% of genotype calls are inconsistent with Mendelian inheritance (Fig S9). To test the genotyping quality of MEGAnE by an orthogonal approach, we deep sequenced over 100 MEV target sites using DNA from 2,221 Japanese individuals. More than 95% of genotype calls were concordant with those determined by targeted deep sequencing (Fig. 1B and C and S10-15). Accurate genotyping allows us to assign MEVs to haplotypes better than alternatives (Fig. S16); more than 90% of ME genotypes imputed using MEGAnE’s output were highly concordant with those inferred using graph-based pangenome references (Fig. 1B, S16 and S17). While read length imposes some intrinsic limitations on MEV discovery, the low false positive rate and accurate genotyping of this tool enabled us to interrogate MEVs in short-read data at a resolution that was previously impossible.

**Fig. 1.**
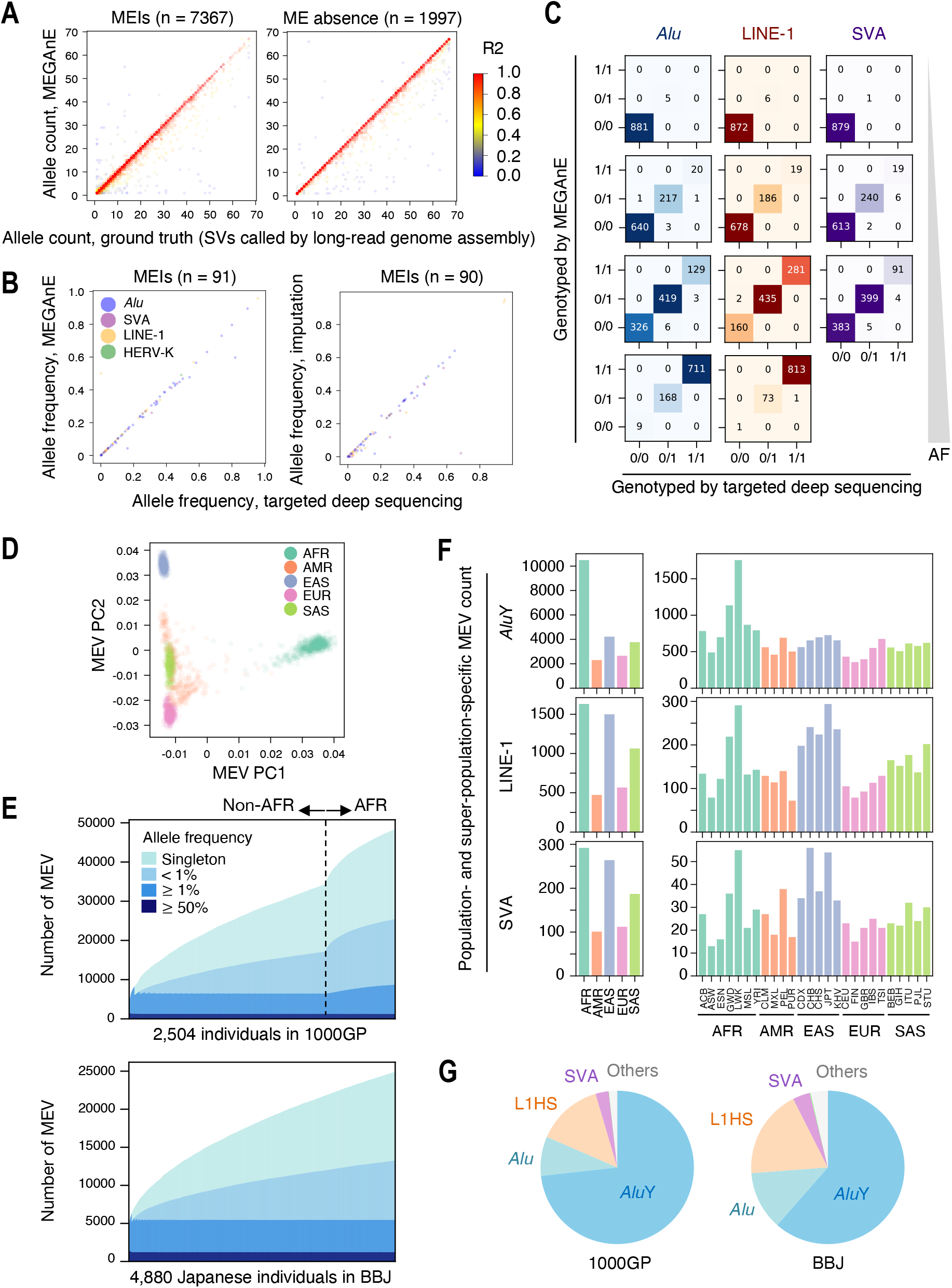
Discovery and accurate genotyping of MEVs in global and Japanese populations. (A) Concordance between MEV genotype called by MEGAnE and an SV callset generated by Phased Assembly Variant caller in 34 individuals. Dot color represents R-squared between the two genotyping results. (B) Concordance between allele frequency called by MEGAnE, or imputed based on MEGAnE calls, and targeted deep sequencing. Genotypes of MEIs in 888 Japanese were directly called by MEGAnE, or imputed using haplotypes in the 1000GP, were compared to those assessed by targeted deep sequencing. 54 *Alu*, 27 L1, 9 SVA, and 1 human endogenous retrovirus (HERV)-K were analyzed. (C) Examples of MEV genotypes called by MEGAnE and targeted deep sequencing. (D) Distribution of first 2 PCs of MEVs discovered in the 1000GP. Color indicates super-population. (E) Discovery of MEVs from diverse populations in the 1000GP (top) and Japanese in BBJ (bottom). The color of bar plots is stratified based on allele frequency of MEVs. (F) The number of super-population-specific (left three panels) and population-specific (right three panels) MEVs found in the 1000GP. (G) Proportion of ME families found in 1000GP (left) and BBJ (right). In this figure, *Alu* represents *Alu* subfamilies other than *Alu*Y.

### Characteristics of MEVs in diverse populations and Japanese

We applied MEGAnE to the 2,504 and 1,235 individuals sequenced at high coverage (30x and 25x) in the 1000 Genomes Project (1000GP) and BioBank Japan (BBJ), respectively. We detected 48,360 and 10,996 MEVs in these respective cohorts, with around 2,500 to 3,000 polymorphisms per individual (Fig. S19). The top 8 principal components (PCs) of MEVs were highly correlated with those of SNVs; like SNVs, MEVs reflect the geographical distribution of human populations (Fig. 1D). MEVs are more abundant in Africans, as are population-specific MEVs (Fig. 1E and F). Population-specific L1 and SVA are more abundant in East Asians, particularly in Japanese, than other non-African populations, whereas the abundance of *Alu* is similar (Fig. 1F). Over half of the MEVs observed as Japanese-specific singletons within 1000GP, which sequenced 104 Japanese subjects, were observed in other subjects in BBJ (Fig. S20). As expected, MEVs predominantly involve young elements known to be active germline mutagens (*Alu*, L1, and SVA) (Fig. 1G and S31).

Fixed MEs enrich in distinct genome regions (*26*). To assess the genomic niches occupied by MEVs, we correlated MEV occurrences with genome features (Fig 2A and B). L1 polymorphisms are positively correlated with markers of heterochromatin, such as DNA methylation and H3K9me3. SVA polymorphisms show the opposite trend, occurring more often in regions with active chromatin markers, such as H3K9Ac and early replication timing. To reduce the degree to which selection may influence this observation, we also analyzed the association with rare, presumably recently-acquired insertions. Singletons found in the 1000GP and BBJ exhibit a similar trend; polymorphisms of L1 and SVA show positive and negative correlations to heterochromatin markers, respectively (Fig. S32A). In addition to singletons, which may have higher false positive rate than non-singletons, we also used 15,718 family-specific heritable insertions, those private to a family yet inherited by at least one offspring, found in Simons Foundation Autism Research Initiative (SFARI) datasets (Fig. S32B). These show the same trend, suggesting that this distribution results from biased insertion, rather than a consequence of selection or technical bias. The opposite insertional bias of these two MEs, which employ the same molecular machinery for insertion (i.e. ORF2p of L1), suggests that other factors, such as recruitment of different RNA-binding protein partners, influence insertional preference. Considering that L1 expression as a prerequisite for SVA transposition, different expression patterns of these RNAs in the context of germline development are unlikely to fully account for this difference (see discussion in Supplementary Note section “SFARI”). As previously reported, L1 and SVA MEVs exhibit the same motif at insertion breakpoints (T/AAAA, Fig. S34), suggesting that the difference of insertion bias is not due to the differences in local sequence recognition by endonuclease.

**Fig. 2.**
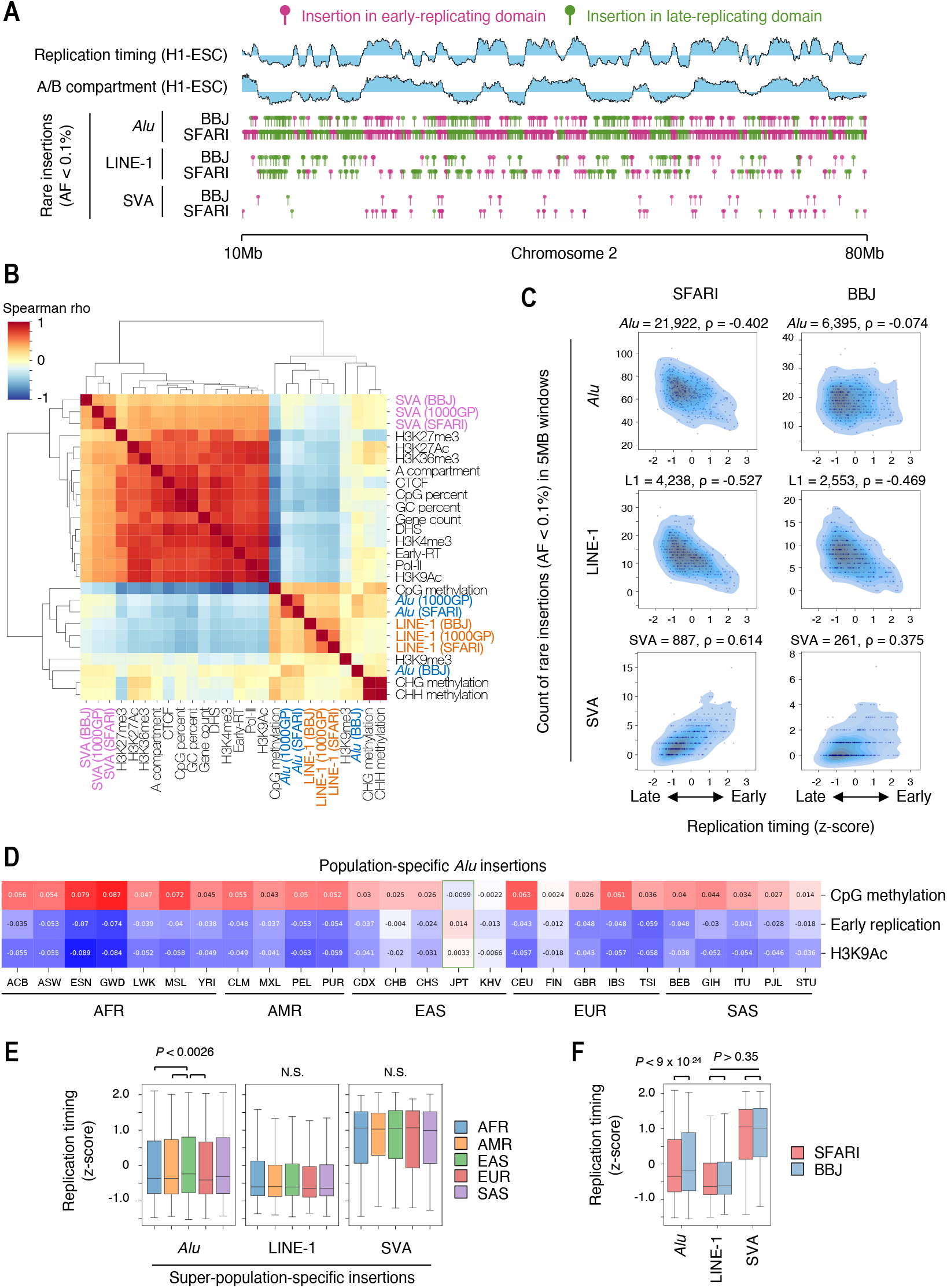
Biased distribution of MEVs. (A) Example of positional distribution of rare MEIs found in BBJ (4,880 subjects) and subjects of PC-inferred European ancestry in SFARI (7,642 subjects). Insertion sites of rare MEIs (AF < 0.1%) in a 70Mb region of chromosome 2 are shown. (B) Heatmap showing correlations between the number of MEIs discovered from the 1000GP, SFARI, and BBJ, and genome features of non-overlapping 1Mb windows. Dendrograms show results of hierarchical clustering. (C) Distribution of replication timing and number of rare MEIs in non-overlapping 5Mb windows. Left three panels show the distributions of MEIs found in subjects of PC-inferred European ancestry (n=7,642) in SFARI, while the right three panels show those of Japanese in BBJ. Kernel density of data points is shown with the actual data points. (D) Heatmap showing correlations between the number of population-specific MEIs discovered from the 1000GP and genome features of non-overlapping 1Mb windows. Japanese in Tokyo (JPT) are highlighted by a green box. (E and F) Distribution of replication timing of the regions in which super-population-specific MEVs are observed in 1000GP (E) or rare MEVs (AF < 0.1%) found in the subjects of PC-inferred European ancestry in SFARI and BBJ (F).

*Alu* insertions from 1000GP and SFARI show weak enrichment in late-replicating domains, whereas this trend is mitigated in BBJ, suggesting that the insertion bias of *Alu* may differ between human populations (Fig. 2B and C). To examine this more closely, we focused on population-specific *Alu* insertions in 1000GP. Compared to other populations’ specific *Alu* insertions, *Alu* found only in JPT show an opposite trend, occurring slightly more often in early-replicating domains (Fig. 2D). This is not a consequence of differences in the chromatin organization of Japanese subject’s genomes, at least as inferred from CpG methylation (Fig. S33C). At the continental population level, *Alu* insertions specific to AFR, AMR, or EUR populations are more biased towards late-replicating domains than those found only in the EAS population (Fig. 2E and S33). Similarly, we observed differences in the replication timing of genome regions bearing rare *Alu* insertions in BBJ subjects compared to PC-inferred Europeans in SFARI (Fig. 2F). We interpret these differences in the distribution of population-specific *Alu* to suggest that *Alu* insertion preference has shifted in East Asians.

### MEVs gene-regulatory effects depend on ME ontology and genomic context

To understand the consequences of MEVs on gene expression, we imputed MEVs in 838 individuals in GTEx and performed eQTL mapping in 49 tissues using both MEVs and SNVs. We defined “ME-eQTLs” as MEVs that are either the lead variants or strongly tagged with the lead SNVs (R^2^ > 0.95). After cross-tissue meta-analysis, we detected 1,073 ME-eQTLs consisting of 778 different MEVs. MEVs were the lead variants of 483 ME-eQTLs in at least one tissue (Fig. 3A). The pattern of the ME-eQTL’s effect size across tissues is similar to that of gene expression (Fig. 3B). More than 60% of detected ME-eQTLs are tissue-specific, and the tissue in which the most tissue-specific and total ME-eQTLs were detected was testis (Fig. S36A).

**Fig. 3.**
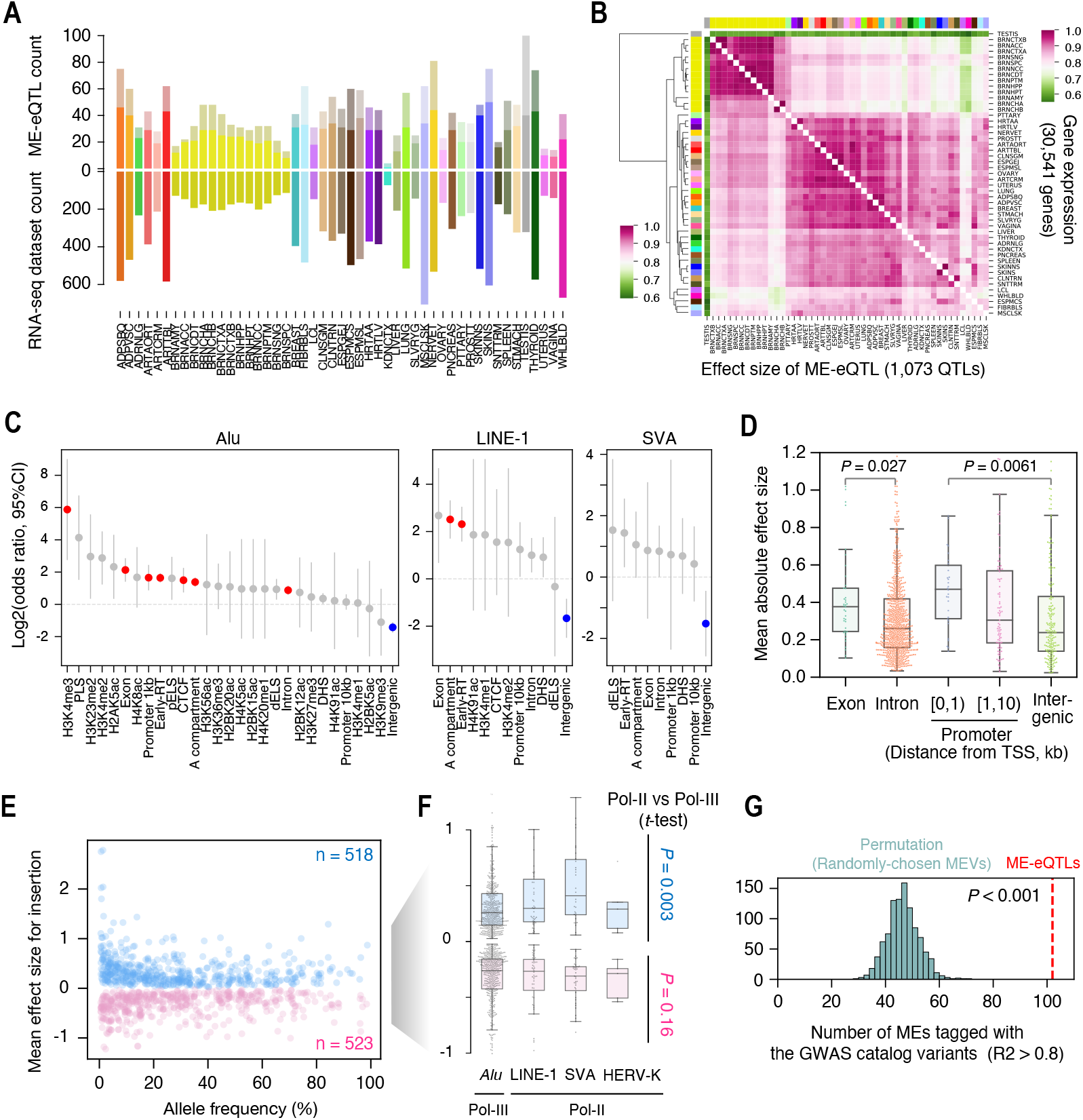
eQTL analysis with MEVs. Number of ME-eQTLs detected in GTEx. Bars with bright and subdued color in the top panel represent the number of multi-tissue and tissue-specific ME-QTLs, respectively. The bottom panel shows the number of RNA-seq datasets used for eQTL analysis. Bar color represents tissue, specified along the horizontal axis. (B) Heatmap showing correlations of gene-expression (upper triangular panel) and effect sizes of ME-eQTLs (lower triangular panel) between tissues. The color of the bars at the top and left indicate tissues as labeled in panel A. Dendrogram shows results of hierarchical clustering. (C) Odds ratios that an ME observed within a designated genome region is detected as an ME-eQTL. Red and blue points are significant enrichments or depletions, having odds ratios significantly different from 1 (*P* < 0.05 after Bonferroni correction). Error bars show 95% confidence intervals. RT, replication timing. (D) Distribution of effect sizes of ME-eQTLs intersecting designated genome features. *P* values of t-test are shown. (E) Distribution of allele frequencies and effect sizes of ME-eQTLs. Effect sizes for presence of an ME are shown, i.e. if a given MEV in a ME-eQTL reflects absence of a reference ME, the sign of the effect size is reversed. (F) Distribution of effect sizes of ME-eQTLs by ME families. (D-F) If a given ME-eQTL is detected in multiple tissues, the mean of the effect sizes across tissues was used for visualization. (E and F) Thirty-two ME-eQTLs that have both positive and negative effects, differing by tissue, were excluded. (G) The number of MEVs in ME-eQTLs in LD with variants in the GWAS catalog. Histogram shows the result of 1000 permutations. Red dot line shows the actual number of MEVs tagged by a GWAS catalog variants. Empirical *P* value is shown.

Gene regulatory effects of MEs plausibly depend on the type of ME and the functional and epigenetic context of the genome, and ME-eQTLs allow us to dissect such determinants. MEVs in regions with active histone marks, such as H3K4me3, and accessible chromatin (represented as early replicating domains and A compartments) are frequently ME-eQTLs. MEVs in exons, promoters (defined as 1kb upstream of TSS), and introns are more often ME-eQTL, while those in intergenic regions are less likely to be detected as ME-eQTLs (Fig. 3C). In concordance with this, ME-eQTLs in exons or promoter regions have larger effects than those in introns or intergenic regions (Fig. 3D). Consistent with the enrichment of genes in early-replicating domains, MEVs in early-replicating domains are more likely to associate with gene expression than those in late-replicating domains. Even when accounting for the increased number of MEV-gene pairs in early-replicating domains, the same trend was observed (*P* < 2.0 x 10^−5^, Fisher exact test). Together this indicates that MEVs in transcriptionally-active regions, regulatory elements, and accessible chromatin often influence gene regulation.

Full-length *Alu* elements contain a Pol-III promoter, while L1, SVA, and human endogenous retrovirus (HERV)-K harbor Pol-II promoters. When comparing the distribution of the effect sizes of *Alu* ME-eQTLs to ME-eQTLs with Pol-II promoter-containing MEs, the latter has larger positive effects, but there was no clear difference when comparing the negative effects, suggesting that MEVs with a Pol-II promoter often function as enhancers of nearby genes (Fig. 3E and F). At the ME family level, SVA is more frequently an ME-eQTL in multiple tissues than *Alu* (Fig. S36B, Fisher exact test, *P* = 0.046), suggesting that SVA has more ubiquitous influence on nearby genes. Thus MEs exert different gene regulatory functions depending on ME family and genomic context.

Compared to the expectation based on permutation, ME-eQTLs are more than twice as often found in high LD with SNVs in the GWAS catalog than expected for non-eQTL MEs variants of the same AF and distance to TSS (Fig. 3G, see methods), suggesting that MEV-associated modulation of gene expression could result in differences in complex traits; thus the integration of ME-eQTLs with GWAS could help refine hypotheses about the molecular mechanisms driving complex traits.

### MEVs often attenuate enhancers

While MEVs with Pol-II promoters often associate with increased expression of nearby genes, some MEVs have negative effects. We hypothesized that ME insertion into an existing gene regulatory element can attenuate that element’s regulatory function, analogous to ME insertion into protein-coding exons generating hypomorphic and loss-of-function alleles. 45 out of 688 MEI-eQTLs fall into distal enhancer-like signatures (dELS) in the ENCODE cCRE dataset. Of these 45 ME-eQTLs, 30 were associated with negative regulation of nearby genes, compared to only 13 with upregulation (Fig. 4A, *P* = 0.007, Fisher exact test), suggesting that ME insertion into an enhancer can decrease its enhancing activity. To test this, we focused on an *Alu* insertion in dELS between genes *DGKE* and *TRIM25* (Fig. 4B). This 297-bp insertion overlaps with a DNase hypersensitive site detected in LCLs and is detected as the lead variant in a *DGKE* eQTL in LCLs; it is in high LD with the lead variant in a *TRIM25* eQTL in LCLs (Fig. 4C and D, R^2^ = 0.98). For both eQTLs, the *Alu* insertion haplotype is associated with decreased gene expression, suggesting that the insertion attenuates enhancer activity. Consistent with this model, the dELS shows enhancer activity in LCLs, while the insertion of *Alu* reduced the reporter activity by half (Fig. 4E). This pattern, of ME-eQTLs in which haplotypes containing *Alu* insertions into dELS are associated with decreased expression of genes presumably regulated by these enhancers, is observed at multiple loci (e.g. Fig S38).

**Fig. 4.**
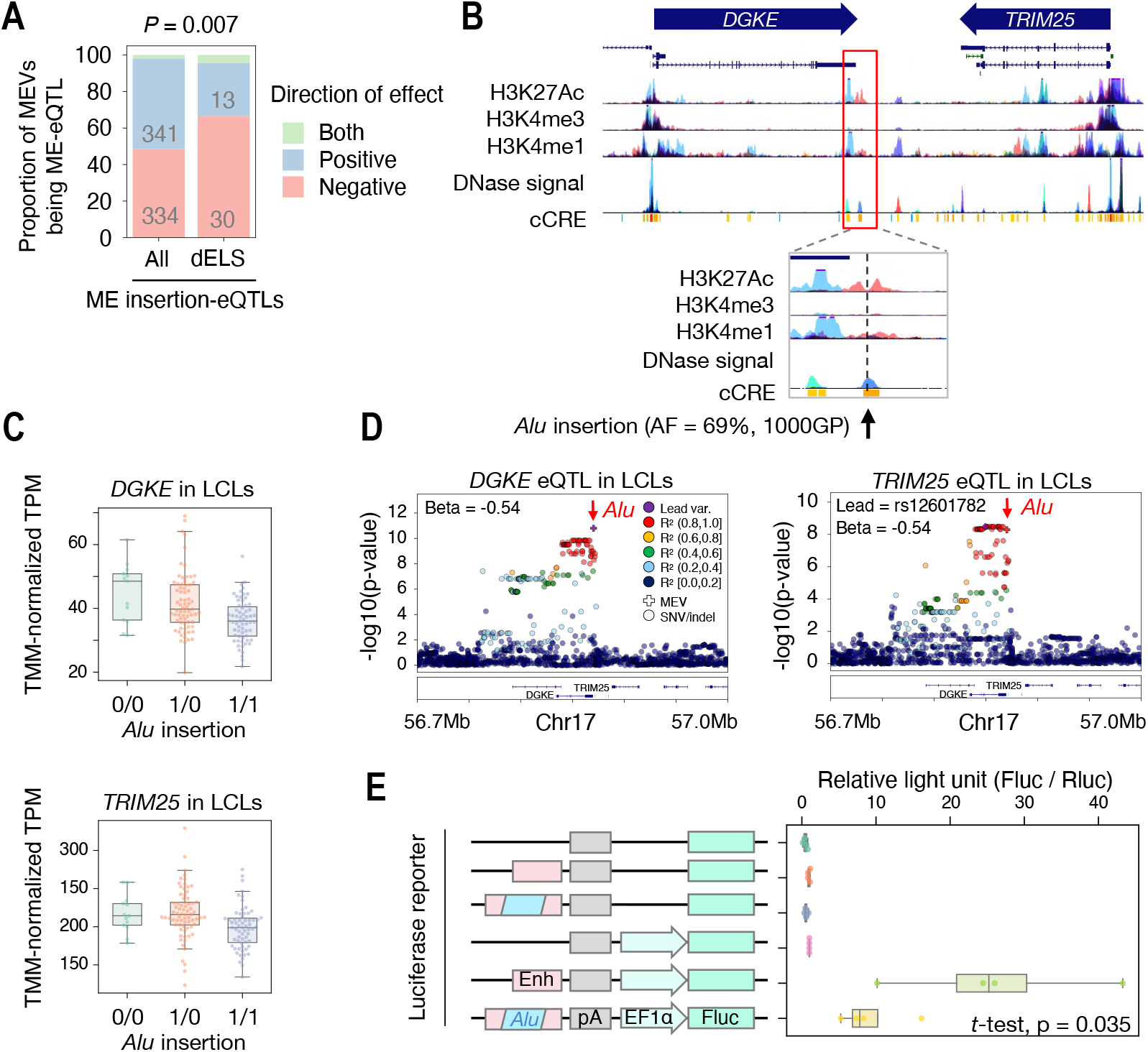
*Alu* insertions in regulatory elements. (A) Comparison between MEVs detected as ME-eQTLs genome-wide and those inserted in distal enhancer-like signature (dELS) candidate cis-regulatory elements (cCRE). Gray numbers in the bar plot show the counts of MEVs used for analysis. P-value of Fisher exact test of whether ME-eQTL variants in dELS more often have negative effect is shown. (B) UCSC genome browser view showing a position of an *Alu* insertion in an enhancer-like sequence near DGKE and TRIM25 genes. The position of the *Alu* insertion is shown with an arrow and a vertical dashed line. (C) Expression levels of DGKE and TRIM25 in LCLs. TMM-normalized TPM grouped by genotypes of *Alu* insertion are shown. (D) Regional association plots showing DGKE-eQTL and TRIM25-eQTL. MEVs and SNPs are shown as plus marks and circles, respectively. The *Alu* insertions are highlighted with red arrows. (E) Dual luciferase reporter assay of the enhancer with or without *Alu* insertion. The illustration shows the structure of Firefly luciferase reporter plasmids. The enhancer and the *Alu* insertion found near *DGKE* and *TRIM25* genes are drawn in red and blue. Plasmids were transfected into a lymphoblastoid cell line (LCL; GM12878). *P* value of t-test between the activities of the bottom two constructs is shown.

### Coherent regulation of gene expression by 3’UTR MEVs

In GTEx, 71 MEVs in 3’UTRs of protein coding genes were used for eQTL mapping. Of these, 20 *Alu* were observed as ME-eQTLs of the genes; 16 were ME-eQTLs in 2 or more tissues. *Alu* in 3’UTR tended to be associated with downregulation of gene expression (Fig. 5A-D). An *Alu* insertion in the 3’UTR of *HSD17B12* was previously reported to downregulate that gene’s expression in iPSCs and LCLs (*9*). This association was replicated in 40 tissues, including LCLs (Fig. 5B-D). To test whether other *Alu* insertions cause differential gene expression, we cloned 3’UTRs of *ADIPOQ* and *MAP3K21* genes (*Alu*-ADIPOQ and *Alu*-MAP3K21, respectively) in a reporter plasmid and generated isogenic controls lacking the *Alu* sequence. The *Alu*-ADIPOQ decreased reporter expression in LCLs, supporting the MEV as causal of the observed association (Fig. 5B-D). Although *Alu*-ADIPOQ was not detected as an ME-eQTL in LCLs, it is detected as an eQTL in all tissues in which *ADIPOQ* is highly expressed. On the other hand, *Alu*-MAP3K21 increased reporter expression in oligodendroglioma cells and basal neuroectoderm-like NT2/D1 cells, but not in LCLs (Fig. 5D-F). This is consistent with the ME-eQTL mapping results; although MAP3K21 is expressed in other tissues, *Alu*-MAP3K21 is an eQTL only in brain tissues. This suggests that factors specific to the brain are required for this particular *Alu* MEV to exert its influence on gene expression. Including singletons, 628 MEVs in the 1000GP datasets were observed in 3’UTRs of protein coding genes. While only 71 were used for eQTL mapping due to low allele frequency in GTEx, which is biased towards European ancestry, these also have the potential to influence gene expression. An East Asian-specific *Alu* insertion in 3’UTR of the pleiotropic gene *EGFR* decreases the expression of the reporter gene (Fig. 5G). While further assessment of the phenotypic consequences of this MEV is warranted, among the 42 diseases tested so far (see below), this variant is modestly associated with asthma (Fig S39, *P* = 0.00018, OR = 1.44).

**Fig. 5.**
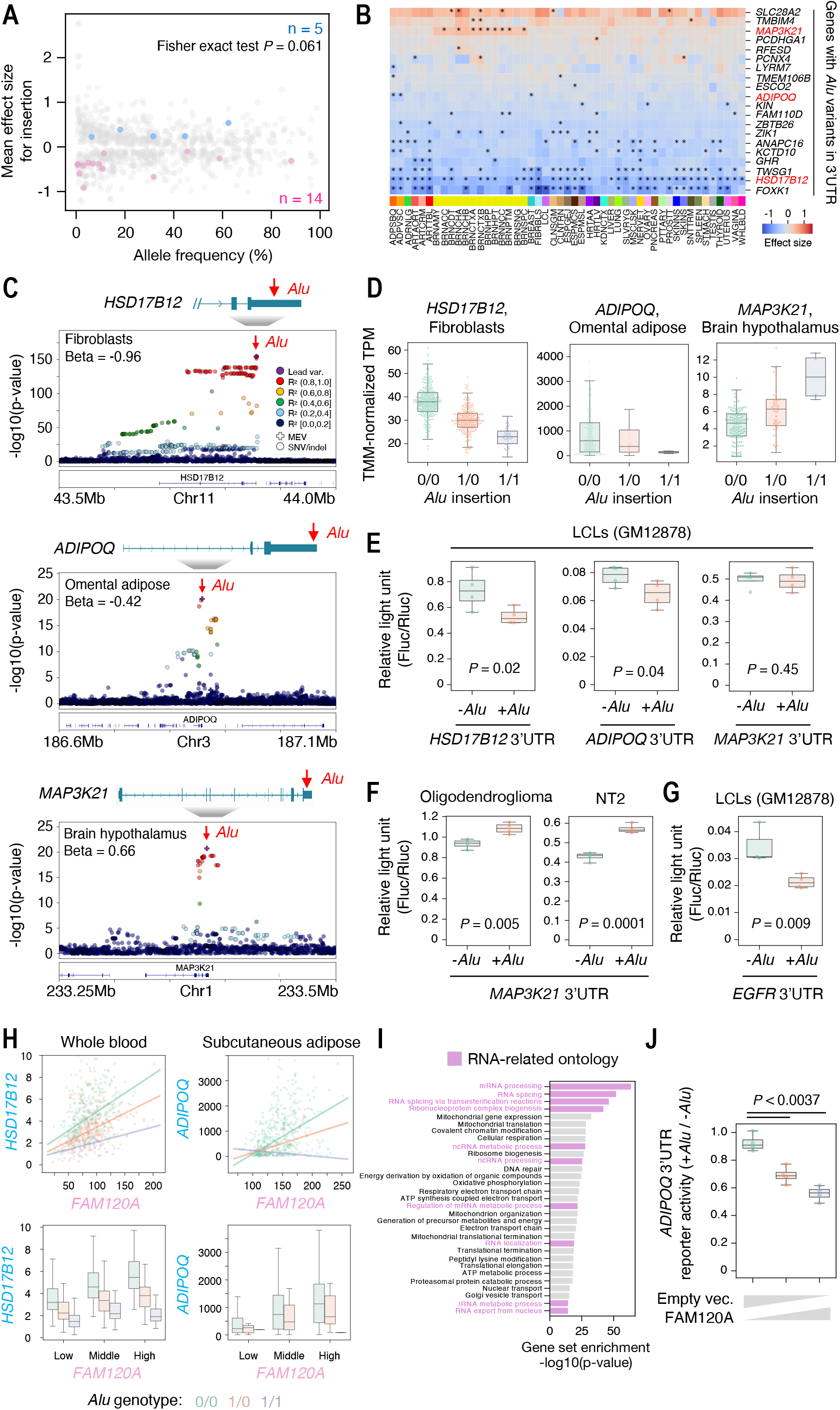
*Alu* insertions in 3’UTRs. (A) Distribution of allele frequencies and effect sizes of ME-eQTLs. Twenty *Alu* insertions in 3’UTRs which are detected as ME-eQTLs of the genes are highlighted with blue and red dots. Effect sizes for presence of ME insertion are shown, i.e. if the MEV in an ME-eQTL represents absence of a reference ME, the sign of the effect size was reversed. (B) Heatmap showing the effects of *Alu* insertions in 3’UTR. Significant associations (LFSR < 0.05) are flagged as asterisks. Color bar at the bottom of the heatmap corresponds to tissue. (C) Reginal association plots showing *HSD17B12* eQTL in fibroblasts (top), *ADIPOQ* eQTL in omental adipose (middle), *MAP3K21* eQTL in hypothalamus. Illustrations on top of plots show the structures of genes and *Alu* insertion sites. MEVs and SNPs are shown as plus marks and circles, respectively. The *Alu* insertions are highlighted with red arrows. (D) Expression levels of *HSD17B12* in fibroblasts (left), *ADIPOQ* in omental adipose (middle), and *MAP3K21* in hypothalamus (right). TMM-normalized TPM grouped by genotypes of the *Alu* insertions are shown. (E) Dual luciferase reporter assays of the *HSD17B12* (left), *ADIPOQ* (middle), and *MAP3K21* (right) 3’UTRs with or without *Alu* insertion. Plasmids were transfected into GM12878. (F) Dual luciferase reporter assays of the *MAP3K21* 3’UTR with or without *Alu* insertion. Plasmids were transfected into Oligodendroglioma (left) and NT2/D1 cells (right). (G) Dual luciferase reporter assays of the *EGFR* 3’UTR with or without *Alu* insertion. Plasmids were transfected into GM12878. (E-G) *P* values of t-test between with- and without-*Alu* insertions are shown. (H) The distributions of expression (TMM-normalized TPM) of eGenes, *HSD17B12* (left) and *ADIPOQ* (right), compared to that of a proxy-gene, *FAM120A*. In the top panels, dot color represents genotype of *Alu* in the two eGenes. Green, orange, and navy lines shows the results of linear regression of the data grouped by *Alu* genotypes 0/0, 0/1, and 1/1, respectively. In the bottom panels, subjects were divided into three groups, low, middle, and high, based on the expression of *FAM120A*. Color of the box plots represents genotype of *Alu* in the two eGenes. (I) Result of gene-set enrichment analysis for proxy-genes. Biological processes with p-value lower than 1e-14 are shown. (J) Ratio of dual luciferase reporter activity of the *ADIPOQ* 3’UTR with *Alu* compared to without *Alu* insertion. The luciferase plasmids were transfected with increasing amounts of a plasmid encoding FAM120A-flag into GM12878.

Our ability to validate the effect of these 3’UTR *Alu* insertions using transient transfection suggests that the mechanism of action does not strictly require the native chromatin context and potential *Alu*-directed chromatin modifications. The *Alu* sequence may recruit factors, such as RNA-binding proteins or nucleases, that stabilize or destabilize the RNA within which it is transcribed. If so, the expression levels of these factors may correlate with the effect of *Alu* on steady-state RNA. In other words, the effect of *Alu* may be dependent on the expression of these other genes, and such genes can be considered as proxies of the *Alu*-eQTL effect (proxy genes). To detect such potential factors, we generated an across-tissue regression model with an interaction term relating *Alu* genotype with proxy gene expression and checked for proxy genes for these 20 ME-eQTLs (see Supplementary Notes). The most often-detected proxy gene was *FAM120A*, which was inferred to be associated with the effect of 11 *Alu* variants (Fig. 5H). The previously-reported *Alu*-binding protein, HNRNPK (*27*), was also detected as a proxy of 4 *Alu* variants. Factors related to RNA degradation, such as *CNOT7* and *EDC3*, and trafficking, such as *XPO7*, are also detected as proxies of more than 6 *Alu* variants. To determine the biological processes enriched among proxy genes, we performed gene set enrichment analysis. Proxy genes are enriched for RNA-related processes, such as mRNA processing and RNA splicing (Fig. 5I), highlighting a list of candidate RNA-binding factors and complexes involved in 3’UTR *Alu*-mediated gene regulation. To validate this approach, we tested the effect of FAM120A overexpression on the regulatory influence of a 3’ UTR *Alu* polymorphism (*Alu*-ADIPOQ) for which it was detected as a proxy. *Alu*-dependent downregulation of reporter gene expression was augmented by the overexpression of FAM120A in a dose-dependent manner (Fig. 5J), consistent with the effect of *Alu*-ADIPOQ being altered by FAM120A.

### Trait association and GWAS including MEVs

To understand the association between MEVs and human traits, we surveyed the LD between MEVs and trait-associated variants identified by GWAS in BBJ and United Kingdom Biobank (Pan-UKB). Out of 4,369 lead variants in 172 GWAS in BBJ, 54 lead variants in 28 GWASs were in high LD with (hereafter, “tagged”) ME polymorphisms (Fig. S40A, R^2^ > 0.8). In Pan-UKB, 833 out of 169,822 lead variants in 7,221 GWASs tagged MEVs; 73 of these lead variants associated with clinically-relevant measurements (Fig. S40B and Table S10).

To demonstrate that MEVs genotyped by MEGAnE can be integrated in GWAS to pinpoint genetic causes of disease risk, we performed GWAS using both SNV/indels and MEVs. ME genotypes were imputed using an imputation reference panel based on 1000GP haplotypes and associated, alongside SNVs, with 42 diseases studied in BBJ. This uncovered 5 MEVs associated with 3 diseases (Fig. 6A-E and S43); one is detected as a lead variant and the four tagged lead variants. Absence of a reference L1 insertion 11-kb upstream of *EVI2A* gene’s TSS (L1-EVI2A, AF = 0.42 in 1000GP) is detected as a new lead variant in GWAS of type 2 diabetes (T2D), replacing the SNV which previously served as the sentinel of this haplotype (Fig. 6B). Whereas this locus has previously been linked to *NF1* as the likely candidate gene (*28*), the L1-EVI2A is also the lead variant of an eQTL of *EVI2A* (encoded from an *NF1* intron) in omental adipose tissue (Fig. 6B). L1-EVI2A also tags a lead SNV in Sex Hormone-Binding Globulin (SHBG) protein GWAS in Pan-UKB (R^2^ = 0.91). The L1-EVI2A haplotype is associated with decreased SHBG, which often inversely correlates with BMI (*29*). Also in T2D GWAS, an *Alu* insertion tagged a lead variant (R^2^ = 0.94) of a locus on chromosome 19 within a cluster of zinc finger proteins (Fig. 6C). This insertion is predominantly found in East Asians; the MAF in JPT and EAS is 2.4% and 4.7%, respectively, while the MAF in other populations is 0.15% or lower. There are only 3 SNVs linked with the lead variant (rs142395395, R^2^ > 0.8), of which one was successfully imputed and used in GWAS. Although non-ME SVs must be considered as well, the possibility that the *Alu* insertion, rather than the SNVs with similar strength of association, is causal can now be weighed. These examples emphasize that integrating MEVs can improve the hypotheses raised by GWAS.

**Fig. 6.**
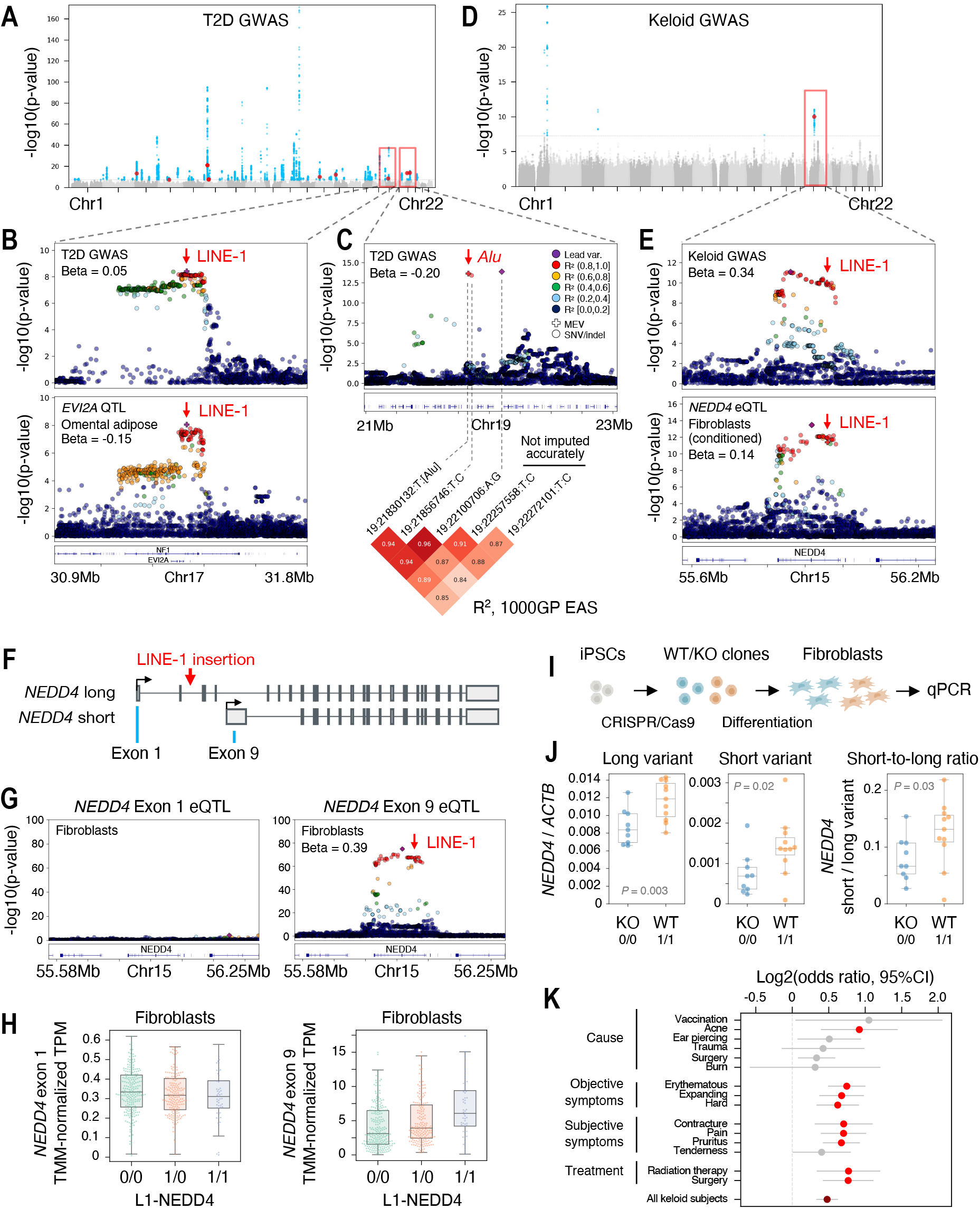
MEVs associate with disease. (A) Manhattan plot of type 2 diabetes (T2D) GWAS in Japanese. (B) Regional association plots showing haplotypes associated with T2D (top) and *EVI2A* expression in omental adipose tissue (bottom). (C) Regional association plots showing haplotypes associated with T2D. The bottom heatmap shows linkage disequilibrium between variants in the LD associated with T2D. Two variants, 19:22257558:C:T and 19:22272101:C:T, are not shown in this plot because these were not accurately imputed (INFO < 0.7). (D) Manhattan plot of keloid GWAS in Japanese. (A, D) Red and blue dots represents MEVs and SNPs, respectively, with *P* value lower than 5e-08. (E) Regional association plots showing LDs associated with keloid (top) and *NEDD4* expression in fibroblasts (bottom). (F) Illustration of the long and short *NEDD4* transcript variants. Location of L1-NEDD4 is depicted with a red arrow. (G) Regional association plots showing no association between variants and exon 1 expression in fibroblasts (left) and association between variants and exon 9 expression in fibroblasts (right). (B, C, E, G) MEVs and SNPs are shown as plus marks and circles, respectively. MEVs are highlighted with red arrows. (H) Expression levels of *NEDD4* exon 1 (left), and exon 9 (right) in fibroblasts. TMM-normalized TPM grouped by genotypes of the L1-NEDD4 are shown. (I) Illustration of the experimental design for L1-NEDD4 knockout (KO) and assessment of NEDD4 gene expression. (J) Expression levels of the *NEDD4* long variant (left), short variant (middle), and short-to-long variant ratio (right). *NEDD4* gene expression was normalized by the expression of *ACTB*. (K) Odds ratios that patients carry L1-NEDD4 based on disease characteristics, including cause of keloid development, signs and symptoms, and keloid treatment history. Red and dark red points show odds ratios significantly above 1 (*P* < 0.05 after Bonferroni correction accounting for additional tests as shown in Fig. S44).

An L1 insertion in an intron of *NEDD4* (L1-NEDD4) associates with keloid and tags the lead SNV, rs16976600 (R^2^ = 0.85, 1000GP EAS) of a chr15 risk locus (Fig. 6D and E). The lead SNV also tags a variant previously associated with keloid severity in the Japanese population, rs8032158 (R^2^ = 0.91, 1000GP EAS) (*30*). Of note, L1-NEDD4 tags lead variants identified in Dupuytren’s disease and fasciitis GWAS in Europeans in Pan-UKB (rs8032158 and rs59912282, R^2^ = 0.71 and 0.86, respectively), suggesting that this haplotype may have a generalizable effect on diseases featuring fibroblast inflammation. L1-NEDD4 is also in high LD with variants associated with increased *NEDD4* expression in fibroblasts. Conditioning by lead variants, we identified three LD blocks that independently reach significance as *NEDD4* eQTLs, one of which is the same as that detected in keloid GWAS (Fig. S42, coloc PP4 = 93%). *NEDD4* has two promoters, expressing long and short transcript variants (Fig. 6F). The short variant is highly expressed in keloid scars and activates inflammatory pathways (*31*). To test whether L1-NEDD4 associates with increased expression of this shorter transcript, we performed exon-eQTL analysis. The expression of exon 9, which is specific to the short variant, is strongly associated with the presence of L1-NEDD4, while exon 1, the long variant-specific exon, is not (Fig. 6G and H). Since L1 often functions as an enhancer, L1-NEDD4 may enhance expression of the short variant and lead to keloid through the previously-proposed activity of this transcript variant on inflammation.

To test the influence of this L1 polymorphism directly, L1-NEDD4 was knocked out in iPSCs derived from a healthy Japanese individual carrying two copies of L1-NEDD4 (Fig. 6I). From a bulk KO cell pool, 9 KO and 11 WT clones were obtained and differentiated into fibroblasts. In cells with biallelic knockout of L1-NEDD4, the expression of *NEDD4* decreased (Fig. 6J). While expression of both variants decreased in KO clones, the effect on the short variant was more pronounced; the ratio of the short variant to the long variant decreased in KO clones. This demonstrates that the L1 insertion functions as an enhancer of *NEDD4*, particularly for the short variant previously implicated in keloid pathogenesis. Because the short variant of *NEDD4* is involved in inflammation (*31*), L1-NEDD4 genotype may explain biological differences underlying heterogeneity in the clinical presentation of keloid. Indeed, L1-NEDD4 increases the odds of developing keloid due to acne, but not after surgery, among BBJ subjects (Fig. 6K and S44, Table S13). L1-NEDD4 also increases the odds of clinical indicators of keloid severity, including contracture and spontaneous pain, as well as history of keloid treatment by radiation or surgical therapy. Thus the molecular pathways activating, and activated by, L1-NEDD4 are rational targets for development of genotype-guided drugs against severe keloid.

## Discussion

Here, enabled by the tool we developed to accurately genotype MEVs from short-read WGS data, we interrogated the consequences of recent ME activity on human genomes and phenotypes. This reveals genetic mechanisms underlying population differentiation, gene expression, and complex traits. Accurate detection of MEVs in diverse human populations allows us to resolve population-specific patterns of recent genome diversification accounted for by ME insertions. For example, L1 and SVA insertions are more abundant in East Asians than in Europeans. These differences may reflect different active copies (*32*) and or differences in the activity or repertoire of factors repressing MEs. While *Alu* insertions tend to be observed in late-replicating domains, this trend was mitigated in East Asians and even reversed in Japanese. This suggests that the insertion preference of *Alu* has shifted as humans have populated the earth. Previous work suggested a similar drift in insertion preference occurred during primate radiation; older, non-polymorphic *Alu* are known to be enriched in early-replicating domains while recent polymorphic ones show the opposite trend (*33*). The factors besides ORF2p that regulate the insertion preferences of human MEs are unknown; changes to the spatiotemporal regulation of transposition-competent ribonucleoproteins could result from accumulation of population-specific mutations in these factors, or in active MEs themselves. A clearer picture of the differences in ME insertion preference, which can be gained by applying the tool we provide to additional genomes from diverse populations, is required to infer the biochemical and phenotypic consequences of differences in ME-derived mutations moving forward.

ME-eQTL analysis provides clues to understand the logic of how MEs influence gene expression depending on ME sequence and genomic context. The *cis*-regulatory function of MEs depends on the genomic context in which they occur, including its chromatin accessibility and epigenetic marks. Insertions in early-replicating regions are more likely to influence gene expression, MEs recruiting Pol-II are more likely to enhance gene expression than *Alu*, and insertions into existing enhancers often attenuate them. These generalizations provide the first comprehensive glimpse at the complex but coherent regulatory logic encoded by MEVs. *Alu* insertions in 3’UTRs are frequently detected as ME-eQTLs and often, but not always, decrease steady state mRNA levels. Although 3’UTR Alus are often detected as multi-tissue eQTL, some are clearly tissue-specific, such as *Alu*-MAP3K21 specific to the brain. It is known that RNA carrying inverted *Alu* repeats, which potentially forms an RNA duplex, is unstable (*34*). HNRNPK, which binds *Alu* RNA, is known to be critical for nuclear retention of RNAs carrying *Alu* (*27*). Thus context (e.g. surrounding sequence and co-expressed genes) is likely critical for licensing *Alu* polymorphisms to exert post-transcriptional regulation. Consistent with this concept, we identified FAM120A as a co-regulator of 3’UTR *Alu*. Disruption of interactions like that of FAM120A could represent a new target for multipurpose precision medicines; the 3’UTR *Alu* MEV in *HSD17B12* causes changes in reporter gene expression and associates with a number of biometric traits and basal metabolic rate (highlighted in Table S10); this variant can thus be considered to cause differences in human weight, and the regulatory contribution of this *Alu* is non-essential. Similarly, a 3’UTR *Alu* in the essential SARS-CoV-2 factor and dementia-linked gene *TMEM106B* (*35*, *36*), detected as an ME-eQTL in several tissues, is associated with a number of neural disease phenotypes (highlighted in Table S10). As highlighted by the East Asian-specific *EGFR* 3’ UTR *Alu*, using WGS to discover and genotype MEVs, as opposed to genotyping only known variants, enables recovery of rare or population-specific MEVs, some of which can be expected to have large effect sizes (*10*).

Inclusion of MEVs in GWAS bridges the gap between known risk loci and previously-overlooked genetic causes, demonstrating a new path to overcome the challenge of connecting GWAS signals to causal variants, especially in non-European populations. We identified five MEs present on known risk haplotypes in Japanese subjects. These include an L1 insertion we show is causal of altered gene expression and therefore suggest mediates the increased keloid risk associated with this haplotype. The observation that a human-specific ME insertion substantially predisposes to keloid, which has not been observed in other primates (*37*), supports the utility of this approach to infer genetic origins of other traits characteristic of our species (*38*). Extending these analyses to larger sets of existing and forthcoming short-read WGS will allow the integration of more MEVs in GWAS of additional phenotypes, leading to the discovery of more disease-causing MEs and motivating development of ME-targeting drugs. By improving detection and prioritization of a class of variants difficult to assess at genome-wide scale, our tool and results are directly applicable to medical genetics (see Supplementary Note “LoF insertions” section). Even so, a major limitation remains: predicting which ME insertions alter phenotype requires additional data integration and statistical testing. However our results also demonstrate that ME ontology relates coherently to MEV effect. Using larger datasets to learn, for example in the framework of natural language processing, the genomic and cellular features that influence the activity of MEs may thus allow us to decipher the semantics of MEs in the genome’s language.

In summary, our analysis leveraged accurate MEV detection and statistical genetics to reveal genetic mechanisms underlying population differentiation and trait diversity. MEVs impact many traits plausibly entangled with fitness in our varied landscapes; here we showed the consequences of several MEVs in disease, providing important information for personalized medicine. Our work provides comprehensive backing to the assertion that MEs are drivers of diversification of genome sequence and function, classic concepts of genome evolution. In addition, we highlight MEs as a source of biased mutation, invoked to account for neutral evolution of complexity (*39*). As the direction and velocity of diversification can be modified by MEs, differences in ME-derived mutation patterns may potentiate differential genome plasticity between lineages.

## Acknowledgement

We are grateful to all of the participants in BBJ, as well as the staff of BBJ for their assistance. BBJ is supported by the Japan Agency for Medical Research and Development (AMED) (Grant Number: JP19km0605001). The Genotype-Tissue Expression (GTEx) Project was supported by the Common Fund of the Office of the Director of the National Institutes of Health, and by NCI, NHGRI, NHLBI, NIDA, NIMH, and NINDS. The GTEx data used for the analyses described in this manuscript were obtained from dbGaP accession number phs000424.v8.p2. We are grateful to all of the families at the participating Simons Simplex Collection (SSC) sites, as well as the principal investigators (A. Beaudet, R. Bernier, J. Constantino, E. Cook, E. Fombonne, D. Geschwind, R. Goin-Kochel, E. Hanson, D. Grice, A. Klin, D. Ledbetter, C. Lord, C. Martin, D. Martin, R. Maxim, J. Miles, O. Ousley, K. Pelphrey, B. Peterson, J. Piggot, C. Saulnier, M. State, W. Stone, J. Sutcliffe, C. Walsh, Z. Warren, E. Wijsman). We appreciate obtaining access to genetic and pedigree data on SFARI Base. The authors wish to acknowledge the resources of the 1,000 Genomes Project and HGDP-CEPH Human Genome Diversity Cell Line Panel. We are grateful to the UK Biobank participants contributing to the results made public via the Pan-UK Biobank Resource and acknowledge the Pan-UKBB team (https://pan.ukbb.broadinstitute.org/team). Super-computing resources were provided by Human Genome Center, the Institute of Medical Science, the University of Tokyo (SHIROKANE), and the Office for Information Systems and Cybersecurity, RIKEN (HOKUSAI General Use project G20021 and Q21537). We acknowledge Dr. Hideyuki Yoshida and Dr. Takahiro Suzuki of RIKEN Center for Integrative Medical Sciences for providing plasmids, Dr. Xun Chen and Dr. Guillaume Bourque of Kyoto University and Dr. Noah Sasa and Dr. Yukinori Okada of Osaka University for testing pre-release versions of MEGAnE, Dr. Kei Sato and Dr. Jumpei Ito of University of Tokyo for helpful discussion, and Ms. Mio Yoshioka for outstanding administrative support. S.Koj., M.N., and J.A.K. acknowledge funding from the Incentive Research Projects of RIKEN, supported in part by Resona Bank. N.F.P. acknowledges funding from Japan Society for the Promotion of Science Grants-in-Aid for Scientific Research(S) KAKENHI 20H05682, Japan Society for the Promotion of Science Grants-in-Aid for Scientific Research(B) KAKENHI 21H02972, RIKEN-McGill International Collaborative grant, Gout and Uric Acid Foundation of Japan, Cluster for Pioneering Research under the Hakubi fellowship program and from the discretionary budget of the Director of the RIKEN Center for Integrative Medical Sciences, Dr. Kazuhiko Yamamoto.

