## Supplementary Note and Figures for "Mobile elements in human population-specific genome and phenotype divergence"

**MEGAnE Development** **5**
Code availability 5
Overview of the algorithm 5
Preparation of MEGAnE *k*-mer set 5
Extract informative reads from WGS BAM/CRAM file 5
Reshape repeat consensus sequences 6
Unmapped read processing 6
Clipped read processing 6
Distantly-mapped read processing 7
PolyA read processing 7
MEI breakpoint search 7
Processing of MEI breakpoint nested in the same ME family 7
Add breakpoint-supporting reads 8
Removal of likely false positive MEI breakpoints 8
ME absence search 9
Genotyping of MEIs 9
Genotyping of ME absences 9
Joint-calling 10
**MEGAnE Benchmarking** **11**
MEV discovery and genotyping accuracy relative to long read sequencing 11
Sensitivity of MEV discovery and genotyping accuracy; comparison with MELT 13
Accuracy of MEV genotyping; comparison with PanGenie and GATK-SV 14
Accuracy of MEV genotyping; Mendelian error 15
Accuracy of MEV genotyping; targeted deep sequencing 15
Imputation accuracy using MEGAnE's callset 16
Computational cost 17
**MEV Discovery and Genotyping using MEGAnE** **19**
Preparation of repeat annotations for MEGAnE 19
1000GP GRCh38 datasets 19
1000GP GRCh37 datasets 19
1000GP GRCh37 datasets, joint calling for EUR 19
BioBank Japan, 25x WGS datasets 19
Joint calling from 25x and 15x WGS datasets in BBJ 20
SFARI 20
Chimpanzee WGS datasets 21
Polymorphism of MEs not reported to be recently active 22
LoF insertions 22
**MEV Discovery and Genotyping using MELT** **24**
**Haplotype Estimation and Genotype Imputation** **25**
Haplotype estimation for MEGAnE callset from 1000GP GRCh38 datasets 25
Haplotype estimation for MEGAnE callset from 1000GP GRCh37 datasets 25

Haplotype estimation for MEGAnE and MELT callset from EUR in 1000GP GRCh37 datasets 25
Genotype imputation for GTEx individuals 26
Genotype imputation in BBJ 26
**Multiplex Targeted Deep Sequencing 27**
Design of PCR primers for targeted sequencing 27
Targeted deep sequencing and mapping 27
Quality control and normalization of DNA-sequencing data 27
Bayesian inference of ME genotypes 28
**Preparation of Genomic Features 30**
Gene count 30
DNA methylation level 30
Replication timing 30
A/B compartment 30
Histone modifications 31
DNase hypersensitive sites 31
TF-binding sites 31
**Analyses Related to MEVs 32**
Detection of motifs at insertion breakpoints 32
PCA of MEVs 32
Intersections between MEVs and gene annotations 32
Correlations between ME insertions and genomic features 32
Preferential insertion of MEVs in the non-transcribed strand of genes 33
Intersections between ME-eQTLs and ENCODE regulatory elements 33
Occurrence of ME-eQTLs by genome features 33
**ME-eQTL Analysis in GTEx 35**
eQTL analysis in 49 tissues 35
Exon expression-QTL analysis in 49 tissues 35
Conditional analysis 35
Across-tissue meta-analysis 36
Detection of ME-eQTL 36
Proxy gene mapping 36
Gene-set enrichment analysis 37
Permutation of MEVs in ME-eQTLs to test linkage disequilibrium with the GWAS catalog variants 37
**Associations Between MEVs and Traits 38**
ME-GWAS of 42 diseases in BBJ 38
Conditional analysis 38
Calculation of LD between MEVs and lead variants in UKB 38
Calculation of LD between MEVs and lead variants in BioBank Japan 38
**Cell-Based Experiments 39**
Cell lines 39
PCR validation of L1-NEDD4 39
Knockout of L1-NEDD4 in iPSCs 39
Differentiation of iPSCs into fibroblasts 39
qRT-PCR of *NEDD4* transcripts 39
Luciferase reporter enhancer assay 40

Luciferase reporter 3'UTR assay 40
Luciferase reporter 3'UTR assay with transient FAM120A expression 40
**Definitions 41**
Box plot 41
**Software Versions 42**
**References 43**
**Supplementary Figure Legend 47**
Fig. S1. Illustration of data processing by MEGAnE 47
Fig. S2. Removal of likely false positives by MEGAnE 47
Fig. S3. Criteria used for genotyping MEIs by MEGAnE 47
Fig. S4. Criteria used for genotyping ME absences by MEGAnE 47
Fig. S5. Comparison of MEV discovery between MEGAnE and PAV using all ME-containing SVs 48
Fig. S6. Comparison of MEV discovery between MEGAnE and PAV using likely TPRT-mediated insertions 48
Fig. S7. Comparison of MEV discovery and genotyping accuracy between MEGAnEand MELT 48
Fig. S8. Comparison of MEV genotyping accuracy between MEGAnE and GATK-SV 49
Fig. S9. Mendelian error rate of MEVs detected by MEGAnE 49
Fig. S10. Processing of targeted deep sequencing data; quality control 49
Fig. S11. Processing of targeted deep sequencing data; target-level normalization and inference of genotypes 49
Fig. S12. Validation of MEGAnE genotyping of 25x depth WGS dataset by targeted deep sequencing 50
Fig. S13. Validation of MEGAnE genotyping result of 15x depth WGS dataset by targeted deep sequencing 50
Fig. S14. Validation of MEGAnE genotyping result of joint calling of 25x and 15x depth WGS datasets by targeted sequencing 50
Fig. S15. Concordance of genotyping between MEGAnE and deep target-sequencing 51
Fig. S16. Comparison of imputation accuracy based on MEV callsets by MEGAnE and MELT 51
Fig. S17. Comparison of imputed MEI genotypes and those determined by targeted deep sequencing 51
Fig. S18. Computational cost of MEGAnE 51
Fig. S19. Number of MEVs discovered from the 1000GP dataset 52
Fig. S20. Comparison of MEV discovery in 1000GP JPT and BBJ 52
Fig. S21. MEV discovery from the SFARI 52
Fig. S22. Thirteen *de novo* L1HS insertions found in SFARI cohort 52
Fig. S23. Eighteen examples of *de novo Alu* insertions found in SFARI cohort 52
Fig. S24. Three *de novo* L1HS insertions shared between two children 52
Fig. S25. Three *de novo Alu* insertions shared between two children 53
Fig. S26. Non-reference MER41A insertion presumably originating from a partial deletion of MER41A in the reference allele 53
Fig. S27. *De novo* THE1_I, a MaLR-like endogenous retrovirus element, insertion found in a trio 53
Fig. S28. Non-reference LTR8A insertion supported by long-reads 53
Fig. S29. Examples of probable novel LoF ME insertions 54
Fig. S30. Principal component analysis using SNVs and MEVs 54
Fig. S31. Super-population- and population-specific MEIs found in the 1000GP dataset 54
Fig. S32. Insertion distribution of singletons and family-specific heritable insertions 54
Fig. S33. Correlation between super-population-specific *Alu* insertion and genome features 54

Fig. S34. Motifs detected at ME insertion breakpoints 55
Fig. S35. Preferential insertion of MEs in non-transcribed strand of genes 55
Fig. S36. Tissue-sharing and distribution of ME-eQTLs 55
Fig. S37. Genome features associated with ME-eQTLs 55
Fig. S38. Three examples of MEIs in gene regulatory elements 55
Fig. S39. Weak association between an *Alu* insertion in the 3'UTR of *EGFR* gene and asthma 55
Fig. S40. Phenograms showing linkage between MEVs associated with traits 56
Fig. S41. PCR validation of L1-NEDD4 allele frequency in Japanese and CRISPR-Cas9 knockout in iPSCs 56
Fig. S42. Conditioning of NEDD4-eQTL and exon-eQTL analysis 56
Fig. S43. T2D and prostate cancer GWAS detect associations with MEVs 56
Fig. S44. Odds ratios of carrying L1-NEDD4 by body part affected by keloid 56

### MEGAnE Development

**Code availability**

MEGAnE is coded in Python 3.7 and C++ 11. Source code is available from GitHub (https://github.com/shohei-kojima/MEGAnE). A complete environment including MEGAnE and other required software is available from Docker Hub (docker://shoheikojima/megane).

**Overview of the algorithm**

MEGAnE finds ME insertions and absences, and genotypes the discovered MEVs. It searches for discordantly mapped reads and finds potential breakpoints from clipped reads. It uses BLASTn to search for similarity between the overhangs of clipped reads and ME insertions. It makes breakpoint pairs that represent the upstream and downstream breakpoints of an ME insertion or absence, or, in most cases, the start and end positions of a target site duplication (TSD). It then extracts breakpoints that are highly likely to derive from ME insertions or absences and fits a gaussian mixture model, which models homozygosity and heterozygosity of the input sample. Based on the modeled distribution, MEGAnE removes likely false positives. After discovering ME insertions and absences, it genotypes the polymorphic MEs based on the number of reads providing evidence of each breakpoint, evidence of breakpoint absence, and read depth of the TSD. It outputs discovered ME insertions and absences in VCF format (Fig. S1).

After MEV discovery and genotyping of multiple samples, MEGAnE can merge them to make a joint callset. It first merges the breakpoint positions in multiple VCF files, then searches for reads providing evidence of the merged breakpoints. If sufficient reads support a breakpoint, discrete genotypes (i.e. '0/1' or '1/1') are assigned. If there are no reads supporting a breakpoint, it genotypes as '0/0'. If there is weak evidence of the breakpoint, it leaves the genotype as missing, i.e. './0'.

**Preparation of MEGAnE *k*-mer set**

MEGAnE first processes the reference genome file. Compressing non-N nucleotide sequences in 2-bit format, it finds all 32-mers that appear more than once (i.e. non-unique 32-mers) in the reference genome. Those 32-mers are used as a reference set of non-unique sequences during the read filtering step before similarity searches between WGS reads and the input genome.

**Extract informative reads from WGS BAM/CRAM file**

MEGAnE takes a BAM or CRAM file as an input and extracts reads that are potentially informative with respect to ME insertions and ME absences, as described below. MEGAnE depends on htslib (*1*) to read the input bam or cram file.

- Chimeric alignment (alignment with the SA:Z tag) mapping to two different genome regions. If the two alignments are on the same chromosome, MEGAnE only extracts such reads if the distance between the two alignments is greater than 20kb. These reads potentially provide evidence of ME insertions.
- Chimeric alignment (alignment with the SA:Z tag) mapping to two different genome regions on the same chromosome, but the distance between the two alignment is shorter than 20kb, which potentially evince absence of a reference ME (i.e. ME absence).

- Reads mapping to distant regions without soft-clipping longer than 20bp (distantly-mapped reads), that is, a read 1 and a read 2 of a paired read are mapping either to the different chromosomes or to the same chromosome, but farther than 2kb.
- PolyA-containing reads, that is a read with soft-clipped region and the proportion of a nucleotide ‘A’ in the soft-clipped region is 70% or more.
- PolyA-containing read, that is a read with soft-clipped region and the proportion of a nucleotide ‘T’ in the soft-clipped region is 70% or more.
- Unmapped reads.

**Reshape repeat consensus sequences**

MEGAnE takes as input a set of consensus repeat sequences (hereafter referred to as consensus MEs) in the RepBase repeat library format. Then it classifies the input consensus MEs into multiple groups based on the pairwise similarity between consensus MEs. It splits the consensus MEs into fragments with the same length as the input WGS read length, and maps the fragments to the consensus ME sequences using BLASTn similarity search (*2*). If there are differently-annotated consensus MEs that show similarity between each other (i.e. any fragments cross-mapping), those MEs will be considered as the same group. This grouping is used when evaluating possible mis-annotation of the ME family of discovered MEIs due to similarities between consensus MEs.

**Unmapped read processing**

MEGAnE maps unmapped reads to the input ME consensus sequences using BLASTn similarity search and finds reads containing any ME sequence. Then, it maps the non-ME parts (i.e. the soft-clipped portion) of such ME-containing reads to the input genome by BLASTn similarity search and finds potential breakpoints of ME-insertions. If a read was mapped to both the input ME consensus sequence(s) and the input genome, i.e. a chimeric read, it annotates the genome position corresponding to the start position of the ME sequence in the chimeric read as an MEI breakpoint. Mapping of sequences to the human genome by BLASTn similarity search is computationally expensive. Particularly, in the case of unmapped read processing, we observed that the non-ME parts of the ME-containing unmapped reads are often heavily multi-mapping to the human genome. To mitigate this computational burden, MEGAnE removes multi-mapping reads before the BLASTn similarity search. It does so by splitting the reads into 32-mers by 2-bit compression and cross-referencing with the MEGAnE *k*-mer set (the set of non-unique 32-mers in the input reference genome) by binary search, removing sequences that potentially multi-map to the input genome.

**Clipped read processing**

MEGAnE maps sequences of the soft-clipped regions of the chimeric reads to the input ME consensus sequences by BLASTn similarity search and finds reads containing ME sequences. If a soft-clipped region maps to the ME consensus sequences, it considers the read as a ME chimeric reads and annotates the genome position corresponding to the start position of the soft-clipping as a potential MEI breakpoint. After the annotation of ME chimeric reads, it processes the non-ME chimeric reads. Even if the soft-clipped regions do not have detectable similarities to the ME consensus sequences, sometimes such reads contain a short ME sequence which is undetectable by the default threshold along with polyA or polyT sequence. To detect such polyA-containing reads, it detects the polyA-containing soft-clipped regions. By default, it keeps reads containing 10 or more ‘A’ or ‘T’ characters in a consecutive 12-nt window. Those polyA-containing reads will be merged with the polyA-reads detected during reading the input alignment.

**Distantly-mapped read processing**

From the distantly-mapped reads, it finds examples in which read 1 maps to ME sequence(s) in the input genome while read 2 maps to a non-ME region in the input genome, or vise versa.

**PolyA read processing**

Some ME insertions have long polyA tails, sometimes longer than the length of the input WGS reads, as a consequence of target-primed reverse transcription (TPRT) acting on poly-adenylated RNA. In such cases, it is difficult to detect chimeric reads mapping to both the input consensus ME sequences and the input genome. To detect breakpoints flanking such long polyA stretches, MEGAnE searches for soft-clipping reads and unmapped reads with polyA and polyT stretches. If the non-polyA or non-polyT region of the polyA read maps to the human genome, it annotates a genome position corresponding to the border of non-polyA and polyA stretch in a WGS read as a potential MEI breakpoint.

**MEI breakpoint search**

MEGAnE gathers the MEI breakpoints found in the unmapped read processing, clipped read processing, and polyA read processing to search for breakpoint pairs. Here, breakpoint pairs represent a pair of upstream and downstream breakpoints of a MEI. Because MEI inserted by TPRT often flank TSDs, the upstream and downstream breakpoints are found at slightly different genome positions, typically 10 to 25-bases away. In the case of MEIs without TSDs (e.g. inserted at short deletions), the positions of upstream and downstream breakpoints will be detected at the same genome position, however, this is a rare case. Therefore, MEGAnE pairs two breakpoints found close to each other (by default, less than 50-bp). It first sorts the positions of the discovered breakpoints, and pairs by the genome sweep algorithm (ref needed?). It pairs breakpoints only when the two breakpoints are thought to be derived from the same ME family. If the result of BLASTn similarity search is available for a breakpoint, it annotates the breakpoint as that of ME(s) detected by BLASTn. If a breakpoint is associated with polyA-containing reads only, it annotates as a polyA-breakpoint; polyA-breakpoints can be paired with ME-defined breakpoints only for MEs that carry polyA tail (in the case of human, *Alu*, L1, and SVA).

We observed that MEs in the input genome are often associated with chimerically-mapped reads. These reads often do not contain non-ME regions, but instead carry two separate sequences coming from the same ME family. During MEGAnE development, we noted that such chimerically-mapped reads result in many MEI breakpoint calls that were not supported in ground truth datasets. We suspect that this phenomenon results from mismapping of reads originating from variations within MEs unrelated to TPRT-mediated mobility, such as a short deletion in an ME, or recombination between two ME copies. To remove such breakpoints, MEGAnE excludes MEI breakpoints nested in the same ME family during breakpoint pairing.

**Processing of MEI breakpoint nested in the same ME family**

As mentioned above, it is difficult to search for true MEIs nested in the same ME family. To recover these, MEGAnE first detects credible MEI breakpoints nested in the same ME family, and then pairs the breakpoints. To detect credible breakpoints, it maps the chimeric reads again to the human reference genome by BLASTn similarity search and finds breakpoints supported by uniquely mappable read(s). As a result of BLASTn search, it searches for uniquely-mapped reads which can be aligned with the human genome with similarity more than 98% only once. The rationale behind this is that if the read contains mutations, typically one or two, that distinguish a specific ME copy from other copies of that ME in the reference genome, those mutations will allow us to map reads to a single ME copy, otherwise we cannot precisely map reads. Searching for such uniquely mappable ME-derived reads is a computationally expensive step, as MEs are abundant in the input genome. To reduce the computational burden, MEGAnE searches for potentially multi-mapping reads by cross-referencing with non-unique 32-mers before BLASTn similarity search using the same method as described in the "Unmapped read processing" paragraph.

**Add breakpoint-supporting reads**

After pairing breakpoints, MEGAnE assigns ME reads detected in the distantly-mapped read processing to the paired breakpoints. If a distantly-mapped read maps to both a consensus ME sequence belonging to the same ME family as the ME of a breakpoint and to the genome region proximal to the MEI breakpoint (by default, within 500-bp) it assigns the distantly-mapped read as a breakpoint-supporting read.

**Removal of likely false positive MEI breakpoints**

After the pairing of breakpoints, MEGAnE removes likely false positives by evaluating the number of breakpoint-supporting reads, i.e. chimeric reads defining the breakpoints and distantly-mapped reads. MEGAnE fits a statistical model to the distribution of the number of breakpoint-supporting reads and determines the cutoff threshold to remove likely false MEIs. It first counts the number of breakpoint-supporting reads for each breakpoint. In the case of MEIs in both the maternal and paternal allele (i.e. homozygous MEIs), the number of supporting reads will be approximately twice as high as the MEIs that are present in either the maternal or paternal allele (i.e. heterozygous MEIs). When there are enough MEIs and the depth of WGS are high enough, those two types of MEIs will generate two Gaussian distributions (when the depth of WGS is not high, it would be the mixture of two Poisson or negative binomial distributions) (Fig. S2A). Therefore, it first extracts breakpoints that are highly-likely to arise from MEIs, which are supported by distantly-mapped reads mapping at both the upstream and downstream of a breakpoint pair, and fits a gaussian mixture model to the number of breakpoint-supporting reads. In more detail, it models heterozygosity and mean, variance, and height of the gaussian distribution coming from heterozygous MEIs. The likelihood of being heterozygous and homozygous can be described by the mean count of breakpoint-supporting reads $\mu$ and the variance of breakpoint-supporting reads $\sigma$:

Heterozygous MEI: $p(y\ |AC = 1, \mu, \sigma) = Normal\_pdf(y\ |\mu, \sigma))$

Homozygous MEI: $p(y\ |AC = 2, \mu, \sigma) = Normal\_pdf(y\ |2\mu, \sqrt{2\sigma^2}))$

The ratio of the two distributions is determined by heterozygosity, h. The likelihood of the model can be described as:

$$p(y \mid \mu, \sigma, h) = h \times Normal\_pdf(y\ |\mu, \sigma)) + (1 - h) \times Normal\_pdf(y\ |2\mu, \sqrt{2\sigma^2}))$$

MEGAnE fits this model to the observed data, the distribution of the breakpoint-supporting reads of MEIs, by the maximum likelihood method. Then, it rejects the breakpoints that have a -1.96 or lower z-score of a gaussian distribution corresponding to the heterozygous MEIs. Application of this filtering to the >30x 2,504 WGS from the 1000GP shows the linear correlation between the WGS read depth and the determined filtering threshold of breakpoint-supporting reads (Fig. S2B), showing that an appropriate filtering threshold can be determined by this method based on the character (i.e. WGS depth and heterogeneity) of an input dataset.

During development, we observed relatively frequent false positive L1 insertions at consecutive A or T stretch in the genome presumably due to the presence of polyT stretch in the L1 consensus sequence. To exclude such false positives, MEGAnE removes such L1 insertions found in polyA or polyT-rich regions defined as L1 insertion candidates with a TSD longer than 5-bp and high A/T percentage (the total A or T content of the TSD exceeds 50%, or total content of A plus T exceeds 90%).

**ME absence search**

To find absent MEs, MEGAnE uses chimeric alignments mapping to two different genome regions on the same chromosome for which the distance between these two alignments is shorter than 20kb. It first finds breakpoint pairs from the chimeric alignments. It then judges whether the region between the upstream and downstream breakpoints is a ME in the input reference genome.

**Genotyping of MEIs**

To genotype MEIs, MEGAnE uses three criteria:

- Number of breakpoint-supporting reads.
- Read depth of TSD.
- Presence of breakpoint-spanning reads, that is, reads that span the breakpoints presumably coming from the reference allele (without an MEI).

As mentioned in the section "Removal of likely false positive MEI breakpoints," the distribution of the numbers of breakpoint-supporting reads are composed of two distributions representing homozygous and heterozygous MEIs. MEGAnE fits a Gaussian mixture model to the distribution and uses this distribution as the first estimation of zygosity. MEIs often flank TSDs. TSDs are sequence duplications, thus the WGS read depth at a TSD should be 1.5 times (in the case of a heterozygous MEI) or 2 times (in the case of a homozygous MEI) as high as the read depth of the surrounding genome region (Fig. S3A). In rare cases, there are short deletions at breakpoints. In this case, the WGS read depth at a short deletion should be half (in the case of heterozygous MEI) or zero depth (in the case of homozygous MEI) compared to the surrounding genome region (Fig. S3B). By using the numbers of breakpoint-support reads and the read depths of TSDs or short deletions, it estimates the allele counts of MEIs (Fig. S3C-F). In addition to the evidence coming from the presence of MEIs, MEGAnE also uses the evidence originating from the absence of MEIs. If a MEI is heterozygous, there may be reads that are mapped over the insertion breakpoint (breakpoint-spanning reads). Presence of such breakpoint-spanning reads is one criterion supporting heterozygous insertion. MEGAnE gathers the three criteria (i.e. number of breakpoint-supporting reads, read depth at TSD or short-deletion, and breakpoint-spanning reads) and judges whether each insertion is more likely homozygous or heterozygous. If the genotype of a given MEI is concordant across all three evidence criteria, it gives the filter "PASS" flag to the variant, otherwise, it is marked with another flag. Finally, it outputs genotypes in the VCF format.

**Genotyping of ME absences**

To genotype absent MEs, MEGAnE uses similar methods as for genotyping of MEIs. It uses the two criteria below:

- Number of breakpoint-supporting reads.
- Presence of breakpoint-spanning reads, that is, reads that span the breakpoints which presumably come from the allele with the ME.

Similar to the genotyping MEIs, it fits the Gaussian mixture model and estimates the distribution resulting from homozygous and heterozygous ME absence (Fig. S4A). If an absent MEV is heterozygous, there may be spanning reads that span the borders of the ME presence of such spanning reads is one criterion supporting heterozygous absence. MEGAnE gathers the evidence and estimates whether each absence is more likely homozygous or heterozygous (Fig. S4B and C). If the genotype predicted using both criteria is concordant, it gives the filter "PASS" flag to the variant, otherwise, the variant is marked with another flag. Finally, it outputs genotypes as the VCF format.

**Joint-calling**

After the discovery and genotyping of multiple samples, MEGAnE merges the MEV discovery and genotyping results and outputs a joint-callset as VCF format. it first merges the breakpoint positions in the multiple VCF files, then searches for reads providing the evidence of the merged breakpoints. The positions of detected breakpoints are sometimes slightly different, often 1 or 2-nt differences probably due to single nucleotide variations near breakpoints, so by default MEGAnE merges MEIs from multiple samples that have TSD or short deletion overlapping each other. If there are enough reads supporting a breakpoint (i.e. the variant was detected in the original VCF file), it genotypes as '0/1' or '1/1'. If there are no reads supporting breakpoint, it genotypes as '0/0'. If there is weak evidence of the breakpoint (i.e. the variant was not detected in the original VCF file, but reads supporting the breakpoint can be found), it leaves the genotype as missing, i.e. './0'. It also adds the filter "PASS" flag when the variant was flagged as "PASS" in the original VCF files; specifically more than 50% of genotypes of a variant across samples are flagged as “PASS,” such variant will be flagged as “PASS,” otherwise, non-“PASS.”

### MEGAnE Benchmarking

**MEV discovery and genotyping accuracy relative to long read sequencing**

We compared MEGAnE's callset to SVs identified in 34 individuals using long read sequencing and haplotyping techniques (*3*). We used the SV callset generated by the phased assembly variant caller (PAV) (http://ftp.1000genomes.ebi.ac.uk/vol1/ftp/data_collections/HGSVC2/release/v2.0/integrated_callset/variants_freeze4_sv_insdel.tsv.gz). The PAV data contains 35 individuals, however, we excluded one individual (NA24385) from this analysis because 30x short-read data from this individual was unavailable in the 1000GP GRCh38 dataset. Referencing the "RMSK" column of the variant file, we considered variants annotated as either "DNA", "LINE", "LTR", "Retroposon", "SINE", or "RC" as polymorphic MEs. The PAV callset contained 14,691 autosomal MEIs and 9,831 autosomal ME absences. We applied MEGAnE to the 30x WGS data (1000GP data mapped to GRCh38) of the same 34 individuals and generated a joint callset. MEGAnE detected 10,797 MEIs and 2,295 ME absences. Of these, 7,599 and 2,131 variants passed the stringent quality filter (i.e. filter “PASS” variants). We counted the number of ME polymorphisms overlapping between PAV and MEGAnE callsets. In the case of MEIs, if an insertion is within 50-bp of an MEI insertion of the same ME family in the other dataset, we consider that the variants are the same. In the case of ME absence, if more than 90% of the length of the SV annotation overlaps between the two callsets, we consider that the variant is the same. Using these parameters, 8,383 MEIs and 2,039 ME absences are called by both MEGAnE and PAV (Fig. S5A and B). In other words, 57% and 21% of SVs annotated as insertions or deletions involving MEs, respectively, identified by long reads were also detected by MEGAnE. Of these, 7,367 MEIs and 1,997 ME absences passed the filter for high-quality variants. When considering variants that pass MEGAnE’s quality filter, 97% of MEGAnE MEIs are true positives (false detection rate is 3%) and 94% of MEGAnE ME absence calls are true positives (false detection rate is 6%) (Fig. S5A). When including the filter non-"PASS" variants, 78% of MEGAnE MEIs are true positives (false detection rate is 22%) and 89% of ME absence calls are true positives (false detection rate is 11%) (Fig. S5A and B). Considering these false positive rates, we decided to use variants with the filter "PASS" flag for most subsequent analyses.

Although the SVs identified by PAV above overlap with some repeat annotation, not all of them were generated through the TPRT mechanism that characterizes ME mobilization, the SV mechanism we designed MEGAnE to study. For example, 6,261 of the 9,831 ME absences detected by PAV (64%) are thought to be partial deletions of MEs or deletions overlapping MEs in the reference genome rather than TPRM-mediated insertion; 3,221 are absences completely nested in a reference ME of the same family ME that is at least 100-bp longer than the overlapping absence (i.e. likely a partial deletion of a reference ME rather than a pre-integration allele), and 3,040 absences involve more than 100-bp of an ME but also overlap more than 100-bp of non-ME sequence (i.e. likely a deletion overlapping a reference ME and flanking regions, thus unlikely to reflect a pre-integration allele, though the possibility that these reflect absence of an ME with at least a 100-bp sequence transduction cannot be excluded). Concordant with this, the distribution of lengths of absent MEs detected by MEGAnE produces peaks of *Alu* and L1, which are approximately 300 bp and 6 kb, while the distribution of lengths of absence calls by PAV also detects variants with various sizes, particularly shorter ones, suggesting that PAV absence callset contains non-TPRT-mediated partial absences of MEs. Similarly, not all of the 14,691 PAV-annotated insertion SVs that contain some ME sequences reflect TPRT-mediated ME insertion. To evaluate the proportion of TPRT-mediated ME insertions and absences that MEGAnE can detect, we filtered the PAV insertions for those with ME-homologous sequences with target-site duplications and polyA tails, identifying 8,362 and 2,210 PAV insertions and deletions with clear evidence of TPRT-mediated mobilization, respectively (Table S2). To search target-site duplications, we used a weighted Smith-Waterman algorithm allowing total mutation cost up to 2. We set an indel cost as 2, a transition cost as 1, and a transversion cost as 2. Alignment length for target-site duplication search was limited between 6 and 25bp. The sequences starting from the upstream and downstream ME insertion breakpoints were used to search for target-site duplications. PolyA was searched from the sequences adjacent from TSD found above by allowing 1nt mutation. If the length of polyA stretch exceeded 8, we defined it as polyA tail. The distribution of TSD lengths was very similar to those detected by MEGAnE (Fig. S6A and B). Thus we also assessed MEGAnE performance after filtering out these likely non-TPRT-mediated SVs. Of the 2,210 PAV deletions, MEGAnE detected 1,769 (80.0%) and 67 (3.1%) as filter "PASS" and non-"PASS" variants, respectively, totaling 83.1% recovery (Fig. S6C and D). Of the 8,362 PAV insertions, MEGAnE detected 6,829 (81.7%) and 608 (7.3%) as filter "PASS" and non-"PASS" variants, respectively, for a total recovery of 89% (Fig. S6C and D). These figures likely more accurately reflect the sensitivity of MEGAnE to detect TPRT-mediated ME mobilizations.

One design goal of MEGAnE was to improve detection of ME insertions nested in MEs in the reference genome. To evaluate the sensitivity to detect such polymorphisms, we looked into the ME polymorphisms called by long-reads that are nested in other MEs and simple repeats. We masked the reference human genome, GRCh38DH, using RepeatMasker with the human repeat library from Dfam v3.2. Out of 14,691 autosomal MEIs identified by PAV, 5,255, 4,407, and 198 variants were nested in an ME of the same ME class, an ME of a different ME class, and simple repeats, respectively. Of these MEIs, MEGAnE detected 1,067, 3,536, and 66 MEIs, respectively (Fig. S5C). In other words, MEGAnE detected 20%, 80% and 33% of MEIs that are nested in an ME of the same ME class, an ME of a different ME class, and simple repeats, respectively. Of the suspected TPRT-mediated MEs, MEGAnE detected 76%, 88%, and 77% respectively.

There are 4,831 MEIs identified by PAV whose insertion did not intersect with MEs in the reference genome or simple repeats. MEGAnE identified 3,714 of these; conversely, it missed 1,117 (Fig. S5C, blue circle). These 1,117 MEIs consist of 423 L1s, 420 *Alu*, 139 ERVs, 69 DNA transposons, and 66 retroposons (Fig. S5D). Out of 420 *Alu*, 235 were *Alu*Y, which are frequently polymorphic in humans, suggesting that those are recent ME insertions missed by MEGAnE. The 423 L1s are composed of 97 L1MC1, 61 L1HS, and others. L1MC1 is an old mobile element and to our knowledge no evidence of recent activity has been reported. These polymorphisms likely reflect deletions of partial L1MC1; in other words, GRCh38 has an allele with partial deletion of L1MC1, while pre-deletion alleles also exist in humans. L1HS is a recently active mobile element, thus these likely represent 61 L1HS inserted by transposition but missed by MEGAnE. The similar trend is the same for ERVs. There are 139 ERV polymorphisms missed by MEGAnE, however, those included only 7 LTR5_Hs, which is recently active, suggesting that most of the ERV polymorphisms missed by MEGAnE are SVs generated by non-transpositional mechanisms, e.g. deletions within or overlapping older ME insertions.

To evaluate the genotyping accuracy of MEGAnE, we compared the genotypes of polymorphic MEs between PAV and MEGAnE callsets from 34 individuals. We used the same criteria to find variants detected by both PAV and MEGAnE as described above. The majority of MEGAnE genotypes are concordant with PAV (Fig. 1A). Considering all MEGAnE MEIs, including the ones with filter non-"PASS," 80% and 88% of the variants have $R^2$ more than 0.98 and 0.8, respectively. Considering only variants with filter "PASS" flag, 85% and 94% of the variants have $R^2$ more than 0.98 and 0.8, respectively. When using all MEGAnE absent ME calls, including the ones with filter non-"PASS," 64% and 85% of the variants have $R^2$ more than 0.98 and 0.8, respectively. Limiting to variants with filter "PASS" flag, 65% and 85% of the variants have $R^2$ more than 0.98 and 0.8, respectively. This demonstrates that the majority of MEGAnE calls are genotyped accurately. In the case of MEIs, more than 80% are genotyped accurately, using as ground truth the genotype based on genomes assembled and phased using long reads and template strand-specific sequencing.

To validate whether MEGAnE detects ME insertion breakpoints precisely, we compared the breakpoints found in the 34 individuals with those identified by PAV (Fig. S5E). 90% of the MEIs with the filter "PASS" flag share exactly the same breakpoint, demonstrating that MEGAnE accurately determines the breakpoint for most insertions.

**Sensitivity of MEV discovery and genotyping accuracy; comparison with MELT**

We compared the output of MELT, a tool often used in population-scale ME genotyping (*4*, *5*), and MEGAnE by applying both tools to 503 European WGS. We downloaded the fastq files used for the 1000GP 30x GRCh38 dataset and mapped the reads on GRCh37 provided from the 1000GP (human_g1k_v37.fasta). The alignment was position-sorted and used as input for MEGAnE and MELT. We compared variants with filter "PASS" flags. In the case of MEIs, if two insertions from each callset fall within a 20-bp distance, we consider that the variant is the same. In the case of ME absence, if more than 90% length of two annotations from each callset overlap with each other, we consider that the variant is the same. In addition to meeting these position and length criteria, we only considered MEs to represent the same variant if they were also of the same family (e.g. SINE1/7SL, L1, SVA). MEGAnE detected 13,112 variants consisting of 10,922 MEIs and 2,190 ME absences. MELT detected 12,984 variants consisting of 9,560 MEIs and 3,515 ME absences (Fig. S7A). We first checked whether those variants are also detected in 35 individuals sequenced by long-reads (*3*). Out of 13,112 and 12,984 variants called by MEGAnE and MELT, respectively, 6,510 and 4,036 MEs were detected by PAV. This suggests that the MEGAnE callset has a lower false positive rate and a higher true positive rate than MELT. Considering MEI, MEGAnE and MELT found 4,632 and 3,719 MEIs that are also detected by PAV (Fig. S7B). In the case of ME absence, MEGAnE and MELT detected 1,878 and 317 absences also called in the PAV dataset, respectively (Fig. S7B). There is substantial overlap between MEGAnE and MELT MEI callsets, however, the ME absence calls have limited overlap (Fig. S7A). Thus while MELT calls more ME absences than MEGAnE, these are very often false positives when compared to ground truth based on long reads. In contrast, MEGAnE can call both MEIs and absences that intersect well with variants called using long reads, although the sensitivity in detecting ME absence from short-read data is lower, as expected.

To assess genotyping accuracy, we compared the ME genotypes by cross-referencing with the PanGenie callset (Fig. S7B-D). Out of 13,112 and 12,984 variants called by MEGAnE and MELT, 5,158 (3,872 MEIs and 1,286 ME absences) and 3,347 (3,151 MEIs and 196 ME absences) variants were also genotyped by PanGenie, respectively. In the case of MEIs, both MEGAnE and MELT have high concordance with PanGenie; 79% and 71% of MEIs from

MEGAnE and MELT, respectively, has very high concordance with the PanGenie callset ($R^2$ > 0.98). In the case of MEGAnE MEI calls, 94% have $R^2$ more than 0.9 with the PanGenie genotypes. In the case of ME absences, 70% of MEGAnE calls have very high concordance with the Pangenie callset ($R^2$ > 0.98) while only 3% of MELT calls have high concordance. In the case of MEGAnE ME absence calls, 90% have $R^2$ more than 0.9 with the PanGenie genotypes.

This demonstrates that both MEGAnE MEI and ME absence calls have high concordance with a graph-based genotyper of SVs detected by long reads, whereas MELT is less sensitive for detecting and less accurate in genotyping MEs, particularly the absence of reference MEs.

**Accuracy of MEV genotyping; comparison with PanGenie and GATK-SV**

The comparison in the previous section only used 34 individuals. To evaluate the genotyping accuracy of MEGAnE on the joint callset of a much larger number of individuals, we compared between PanGenie and MEGAnE callsets from 3,202 individuals. PanGenie is a short-read SV genotyper using graph-based pangenome references. We used the stringent set of PanGenie genotypes for PAV variants (http://ftp.1000genomes.ebi.ac.uk/vol1/ftp/data_collections/HGSVC2/release/v1.0/PanGenie_filters/20201009_HHU_genotyping_pangenie-filters-freeze3_scored-sv_README.txt) as ground truth. There are 8,750 MEIs and 3,398 absence calls in the PanGenie callset. Of these, 6,919 MEIs and 1,399 absent MEs were shared with our MEGAnE callset. The majority of the MEGAnE genotypes are concordant with PanGenie. When using all MEGAnE MEIs, 74% and 87% of the variants have $R^2$ more than 0.95 and 0.8, respectively. When only considering variants with a filter "PASS" flag (6,110 variants), 80% and 93% of the variants have $R^2$ more than 0.95 and 0.8, respectively (Fig. S8A and C). When using all MEGAnE absent ME calls, 60% and 86% of the variants have $R^2$ more than 0.95 and 0.8, respectively. Considering only filter “PASS” variants (1,362 variants), 61% and 88% of the variants have $R^2$ more than 0.95 and 0.8, respectively (Fig. S8A and C).

Ebert *et al*. (*3*) provide the GATK-SV callset from the same 3,202 individuals (http://ftp.1000genomes.ebi.ac.uk/vol1/ftp/data_collections/1000G_2504_high_coverage/working/20210124.SV_Illumina_Integration/1KGP_3202.Illumina_ensemble_callset.freeze_V1.vcf.gz). GATK-SV is a pipeline wrapping multiple short-read SV callers to generate an ensemble callset. We compared the PanGenie and GATK-SV callsets (Fig. S8B and C). Out of 8,750 MEIs that were genotyped by PanGenie, 6,486 MEIs are detected in GATK-SV (MEGAnE detected 6,110 MEIs with the filter "PASS" flag). We calculated the $R^2$ between PanGenie and GATK-SV callset; 22% and 74% of the variants were more than 0.95 and 0.8, respectively. Out of 3,398 ME absences that were genotyped by PanGenie, 2,440 MEIs are detected in GATK-SV (MEGAnE detected 1,362 ME absence variants with the filter "PASS" flag). We calculated the $R^2$ between PanGenie and GATK-SV callsets; 13% and 75% of the variants were more than 0.95 and 0.8, respectively. Thus MEGAnE and GATK-SV call a similar number of MEIs, yet more than 80% of MEGAnE filter "PASS" calls exceed $R^2$ 0.95, a cutoff only 22% of GATK-SV MEI calls exceed. In summary MEGAnE genotypes MEs better than the ensemble output of current short-read SV callers. In the case of ME absence, while detection by MEGAnE is not as sensitive as GATK-SV, genotyping by MEGAnE outperforms GATK-SV (61% and 22% of MEGAnE and GATK-SV variants shows $R^2$ higher than 0.95 with the PanGenie callset). Thus when comparing the number of accurately-genotyped ME absences, MEGAnE shows better performance than GATK-SV (826 and 307, respectively). Although MEGAnE improves the genotyping accuracy of absence MEs, these variants remain more less accurately genotyped than MEIs.

**Accuracy of MEV genotyping; Mendelian error**

To further corroborate the genotyping accuracy of MEGAnE, we evaluated the Mendelian error rate in trios. We used 602 trios sequenced at 30x in the 1000GP. We applied MEGAnE to the 3,202 individuals in 1000GP and generated a joint callset. For this analysis, we only used MEIs with the filter "PASS" flag. The average Mendelian error rate in a trio is 1.75% (Fig. S9A). More than half of the Mendelian errors reflect the child having an insertion not observed in either parent. This suggests that the Mendelian error mainly derives from genotyping error in the parents, false positive detection in children, or *de novo* insertion. Rare insertions, including singletons, may be especially difficult to detect. To evaluate the Mendelian error rate of rare MEIs, we focused on family-specific MEIs. We looked into variants that are observed in only one family in the 3,202 individuals, regardless of allele count. Of those variants detected in a child, we checked whether the variant was also detected in the parents. As a result, 95% of family-specific variants detected in children were not errors (Fig. S9B). The remaining 5% were Mendelian error; the insertion was not detected in parents. This demonstrates that the MEGAnE callset has relatively low detection and genotyping error rate even if the variant was not commonly observed in humans.

**Accuracy of MEV genotyping; targeted deep sequencing**

To evaluate the genotyping accuracy of MEGAnE by an orthogonal method, we performed targeted deep sequencing of polymorphic ME loci. We selected 160 ME polymorphisms found in Japanese subjects and designed PCR primers expected to amplify the polymorphic loci in such a way that the NGS reads should span the breakpoint of the mobile element insertions. We amplified the 160 target sites in 2221 individuals and sequenced the PCR amplicons by NGS. For a combination of reasons including technical error and PCR failure, 50 of the primer pairs/ME polymorphisms were non-informative. By Bayesian inference, we were able to genotype 110 ME polymorphisms from the NGS read counts of target amplicons (see “Multiplex Targeted Deep Sequencing” section).

The individuals used for target sequencing were selected based on having previously been sequenced at either 25x or 15x depth in Biobank Japan (BBJ) 1st and 2nd cohorts (*6*, *7*). We applied MEGAnE to those individuals and generated joint callsets. We generated three joint callsets from 1) only 25x, 2) only 15x, and 3) both 25x and 15x depth WGS. In all joint calling patterns, MEGAnE genotyping agrees well with deep sequencing.

We compared the genotyping results between deep sequencing and MEGAnE applied to 25x WGS (Fig. S12). All targets were genotyped in at least 861 individuals. When using all MEIs, 89% of targets had Pearson $r > 0.8$ (97 out of 109 targets). When it comes to MEIs with the filter flag "PASS," 95% of targets have Pearson $r > 0.8$ (86 out of 91 targets) (Fig. S12A). As expected based on the similar genotyping results, the allele frequency of ME polymorphisms were also largely concordant with each other (Fig. S12B). When considering all genotyped sites in individuals passing quality control (see “Multiplex Targeted Deep Sequencing” section), more than 95% of MEGAnE genotypes were concordant with the deep sequencing result (Fig. S15). As expected, variants with the filter flag non-"PASS" were less accurate; about 87% of the genotyped sites showed concordance. The same trend is true for MEGAnE results of 15x WGS and both 25x and 15x depth WGS (Fig. S13 and S14). Correlation between MEGAnE and the deep sequencing was consistently high (most targets had r > 0.8) regardless of the allele frequency or ME family (Fig. S12B and C, S13B and C, and S14B and C). In concordance with the low Mendelian error for rare variants, it is notable that MEGAnE has good genotyping accuracy for very rare variants, including accurately genotyping a variant detected in only one of the individuals used for targeted sequencing. The few targets with discrepancies between MEGAnE and deep sequencing suggest MEGAnE may sometimes mis-genotype a homozygous insertion as heterozygous. Inferring the commonalities of the variants in which this occurs and designing an approach to properly genotype such variants will be an area for improvement of future releases of MEGAnE.

**Imputation accuracy using MEGAnE's callset**

We performed haplotype phasing by merging SNVs and polymorphic MEs called and genotyped by MEGAnE and MELT. We applied MEGAnE and MELT to 503 EUR individuals in 1000GP and generated joint callsets (described in "Sensitivity of MEV discovery; comparison with MELT" section). For downstream analysis, we used MEGAnE variants with the filter "PASS" flag. In the case of MELT, we used all variants. We merged the ME variants with the SNV callset from 1000GP and removed variants with a minor allele count of 1. We removed SNVs that violate Hardy-Weinburg equilibrium (HWE), however we did not remove ME variants that violate HWE in order to evaluate imputation quality for all the ME variants in the final output from MEGAnE and MELT. To generate imputed ME genotypes, we split the 503 individuals into 5 groups consisting of either 100 or 101 individuals, and performed leave-one-group-out haplotype estimation and genotype imputation. The haplotypes were estimated by SHAPEIT4 software (*8*) and converted to the imputation reference panel by Minimac3 software (*9*). To simulate the situation of genotype imputation from SNV array data, we only used the SNVs present on the Affimetrix Axiom array as target haplotypes. For example, we estimated the haplotypes of 403 individuals and generated an imputation reference panel. Then, imputation of ME genotypes in the remaining 100 individuals were performed. By doing this for all five groups, we generated the imputed ME genotypes for all 503 individuals. We compared the imputed genotypes and the PanGenie callset (Fig. S16). Using the ME callsets from MEGAnE and MELT respectively, we were able to impute 4,557 (3,300 MEIs and 1,257 ME absences) and 2,908 (2,729 MEIs and 179 ME absences) ME polymorphisms that are also present in the PanGenie callset. In the case of MEGAnE, 56% of MEIs (n = 1,869) and 59% of ME absences (n = 737) had very high concordance with the genotypes from PanGenie ($R^2 > 0.98$) (Fig. S16A). The majority of MEs (90% of MEIs and 91% ME absences) imputed using MEGAnE to generate the imputation reference panel have high concordance with PanGenie callset ($R^2 > 0.9$). On the other hand, 49% of MEIs (n = 1,335) and 3% of ME absences (n = 6) imputed using MELT to generate the reference panel had very high concordance with the genotypes from PanGenie ($R^2 > 0.98$) (Fig. S16A). Notably, the majority of ME absence imputed using MELT to generate the imputation reference panel were not concordant with PanGenie, while MEGAnE performed as well for absences as for MEIs. Similarly, for MELT-imputed genotypes, the $R^2$ drops as the allele frequency increases (Fig. S16 B and C). In summary, MEGAnE was able to generate a larger number of ME genotypes that can be used for haplotype estimation and genotype imputation, compared to a leading existing ME discovery and genotyping tool.

To validate the imputation accuracy by an orthogonal method, we compared imputed genotypes with genotypes determined by deep targeted sequencing. We applied the imputation reference panel generated from the 2504 individuals in 1000GP GRCh37 datasets, which includes 104 Japanese individuals, to the 1,549 Japanese genotyped by Illumina

OminExpressExome SNV array in BBJ (*6*, *7*). Out of 90 PCR targets imputed by this imputation reference panel, 67 (74%) have Pearson r with larger than 0.8 between the genotypes determined by the targeted sequencing and imputation (Fig. 1B and S17A). The majority of the MEs that are imputed with high Minimac imputation R2 ($R^2 > 0.7$) had high concordance between the genotypes determined by the targeted sequencing and imputation. As expected, the imputation accuracy is relatively high; for most, Pearson r is higher than 0.8, while the relatively rare ones are less accurately imputed (Fig. S17B and C).

Overall, MEGAnE allows the imputation of MEVs more accurately than the current standard tool used for detection and genotyping of MEVs from short-read data. While common MEVs will also be readily imputable using new tools based on phased assemblies and the haplotypes in 1000GP, MEGAnE will enable accurate imputation of increasingly-rare MEVs, scaling with the availability of short-read WGS data.

**Computational cost**

To demonstrate the typical computational requirement for MEGAnE, we ran MEGAnE on a supercomputer, SHIROKANE. SHIROKANE provides a computational environment for biobank-scale genome analysis, and is composed of a Intel Xeon Gold 6154 (3.0 GHz) processor, a Lustre parallel distributed file system, and Altair Grid Engine. In summary, MEGAnE can analyze the 2,504 30x WGS CRAM files in the 1000GP within 2 days using 200 threads; thus it can be applied to biobank-scale WGS to efficiently generate a joint callset using massive job parallelization possible with supercomputers. Each step of this process is detailed below.

*Step 0 - generation of MEGAnE k-mer set*

In step 0, MEGAnE generates a *k*-mer set file. We ran this step with the human genome reference, GRCh38DH. This took 13 min and 49.3GB of memory (Fig. S18B).

*Step 1 - calling and genotyping of polymorphic MEs*

First, we applied MEGAnE to 35x WGS (NA12878 mapped on GRCh38 as CRAM format, 1000GP dataset) (Fig. S18A). We used MEGAnE on a Singularity container available from docker://shoheikojima/megane. The CPU time required for this analysis was about 2 hours. When one thread was allocated, the wall time was 1 hour and 56 minutes and the maximum memory usage was 5GB. The wall time was shortened to less than 1 hour when 4 or more threads were allocated. The maximum memory usage was less than 3 GB times the number of threads when the multi-threading was used. Because some steps in MEGAnE cannot be forked to multi-threads, the wall time does not shorten linearly; we recommend setting 2 to 4 threads per run to balance wall time with computational resources.

Software run across a large number of parallel jobs sometimes behave differently than when run with one sample due to technical limitations, such as file access speed (i.e. I/O). To evaluate whether MEGAnE can be used in a large number of parallel jobs, we applied MEGAnE in the Singularity container to 2,504 30x WGS in 1000GP with 100 jobs parallelized (Fig. S18B and C). We allocated 2 threads and 15GB memory per job. The average wall time to analyze one sample was 1 hour 43 minutes. Analysis of NA12878 with 100 jobs parallelized took 1 hour and 36 minutes. This is 30% longer than analyzing only one file at once. This difference would be attributed to the slower file access speed to the same files (e.g. human genome reference file, repeat annotation file, BLAST index, etc) from the 100 parallelized jobs. The longest wall time required to analyze one sample was 5 hours 9 minutes (HG03611). This sample was sequencing the most deeply (74x) among the 2,504 samples. Most of the jobs required less than 6GB memory, however, 12 samples required more than 7GB memory, and 2 samples required more than 10GB but less than 12GB (HG03829 and HG03885). Those two samples contain the third highest and highest number of discordantly mapped reads across 2,504 samples, respectively. In summary, MEGAnE can analyze a 30x human WGS within 2 hours and realistic memory usage under massive parallelization in a supercomputer.

*Step 3 - joint calling*

The joint calling step of MEGAnE consists of two jobs; joint calling of MEIs and joint calling of ME absences. Joint calling of MEI of the 2,504 samples took 2 hours and 3 minutes and 8.9GB memory. Joint calling of ME absences took 30 minutes and 5.5GB memory (Fig. S18A).

### MEV Discovery and Genotyping using MEGAnE

**Preparation of repeat annotations for MEGAnE**

We masked the reference human genomes, GRCh38DH, hs37d5, and human_g1k_v37, by RepeatMasker version 4.1.0 with a human repeat library from RepBase, version 24.01.

**1000GP GRCh38 datasets**

The 30x WGS data from 3,202 individuals mapping to GRCh38DH were downloaded from the 1000GP website (http://ftp.1000genomes.ebi.ac.uk/vol1/ftp/data_collections/1000G_2504_high_coverage/). Throughout this paper, we refer to this dataset as "1000GP GRCh38 datasets." MEs were discovered and genotyped using MEGAnE's call_genotype_38 command. The joint callset was generated using MEGAnE's joint_calling_hs command. We also generated a joint callset from 2,503 individuals, which does not include relatives, using from the same dataset. We generated a separate joint callset for 34 individuals who were sequenced using PacBio in the 1000GP HGSVC project. The HGSVC sequenced 35 individuals by PacBio, however, we excluded one individual, HG002, from our joint callset, since the individual was not included in the 3,202 individuals who were sequenced in the 1000GP 30x WGS. In 2,503 individuals analyzed here, MEGAnE detected 48,248 MEVs with the filter “PASS” flag. Of those, 8,609 (18% of total) were common variants (AF > 1%).

**1000GP GRCh37 datasets**

The raw fastq reads of the 2,504 individuals in the 1000GP 30x GRCh38 datasets were downloaded from the 1000GP website (http://ftp.1000genomes.ebi.ac.uk/vol1/ftp/data_collections/1000G_2504_high_coverage/). The fastq reads were mapped on the human reference genome build, human_g1k_v37 by BWA MEM using the same options as used by 1000GP to map on GRCh38DH (http://ftp.1000genomes.ebi.ac.uk/vol1/ftp/data_collections/1000G_2504_high_coverage/20190405_NYGC_b38_pipeline_description.pdf). In brief, we used the -Y option with the -K 100000000 option. Throughout this paper, we refer to this dataset as "1000GP GRCh37 datasets." The output alignment was converted to CRAM format and analyzed using MEGAnE's call_genotype_37 command. The joint callset was generated using MEGAnE's joint_calling_hs command. In 2,504 individuals analyzed here, MEGAnE detected 48,360 MEVs with the filter “PASS” flag. Of those, 8,665 (18% of total) were common variants (AF > 1%) (Fig. 1E).

**1000GP GRCh37 datasets, joint calling for EUR**

We also generated a joint callset from 503 individuals within European (EUR) populations to compare the detection accuracy and genotyping of MEVs between MEGAnE and MELT. For this, we used the results from the MEGAnE's call_genotype_37 command described in the section above. We generated the joint callset from the 503 individuals using MEGAnE's joint_calling_hs command.

**BioBank Japan, 25x WGS datasets**

We applied MEGAnE to the 25x WGS (either 160 or 150-bp paired-end) from 1,235 individuals in BBJ (*6*, *7*). We mapped the raw fastq reads to the human reference genome hs37d5 by BWA MEM using the same option as we used for mapping of 1000GP dataset, and saved as CRAM format. We did not perform further individual-level QC since the dataset was already subjected to the QC. The output CRAM files were analyzed by the MEGAnE's call_genotype_37 command. The joint callset was generated by the MEGAnE's joint_calling_hs command. In 1,235 Japanese individuals analyzed here, it detected 10,996 MEVs with the filter "PASS" flag. Of those, 4,943 (45% of total) were common variants (AF > 1%). This callset was used for evaluating linkage disequilibrium between MEVs and SNVs.

**Joint calling from 25x and 15x WGS datasets in BBJ**

To find rare insertions in Japanese subjects, we generated a joint callset by merging as many Japanese individuals as possible. To this end, we analyzed additional 30x WGS (either 125 or 124-nt paired-end) from 256 individuals and 15x WGS (150-bp paired-end) from 3,389 individuals by MEGAnE and merged with the MEVs detected from 1,235 individuals described above. When analyzing 15x WGS, we used the `-lowdep` option of MEGAnE, which assumes non-Gaussian distributions of supporting read counts in heterozygous and homozygous insertions. In total, we merged MEVs from 4,480 Japanese individuals using the joint_calling_hs command. In 4,880 Japanese individuals analyzed here, MEGAnE detected 24,933 MEVs with the filter "PASS" flag. Of those, 5,452 (22% of total) were common variants (AF > 1%) (Fig. 1E). This joint callset was used to investigate ME insertion preferences in Japanese.

**SFARI**

We applied MEGAnE to the 30x WGS from 9,201 individuals in SFARI (SFARI BASE request ID# 10109.2.2). The CRAM files were analyzed using MEGAnE's call_genotype_38 command. The joint callset was generated using MEGAnE's joint_calling_hs command. After joint calling, we checked the number of detected MEIs and removed two individuals who had very low counts of MEIs. We also removed 12 individuals with atypical read depth of sex chromosomes and one individual with an atypical chromosome 21 read depth (consistent with trisomy). This resulted in a joint callset consisting of 9,186 individuals. In these 9,186 individuals, we detected 52,401 MEVs with the filter "PASS" flag. Of those, 6,376 (12% of total) were common variants (AF > 1%) (Fig. S21A).

To investigate the insertion distribution of MEs in subjects of European ancestry, we filtered subjects based on analysis of principal components. The first 2 principal components were calculated from MEVs in 9,186 individuals (see "PCA of MEVs" section). The distribution of PC1 and 2 was very similar to that observed in PCA of MEVs discovered from 26 populations in 1000GP (Fig. 1D). We defined 7,642 individuals whose PC1 and 2 fall into the region corresponding to a cluster of EUR subjects in the 1000GP (red rectangle in Fig. S21B) as ones with PC-inferred European ancestry.

To find *de novo* insertions, we first searched for singletons that are observed in a child but not in the other individuals, including both father and mother. To generate a stringent credible set, we next checked the presence of chimeric reads and polyA- and polyT-containing reads mapping to the ME insertion breakpoint in parents. If at least one parent has such reads at the ME insertion site, we excluded such candidates from the analysis. As a result, we found 119 *Alu*, 15 L1 and 3 SVA *de novo* insertion candidates from 2,390 families with 4,311 children. To validate if our *de novo* ME insertion set is credible, we visualized the insertion sites of 20 randomly selected *Alu* and all 15 L1 and 3 SVA insertions using IGV. By manually and visually checking the breakpoint-supporting reads in children and parents, we validated the majority of *de novo Alu* and L1 are true positives (Fig. S22 and S23). Out of 20 randomly selected *Alu de novo* insertions, 18 were confirmed to be likely *de novo* by checking of breakpoint-supporting reads, while 2 were insertions inherited from one of the parents, showing that around 90% of *Alu de novo* calls are likely true positives. Of 15 L1 insertions, 13 were found to be likely true. All the 3 SVA were found to be likely present, in at least a detectable fraction of sequenced cells, in one of the parents; detection of *de novo* SVA and differentiation from mosaic insertions is a challenge MEGAnE is unsuited to overcome at this time.

To find *de novo* insertions that are shared between two children, we searched for doubletons that are observed in two children in a family, but not present in either father and mother (Table S6). We excluded twins from this analysis. We next checked the presence of chimeric reads and polyA- and polyT-containing reads mapping to the ME insertion breakpoint in parents. If at least one parent has such reads at the ME insertion site, we excluded such candidates from the analysis. Finally, we visualized the insertion sites of all members of the families with such insertions by IGV genome browser and manually validated the presence of insertions in children and absence in parents (Fig. S24 and S25). As a result, we found 3 L1HS and 3 *Alu de novo* doubletons that are shared between two children in a quad but not present in either parent's sequenced blood cells. Such mosaic *de novo* insertions can arise through early zygotic mutation in the parent, such as an insertion during 2-cell to blastocyst differentiation, and unequal contribution of the mosaic ME variant to germline and somatic tissues. Such mosaicism can also arise through ME insertion in the early stages of spermatogenesis and oogenesis, such as in primordial germ cells (PGCs) which differentiate into sperm cells or oocytes. If two children are derived from the same germline stem cells carrying a *de novo* ME insertion, both children should carry the ME insertion, which cannot be found in the blood of parents. Based on the observation of 6 mosaic insertions in X quads, it is estimated that at least 0.6% of individuals carry insertions for which one of their parents exhibits mosaicism, if we consider that mosaic insertions will be passed onto offspring with 50% probability. This argues that the majority of heritable ME insertions occur during spermatogenesis or oogenesis, while few are inserted at the early stages of germline or the early zygotic development (*10*–*12*).

#### Chimpanzee WGS datasets

To find MEVs that are shared between humans and chimpanzees, 39 chimpanzee WGS data (PRJNA635393 (*13*)) were downloaded and mapped to a chimpanzee genome build, panTro5 by BWA-MEM (*14*) with a `-K 100000000` option. To analyze chimpanzees by MEGAnE, we first annotated MEs and repeat sequences in the panTro5 genome build by RepeatMasker (*15*). Because the purpose of this analysis is to identify the MEVs that are shared between humans and chimpanzees, we used the same human repeat library (RepBase version24.01) as we used to mask the human genome. As a result, 7,230,205 MEs were annotated in panTro5, which is close to the number of those annotated in GRCh38 (7,341,504). Using the human repeat library and the ME annotations in the chimpanzee genome, we applied MEGAnE to the 39 chimpanzee WGS. As a result of the joint calling, we detected 9,395 and 2,071 MEIs and ME absences, respectively, including 6,281 and 1,953 MEIs and ME absences with the filter-"PASS" flag (Table S3 and 4). To cross-reference between chimpanzee and human MEVs, the genome positions of the chimpanzee MEVs were lifted over to the human genome, build GRCh38. In the case of MEIs, we lifted over the positions of breakpoints, i.e. start and end positions of TSD. We observed that some non-reference MEIs in the chimpanzee genome are observed as MEs in the human reference genome. Because of this, we accepted cases in which liftover of the start and end positions of TSDs to two positions in the human genome fall within 10kb, but removed if the distance was larger than 10kb. In the case of ME absences, we lifted over the positions of 50bp upstream of 5'-end and 50bp downstream of 3'-end of the absent sequences. Similar to the situation of MEIs, some MEs in the reference chimpanzee genome are not found in the human genome, and TSD flanked with such MEs, which are two copies in the chimpanzee genome, exist as a single copy in the human genome. Because of this, the exact positions of the 5'-end and/or 3'-end of ME absences frequently failed to be lifted over to the human genome. To solve this, we lifted over the positions of 50bp upstream of 5'-end and 50bp downstream of 3'-end of the absent sequences, based on the fact that the length of the majority of TSD is less than 25bp. After the liftover, we added and subtracted 50bp from the two lifted-ever positions (i.e. the start and end of ME absences), respectively. As a result 8,555 and 1,903 chimpanzee MEIs and ME absences, respectively, were lifted over.

**Polymorphism of MEs not reported to be recently active**

Out of 45,317 and 2,931 MEIs and ME absences with the filter "PASS" flag identified from the 1000GP GRCh38 datasets, 1,562 and 294, respectively, were not variants involving recently active MEs (*Alu*Y, L1HS, SVA, LTR5_Hs). There are several possible explanations why presumably long-inactive MEs are still polymorphic. One common reason is partial deletions within old MEs (Fig. S26). A polymorphic partial deletion of an old ME in the reference genome (e.g. GRCh38DH) may be detected as an insertion compared to the reference. Another possibility is templated-sequence insertion (TSI) (*16*). One *de novo* insertion found in SFARI is a partial insertion of an endogenous retrovirus THE1_I, which is not reported to be active (Fig. S27). This *de novo* insertion has signatures consistent with TSI, including representing a partial ME insertion, presence of TSD, and perfect identity to a distant genome region. Another possibility is nonallelic gene conversion between MEs (*17*). Nonallelic gene conversion between old and young polymorphic *Alu* copies may generate polymorphisms of old *Alu*. It is also possible that polymorphisms generated by transposition in the distant past are retained by balancing selection (*18*). Analyzing 39 chimpanzee genomes, we found 15 polymorphisms shared between human and chimpanzees. Five, including L1PA4 which is relatively old L1, were found in the HLA locus, which is known to harbor variant haplotypes maintained by balancing selection (Table S5). Lastly, it is also possible that excision of an insertion by recombination between TSDs re-generates the pre-integration site (*19*). For example, we found a non-reference insertion of LTR8A copy which is present in macaque and marmoset genomes (Fig. S28). This shows that the insertion occurred before the divergence of human and marmoset, and it is likely that the polymorphism is generated by deletion of the whole LTR8 copy, presumably by recombination between TSDs.

**LoF insertions**

Some MEIs are found in CDS regions, and these are likely to generate loss-of-function (LoF) variants. We cross-referenced the MEI positions with a gene annotation downloaded from GenCode (v26 for MEIs found in the 1000GP and v26lift37 for those found in BBJ). In MEIs identified from 2,503 individuals in the 1000GP GRCh38 dataset, 81 MEIs flagged with filter

"PASS" were found in CDS regions. Of these, 22 were found in two or more individuals. In MEIs flagged with filter "PASS" found in the 4,880 Japanese individuals in BBJ, 51 were found in CDS, and 17 were not singletons. Some are previously-reported disease-causing mutations. For example, we detected an *Alu* insertion in the CDS of *RP1* gene in 26 Japanese in BBJ, which causes retinal degeneration when present in two copies (*20*). We also found an L1 insertion in the CDS of *CC2D2A* gene, which was previously reported as a recessive mutation causing occipital encephalocele, post-axial polydactyly, and multicystic renal disease (*21*). This L1 insertion was reported in a single individual, however, we found this insertion in 4 Japanese subjects, demonstrating that this L1 insertion is not private to the previously-studied kindred. We also discovered previously unreported insertions. For example, an *Alu* insertion in CDS of *ADGRE2* gene was found in two Africans in the 1000GP. A missense substitution in *ADGRE2* is a known risk for vibratory urticaria, suggesting that the *Alu* insertion may also be a risk for this condition (Fig. S29A and B). In 4 Japanese subjects, we identified a L1 insertion in CDS of *DLEC1* gene, a tumor suppressor gene often mutated in a variety of cancers (*22*), for which further assessment of cancer-specific risk is warranted (Fig. S29C and D).

#### MEV Discovery and Genotyping using MELT

We applied MELT version 2.1.5 (*4*) to the 503 individuals in 1000GP who belong to EUR. As input files, we used alignments mapping to the human reference genome build, human_g1k_v37 (see "1000GP GRCh37 datasets" section). We discovered and genotyped *Alu*, L1, and SVA polymorphisms according to the instructions available in the ReadMe file. In brief, we used MELT's Preprocess, IndivAnalysis, and Deletion-Genotype commands to discover and genotype ME polymorphisms from each BAM file. Then, we used the GroupAnalysis, Genotype, MakeVCF, and Deletion-Merge commands to make joint callsets.

### Haplotype Estimation and Genotype Imputation

**Haplotype estimation for MEGAnE callset from 1000GP GRCh38 datasets**

First, we merged the MEI and ME absence callsets from MEGAnE. We used MEGAnE's reshape_vcf command to merge these two callsets and remove multi-alleic ME variants. To estimate haplotypes of 2,503 individuals in 1000GP phase3, we merged the ME callset with SNVs. For quality control (QC), we first split the ME callset into individuals belonging to each of 5 super-populations and evaluated Hardy-Weinburg equilibrium. Variants which violated Hardy-Weinburg equilibrium ($P < 1 \times 10^{-6}$) in at least one super-population were removed. SNVs that overlap with polymorphic MEs were removed. Singleton SNVs and MEs were also removed. Then, the QC-ed ME callset was merged with the SNV callset (1000GP, GRCh38_v1a) without variants violating Hardy-Weinburg equilibrium ($P < 1 \times 10^{-6}$). Each chromosome of the merged callset was saved in VCF format and phased using SHAPEIT4 software with default genetic maps. The phased haplotypes were converted to an imputation reference panel using Minimac3 software. Due to the unavailability of SNVs on sex chromosomes, we estimated the haplotypes for MEs only on autosomes and PARs.

**Haplotype estimation for MEGAnE callset from 1000GP GRCh37 datasets**

First, we merged the MEI and ME absence callsets from MEGAnE using the same MEGAnE command described in the previous section. To estimate haplotypes of 2,504 individuals in 1000GP phase3, we merged the ME callset with SNVs and indels. We first split the ME callset into individuals belonging to each of 5 super-populations and evaluated Hardy-Weinburg equilibrium. Variants which violate Hardy-Weinburg equilibrium ($P < 1 \times 10^{-6}$) in at least one super-population were removed. SNVs and indels that overlap with polymorphic MEs were removed. Singleton SNVs, indels, and MEs were also removed. Then, the QC-ed ME callset was merged with the SNV and indel callset (1000GP, v5a) without variants violating Hardy-Weinburg equilibrium ($P < 1 \times 10^{-6}$). Each chromosome of the merged callset was saved in VCF format and phased by SHAPEIT4 software with default genetic maps. An imputation reference panel was made using Minimac3 software. Due to the unavailability of SNVs on the Y chromosome, we estimated haplotypes for MEs only on autosomes and the X chromosome.

**Haplotype estimation for MEGAnE and MELT callset from EUR in 1000GP GRCh37 datasets**

We generated imputation reference panels from MEGAnE and MELT callsets from 503 European individuals from 1000GP, primarily to evaluate imputation quality between MEGAnE and MELT. With the goal of comparison in mind, we used all ME variants, including those which violate Hardy-Weinburg equilibrium. To perform leave-one-group-out haplotype estimation and imputation, we randomly assigned the 503 individuals into 5 groups composed of either 100 or 101 individuals. We kept the individuals in one group, and made ME callsets from the individuals in the other 4 groups. Singleton MEs in the callsets were removed. The ME callsets were merged with the QC-ed SNV and indel callset (1000GP, v5a) without variants violating Hardy-Weinburg equilibrium ($P < 1 \times 10^{-6}$). The merged callsets were saved as VCF format by each chromosome and phased by SHAPEIT4 software with the default genetic maps. The phased haplotypes were converted to an imputation reference panel for autosomes only using Minimac3. ME genotypes of the individuals in the group left out in the haplotype estimation step were then imputed from the corresponding imputation reference panel (which was generated from the individuals in the other four groups). To simulate the situation of genotype imputation from SNV array data, we only used the SNVs that can be genotyped in the UK BioBank Affimetrix Axiom Array as target haplotypes. Genotypes were imputed by Minimac3 software by each chromosome. We performed this imputation using both the imputation reference panels for MEVs detected by MEGAnE and MELT. We performed this haplotype estimation and imputation for all 5 groups to generate the imputed ME genotypes for all 503 individuals.

**Genotype imputation for GTEx individuals**

To impute ME genotypes in 838 individuals recruited in the GTEx v8, we used the 5006 haplotypes in 1000GP. We used the phased SNVs and indels provided from GTEx (GTEx_Analysis_2017-06-05_v8_WholeGenomeSeq_838Indiv_Analysis_Freeze.SHAPEIT2_phased.vcf.gz) as target haplotypes. Variants violating Hardy-Weinburg equilibrium ($P$ value < 1 x $10^{-6}$) were removed before imputation. ME genotypes on autosomes and PARs were imputed using Minimac3 software with the imputation reference panel generated from the 1000GP GRCh38 callset. After imputation, ME genotypes were extracted and merged with the original SNV and indel calls. MEs violating Hardy-Weinburg equilibrium ($P < 1 \times 10^{-6}$) and/or having Minimac $R^2$ lower than 0.5 were removed. Variants with allele frequency lower than 0.5% were removed, leaving 9,836 MEVs for use in eQTL analysis.

**Genotype imputation in BBJ**

To impute ME genotypes of participants in BBJ, we used the 5,008 haplotypes in the 1000GP GRCh37 dataset. We used phased SNVs genotyped by SNV array as target haplotypes. ME genotypes on autosomes were imputed using Minimac3 software with the imputation reference panel generated from the 1000GP GRCh37 callset. After imputation, variants violating Hardy-Weinburg equilibrium ($P < 1 \times 10^{-6}$) and those with Minimac $R^2$ lower than 0.7 were removed. All variants with minor allele count lower than 10 were removed and the remaining variants were used for GWAS.

### Multiplex Targeted Deep Sequencing

**Design of PCR primers for targeted sequencing**

To enable targeted sequencing, we first aimed to design PCR primers targeting MEI sites detected in Japanese subjects (Fig. S10A). We generated a joint callset from 104 JPT individuals in 1000GP. We also generated a joint callset from 1,235 individuals sequenced at 25x or higher depth in BBJ. We extracted MEIs detected in two or more individuals in the 1000GP callset or MEIs intersecting with CDS, UTR, or promoters of protein coding genes (GTF from GenBank, transcript support level 1). We used Primer3 to design primers amplifying the insertion sites using the following options: minimal amplicon size = 200, maximum amplicon size = 500, primer optimal length = 20, primer minimal length = 19, primer maximum length = 25, primer optimal TM = 60°C, primer minimal TM = 58°C, primer maximum TM = 62°C, and primer optimal GC = 50%. When designing the primers, we specified the primer position in the way that the NGS reads were expected to span the ME insertion breakpoints: specifically, that a 150-bp NGS read would span the breakpoint between the 30th and 120th nucleotide positions. We also filtered out primers that intersected with SNVs common in Japanese (> 1% minor allele frequency in participants in BBJ). After the design of PCR primers, we pooled the primers *in silico* to avoid primers that would generate primer dimers or off-target amplicons when included with those targeting other MEIs. We evaluated primer dimers formation using mfeprimer software version 3.2.0 (*23*). We performed BLASTn similarity search, with the designed primers as a query, against the reference human genome, build GRCh19. After adding one primer set to the *in silico* primer pool, we confirmed the absence of primer dimers and potential off-target amplicons shorter than 10kb; if primer dimers and/or off-targets were predicted, we did not pool the primer set. Because *Alu* (SINE1/7SL) is the most often observed MEIs, we prioritized sequencing L1 and SVA. In addition to the ME sites, we designed control primer sets amplifying regions without common copy number or structural variants; we designed primers in *GAPDH*, *B2M*, *RPPH1*, and *ATP5F1* genes. As a result, we designed a primer pool amplifying 262 ME targets and 4 control targets. As an initial test, we performed multiplex PCR in 166 Japanese and sequenced the amplicons by MiSeq. Some MEVs and control targets failed to be amplified (data not shown). We excluded 102 ME and 2 control targets with low PCR efficiency and finally we used a primer pool targeting 160 MEIs and 2 control genes.

**Targeted deep sequencing and mapping**

We performed the multiplex PCR targeting 162 regions using DNA from 2221 Japanese subjects. We conducted this multiplex PCR in six 384 plates. Barcoded adapter sequences distinguishing individuals were added to the amplicons, which were then pooled and sequenced using an Illumina NextSeq 2000 sequencing system with P2 reagents for 300 cycles. The NGS reads were mapped to the human reference genome, hs37d5. We next counted the number of reads mapping to the target regions. In this step, we only counted the reads overlapping the ME insertion breakpoints. Those spanning reads should derive only from the alleles without MEIs.

**Quality control and normalization of DNA-sequencing data**

We first counted the number of reads sequenced for each individual. We also counted the number of on-target reads. Most individuals (n = 2,027) had more than 100,000 reads (the mean depth of each PCR target is more than 600x depth) (Fig. S10B). Of these, 2,024 individuals had more than 80% on-target mapping rate (Fig. S10C). We used those 2,024 individuals for downstream analysis.

We first checked for a batch effect between PCR plates (Fig. S10D). We visualized the correlation between read count per individual and those of control genes. We found that the PCR amplification efficiency of *GAPDH* was strongly influenced by PCR plate, while that of *B2M* was not affected. Due to the plate-level batch effect, which could presumably affect MEI targets as well, we normalized the data by four different methods (Fig. S11A and B). First the read count of ME targets were normalized by the number of reads mapping to each of *GAPDH* and *B2M*. Then, we further normalized the read counts at plate-level. For each ME target, we calculated the mean read counts of the 6 plates and normalized in such a way that the mean counts of all plates were the same. We visualized the normalized read counts of all individuals for all ME targets and manually selected ME targets with sufficient read counts and clear normalization. The most common cause of exclusion was low read counts, indicative of low PCR efficiency for the specific target (Fig. S11C). Some targets also had discordant normalization patterns between *GAPDH* and *B2M* normalization, so we removed such targets. In total, 110 ME targets (62 *Alu*, 30 L1, 17 SVA, and 1 HERV) were used for downstream analysis.

**Bayesian inference of ME genotypes**

We generated a latent Bayesian model that models three allelic states; alleles with no MEI, heterozygous MEI, and homozygous MEI. In addition, we added an explanatory variable that models the plate-level batch effect. The allele frequency (AF) of ME-absent allele was modeled as a continuous value between 0 and 1. We assumed that the distribution of normalized read counts would be normal, since the depth of targets retained for genotyping was high (the target with the lowest max depth still had 346x maximum depth). The likelihood of being homozygous and heterozygous are described by the normalized read count y, the mean of normalized read count derived from heterozygous ME-absence μ, and the variance of the normalized read count derived from heterozygous ME-absence σ:

Homozygous ME-absence: $p(y\ |AC=2,\mu,\sigma)=Normal\_pdf(y\ |2\mu,\sqrt{2\sigma^2}))$

Heterozygous ME-absence: $p(y\ |AC=1,\mu,\sigma)=Normal\_pdf(y\ |\mu,\sigma))$

No ME-absence: $p(y\ |AC=0)=Normal\_pdf(y\ |0,1))$

In addition to homozygous and heterozygous ME-absence, we modeled the result of having no ME-absence allele (i.e. homozygous ME presence) as a normal distribution with mean 0 and sigma 1, since index hopping in the dataset was negligible. We modeled the plate-level batch effect using hierarchical modeling. We assumed that each of the 6 PCR plates has a PCR amplification coefficient between 0.9 to 1.1 and follows a normal distribution with mean 1. The logarithm likelihood of the model is:

$\sum_{AC=0}^{2}(\ \log\big(p(AC|AF)\big)+\log\big(p(y|AC,\mu\times plate\_coef,\sigma\times plate\_coef)\big)\ ),$

where $plate\_coef=Normal\_pdf(1,\sigma_{plate}))$

We fit the AF of ME-absence, the mean of normalized read count derived from heterozygous ME-absence, the variance of the normalized read count derived from heterozygous ME-absence, plate-level batch effects, and the variance of the plate-level batch effects by MCMC method implemented in Stan (*24*). We ran 5 MCMC chains with 1000 burn-in and 2000 sampling iterations. Convergence was evaluated by calculating Rhat for all parameters, and we excluded any fittings having Rhat larger than 1.2. For 5 ME targets, the model did not converge. For those 5 datasets, we used a simpler model, lacking plate-level batch effects. The fitting of the model for every target was visualized by plotting the original data and the posterior probabilities, and manually checked (Fig. S11A and B).

After model fitting, we inferred the genotypes. We calculated the posterior probability of an individual having an ME-absence allele count 0, 1, and 2. The genotype dosage was calculated by summarizing the product of all possible allele counts (i.e. 0, 1, and 2) and the probability of being the allele count. If the cumulative probability of those was lower than 0.01 or larger than 0.99, we excluded the data point as outlier. As the goal of this analysis was to compare MEGAnE genotypes calls to genotypes determined unambiguously by an experimental method, we also removed data points with ambiguous genotypes (genotype dosage between 1.1 and 1.9; this resulted in removal of 706 data points, or 0.31% of the total data points). The inferred genotype dosage was then rounded to an integer and used for downstream analysis.

### Preparation of Genomic Features

**Gene count**

Gene counts across genome positions were calculated based on the gene annotation from GenCode (GRCh38, version 26). Here, we generated two gene count tables, one considering all genes in the gene annotation, and the other considering only protein coding gene counts. As protein coding genes, we used 19,271 genes that are annotated as "protein_coding" in the attribution column in the GTF file. For all genes, we used 35,041 genes, excluding those annotated as "pseudogenes," "TEC," or "level 3" genes, which represent automatically-annotated loci.

**DNA methylation level**

We calculated the DNA methylation levels at three methylation sites, CpG, CHG, and CHH, in H1-hESCs. We used the DNA methylation levels analyzed by whole-genome bisulfite sequencing from H1-hESC in the ENCODE project (*25*, *26*). For CpG methylation datasets, we used ENCFF601NBW and ENCFF524BMX. For CHG methylation datasets, we used ENCFF379ZXG and ENCFF086MMC. For CHH methylation datasets, we used ENCFF417VRB and ENCFF918PML. All datasets were mapped on GRCh38. Methylation sites with at least 5 reads were used for analysis. To calculate the methylation level for 100kb windows, the methylation levels of all methylation sites in each window were averaged. For downstream analysis, we excluded the windows which have less than 500, 2500, and 10,000 CpG, CHG, and CHH methylation sites, respectively.

The CpG methylation levels in 5 iPSCs from Caucasians, 9 iPSCs from Japanese, and 2 ESCs from Caucasians were also calculated and correlated with population-specific *Alu* (Fig. S33). Methylation data taken by genome tiling array (GSE60821) published by Nishizawa *et al* (*27*) were used. To generate methylation status data on GRCh38, positions of the array probes were lifted over from GRCh37 to GRCh38. Out of 485,512 probe positions, 484,111 were successfully lifted over. Then, the methylation levels for 100kb non-overlapping windows are calculated by averaging methylation levels of probes fell in each 100kb window. If a given window does not contain 3 or more probes, such a window was removed from analysis.

**Replication timing**

We calculated the replication timing (RT) for H1-hESC and H7-hESC from Repli-Chip datasets in the ENCODE project. For RT of H1-hESC, we used ENCFF000KUF and ENCFF000KUG. For RT of H7-hESC, we used ENCFF000KUK. The Repli-Chip probes were mapped on GRCh37 in the original data, thus we lifted over the probe positions to GRCh38. We performed reciprocal liftover of probe positions and determined the corresponding genome positions on GRCh38. Out of 2,154,503 probe sites, 2,148,824 probes were reciprocally lifted over (5,679 probes failed reciprocal liftover). Based on the lifted over probe positions, we recalculated RT. To calculate the methylation level for 100kb windows, windows with at least 25 probes were used and the average RT of each window was calculated.

**A/B compartment**

We inferred the A/B compartments in H1-hESC. We used the Hi-C data taken by Oksuz *et al* (*28*). We used two datasets, Hi-C processed by formaldehyde fixation followed by HindIII cleavage

(GSM5057489) and formaldehyde and DSG fixation followed by HindIII cleavage (GSM5057481). We downloaded the alignments in the mcool format, and PCA of each 100kb window was performed using the cooltools Python package. The first eigenvector was then used to infer A/B compartments.

**Histone modifications**

We calculated the number of histone modification peaks in H1-hESC. We used the histone Chip-seq datasets in the ENCODE project. The histone modifications we used and the accession numbers of the datasets are listed in Table S8. The number of peaks in every 100kb window was calculated for each dataset. If there are replicates, we calculated the average number of peaks in the 100kb windows.

**DNase hypersensitive sites**

We calculated the number of DNase-seq peaks in H1-hESC. We used the histone DNase-seq datasets in the ENCODE project. The accession numbers are ENCFF905XDS, ENCFF574LKL, and ENCFF338KTY. The number of peaks in every 100kb window was calculated for each dataset. Then, we calculated the average number of peaks in the 100kb windows across the three datasets.

**TF-binding sites**

We calculated the number of CTCF, phospho-Pol-II A, Pol-II, and EP300-binding peaks in H1-hESC. We used the TF Chip-seq datasets in the ENCODE project, for provide peak calls in BED format. The accession numbers are ENCFF821AQO, ENCFF418QVJ, ENCFF422HDN, and ENCFF834UVX, respectively. The number of peaks in every 100kb window was calculated for each dataset.

### Analyses Related to MEVs

**Detection of motifs at insertion breakpoints**

To detect the DNA sequence motifs that enrich at insertion breakpoints, we used the "findMotifs.pl" script in HOMER software (*29*). For this analysis, we used the MEI callsets from 1000GP GRCh38 datasets, SFARI, and BBJ 25x WGS (Table S7). We only used MEGAnE MEI calls that have the filter "PASS" flag and a predicted insertion direction. 14-bp sequences, composed of 4-bp upstream and 10-bp downstream from the predicted insertion breakpoint, were compiled and used as the target sequences. As control sequences, random 14-bp sequences from the human reference genome, GRCh38DH, were selected using the bedtools `shuffle` command. We used 10 times the number of control sequences as actual MEI breakpoints. The consensus sequence that is recognized by L1 endonuclease (T/AAAA) is enriched at the breakpoints of all TPRT-mobilized ME families, but not HERV-K (Fig. S34).

**PCA of MEVs**

The principal components (PCs) of ME polymorphisms called from 1000GP GRCh37 datasets and the SFARI cohort were calculated by Plink2 software. We first removed MEVs violating Hardy-Weinburg equilibrium ($P < 1 \times 10^{-6}$), those with minor allele frequency lower than 1%, and those in regions of long-range high linkage disequilibrium (https://genome.sph.umich.edu/wiki/Regions_of_high_linkage_disequilibrium_(LD)). The variants were then pruned wby Plink2 software with `--indep-pairwise 500 5 0.2` option. The top 10 PCs were calculated using the plink2 --pca command.

**Intersections between MEVs and gene annotations**

To compile MEVs that intersect with exons, CDS, and promoters, we first reshaped gene annotation files downloaded from GenCode using a script provided in the GTEx pipelines (https://github.com/broadinstitute/gtex-pipeline/blob/master/gene_model/collapse_annotation.py). We defined the 1kb regions upstream from transcription start sites as promoters. All gene annotations in the GTF file were used for this analysis. To see intersection with MEVs called from 1000GP, we used 48,241 MEVs with the filter "PASS" flag called from 1000GP GRCh38 datasets. For this analysis, we used a GenCode GTF version 26. To see intersection with MEVs called from BBJ, we used 10,997 MEVs with the filter "PASS" flag called from 1,235 individuals sequenced at 25x depth WGS. For this analysis, we used a GenCode GTF version 26lift37.

**Correlations between ME insertions and genomic features**

To evaluate the characteristics of genome features found to have insertions of MEs, the correlation between the number of ME insertions and genomic features were calculated. We calculated the genomic features for non-overlapping 100-kb windows (see the “Preparation of genomic features” section).  Because L1 and SVA insertions are sparse, we first resized the window size to 1 Mb and 5 Mb, respectively. To this end, the average values were calculated for each non-overlapping window. 1Mb and 5Mb windows that contain one or more 100-kb window(s) with missing value and ones with at least one ‘N’ character in the human genome assembly, GRCh38DH, were excluded from the analysis. The Spearman correlation coefficients were calculated using the SciPy module in Python.

**Preferential insertion of MEVs in the non-transcribed strand of genes**

L1HS copies in the reference human genome and the experimental L1HS insertions from a reporter plasmid are more frequently found in non-transcribed strand than transcribed strand (*30*). To check whether the MEIs enrich in the non-transcribed strand of genes, we compiled all MEIs found in SFARI that intersect with gene annotations from GenCode (v26, GRCh38). We removed non-transcribed genes annotated as "pseudogene," "TEC," and automatically-annotated "level 3" genes from this analysis. As a result, we observed the same trend in family-specific heritable L1HS, *Alu*, and SVA (Fig. S35A). This suggests that the three polymorphic MEs may preferentially target accessible single-stranded DNA at transcribed loci. We thus evaluated the early embryonic and germline developmental stages at which genes containing recent ME insertions are often expressed. To characterize genes expressed during spermatogenesis and early embryonic stages, we used the gene expression datasets produced by scRNA-seq published in Guo *et al* (*31*) and Yan *et al* (*32*). The gene expression matrices of single cells were converted to pseudo-bulk expression tables grouped by cells annotated as spermatogonia, spermatocytes, early spermatids, late spermatids, oocytes, zygotes, 2-cell, 4-cell, 8-cell, morulae, trophectoderm, endoderm, and epiblast. Family-specific heritable L1 and *Alu* insertions were frequently found in genes highly-expressed in spermatids (Fig. S35B). The same trend was also observed in *de novo Alu* insertions (Fig. S35C). This suggests the possibility that heritable *Alu* and L1 insertion may often occur in spermatids.

**Intersections between ME-eQTLs and ENCODE regulatory elements**

We tested the overlap between MEVs in ME-eQTLs and ENCODE regulatory element annotations, i.e. cCRE and DNase hypersensitive sites (DHS). For pELS (proximal enhancer-like signatures), dELS (distal enhancer-like signatures), CTCF (CTCF-bounded signatures), PLS (promoter-like signatures), the hg38 ENCODE cCRE dataset (https://screen.encodeproject.org) was used. For DHS, those from ENCODE (ENCFF503GCK) were used. Out of 1,073 ME-eQTLs, MEVs in 101 and 221 ME-eQTLs intersected with ENCODE cCRE and DHS annotations, respectively. Of 101 ME-eQTLs, MEVs in 78, 16, 4, and 3 ME-eQTLs intersect with dELS, pELS, PLS, and CTCF-peak, respectively. Out of non-reference MEVs in 668 ME-eQTLs, those in 45, 10, 4, and 2 ME-eQTLs intersect with dELS, pELS, PLS, and CTCF-peak, respectively.

**Occurrence of ME-eQTLs by genome features**

To understand how MEVs detected as ME-eQTLs differ from non-ME-eQTL MEVs based on genome features, we calculated odds ratios. We treated genome features as the exposure and being detected as the MEV of an ME-eQTL (which occurred for 778 of 9,836 MEVs used for eQTL analysis) as the outcome. For A/B compartments, we used the first principal component of Hi-C (see the "Preparation of genomic features" section). For early-replicating domains, we used replication timing, as determined from Repli-Seq, greater than 0 (see the "Preparation of genomic features" section). MEs in the genome windows lacking replication timing and/or A/B compartment information were excluded from the analysis. We used intersection with the called ChIP peak ranges from ENCODE as indicator of histone modification (Table S8). For exon, intron, promoter, and intergenic regions, we used the gene models provided from GenCode (GRCh38, v26).

To assess the occurrence of significant ME and gene expression association, we calculated odds ratios. In the eQTL analysis in the 49 tissues, 266,530 unique ME-gene pairs were tested for association. Of these, 1,073 ME-gene pairs that are significant in at least one tissue (LFSR $< 0.05$), i.e. ME-eQTLs, were used as the outcome, and the same genome features described in the previous paragraph were used as exposure. MEs in genome windows lacking replication timing and/or A/B compartment information were excluded from analysis.

### ME-eQTL Analysis in GTEx

**eQTL analysis in 49 tissues**

We performed eQTL mapping using MEVs. We followed the eQTL mapping method used in GTEx v8. As for GTEx v8, we excluded 5 tissues out of the 54 tissues (Bladder, Cervix_Ectocervix, Cervix_Endocervix, Fallopian_Tube, and Kidney_Medulla) from analysis due to the few available RNA-seq samples. First, expression profiles of the 49 tissues were prepared. The TPM count matrices provided from GTEx (GTEx_Analysis_2017-06-05_v8_RNASeQCv1.1.9_gene_tpm.gct) were normalized across samples by TMM normalization using the script provided from GTEx (https://github.com/broadinstitute/gtex-pipeline/blob/master/qtl/src/eqtl_prepare_expression.py), then the genes that are expressed (≥ 0.1 TPM in ≥ 20% samples and ≥6 reads in ≥20% samples) were retained for eQTL mapping (38,471 genes in total of 49 tissues). Each gene was then inverse-normal transformed across samples. Next, we performed eQTL mapping by fastQTL software (*33*) with the same analysis options as for the previous eQTL mapping (https://github.com/broadinstitute/gtex-pipeline/tree/master/qtl). We also used the same covariates as those used for QTL mapping in GTEx; 5 genetic PCs, PEER factors, library preparation methods, sequencing platforms, and sex. Genetic variants within 1Mb from a gene were tested for associations. The 9,836 and 13,498,030 QCed ME and non-ME (i.e. SNVs and indels) variants, respectively, were used for eQTL mapping.

**Exon expression-QTL analysis in 49 tissues**

To assess the association between L1-NEDD4 and *NEDD4* exon expression, we performed exon expression-QTL analysis. First, the expression profile of *NEDD4* exons was prepared. The read count table was downloaded from GTEx (https://storage.googleapis.com/gtex_analysis_v8/rna_seq_data/GTEx_Analysis_2017-06-05_v8_RNASeQCv1.1.9_exon_reads.parquet). To apply TMM normalization, we used the normalization factor calculated from the read count table of genes (GTEx_Analysis_2017-06-05_v8_RNASeQCv1.1.9_gene_tpm.gct); the exon expression values (TPM) are lower than those of the genes, meaning we would remove a larger fraction of read counts before TMM normalization if normalizing only by exon read counts. We removed genes with read count less than 6 and calculated TMM normalization factor for each tissue. Using the TMM normalization factors, the exon expression profiles across samples were normalized, then, each exon was then inverse-normal transformed across samples. The exon expression-QTL mapping was done using the same protocol as eQTL mapping; we used as covariates the first 5 genetic PCs, PEER factors, library preparation methods, sequencing platforms, and sex. Genetic variants within 1Mb from the *NEDD4* gene were tested for associations. Thirty-four *NEDD4* exons in the GenCode gene annotation (GRCh38, v26) were used for analysis. FastQTL software was used.

**Conditional analysis**

To detect all haplotype blocks associated with *NEDD4* gene expression, we performed conditional analysis by including the genotypes of lead variants as covariates. The same expression profiles and software as eQTL analysis were used. We conditioned the QTLs serially until no significant associations were detected.

**Across-tissue meta-analysis**

After the eQTL mapping in each tissue, we performed across-tissue meta-analysis using the same method as performed in GTEx v8. First we formatted the fastQTL results for MASH software (*34*). Then the MASH model was trained by the same protocol as GTEx v8 performed (https://github.com/stephenslab/gtexresults/blob/master/workflows/fastqtl_to_mash.ipynb). The trained model was applied to ME-eGene pairs.

**Detection of ME-eQTL**

We defined ME-eQTLs as those which satisfy these criteria: 1) in the fastQTL output, an MEV is either the lead variant or has $R^2 > 0.95$ to the lead variant in at least one tissue, and 2) in the result of across-tissue meta analysis, the MEV has local false sign rate (LFSR) < 0.05 in at least one tissue.

**Proxy gene mapping**

We used a linear mixed model to detect the association between eGenes (genes that are associated with MEVs, in this case, 3' UTR *Alu* elements) and pGenes (proxy genes). To perform across-tissue regression, we modeled according to the rationale below.

1) The eGene and pGene expression may be correlated even if the eGene does not have *Alu* in its 3'UTR.

2) Correlation coefficient between eGene and pGene is different depending on the presence and absence of *Alu*. In other words, there is the eGene and pGene interaction.

3) The expression level of eGene is different between tissues regardless of the level of pGene expression. This can be considered as a random effect of tissue (tissue is a moderator of the intercept of eGene-pGene correlation).

4) The expression level of eGene depends on *Alu* genotype regardless of the level of pGene expression. This can be considered as a random effect of *Alu* on eGene expression (*Alu* is a moderator of the intercept of eGene-pGene correlation).

5) Furthermore, the effect described above may be different between tissues, due to the difference of other proxy gene expressions which influence this. In other words, the interaction between *Alu* and tissue is the moderator of the intercept of eGene-pGene correlation.

6) The effect of pGene on the eGene expression may be different between tissues. This can be considered as a random effect of tissue on the effect of pGene to eGene. In other words, tissue is a moderator of the slope of eGene-pGene correlation.

The full model can be described as below. To test the significance of *Alu*:pGene interaction, we performed a likelihood ratio test between the full model and the restricted model, which does not include an *Alu*:pGene interaction term.

Full model: *eGene ~ pGene + Alu:pGene + (1 | tissue) + (1 | Alu) + (1 | tissue:Alu) + (0 + pGene | tissue)*

Restricted model: *eGene ~ pGene + (1 | tissue) + (1 | Alu) + (1 | tissue:Alu) + (0 + pGene | tissue)*

We applied this model to 20 *Alu*-eGene pairs in the tissues in which the pairs are detected as ME-eQTL. Out of the 20 pairs, 17 were detected as ME-eQTL in multiple tissues, while 3 pairs were tissue-specific. For those three, we applied a simpler model that does not have the tissue term in the model. We fit this model using the lme4 R package (*35*). The gene expression profiles used for the proxy gene mapping was prepared from the TPM count matrices provided from GTEx (GTEx_Analysis_2017-06-05_v8_RNASeQCv1.1.9_gene_tpm.gct) using the same method as used for the eQTL mapping (see "eQTL mapping in 49 tissues" section). For pGenes, we only used those that have TPM larger than 2 based on an assumption that low expression of a pGene is less likely to impact eGene expression in a relevant manner. Each gene was then inverse normal transformed across samples. Then, we fit the same covariates used for the eQTL mapping (see "eQTL mapping in 49 tissues" section) and calculated the residuals. The residuals were inverse normal transformed across samples in such a way that the data will have the same mean and standard deviation as the original data (TMM-normalized TPM). We fit the linear mixed model described above to this expression data. We excluded genes that fall within 1Mb from eGene from proxy gene mapping, because expression levels of neighboring genes may show a collinear relationship due to transcriptional regulation by the same regulatory elements.

**Gene-set enrichment analysis**

We performed Gene-set enrichment analysis (GSEA) using the *P* values of proxy gene mapping results. The genes used for proxy gene mapping were ranked by $-\log_{10}$ transformed *P* values and tested for enrichment of the 7,481 gene-sets in the "biological processes" category of the Gene Ontology database (c5, version 7.4, (*36*, *37*)) using fgsea R package (*38*). After performing GSEA independently for all 20 *Alu*-eGene pairs, we performed across-*Alu* meta-analysis by Fisher's combined probability test and calculated the *P* value for each pathway used in GSEA.

**Permutation of MEVs in ME-eQTLs to test linkage disequilibrium with the GWAS catalog variants**

Of the 778 MEVs associated with at least one gene expression in one tissue, 102 MEVs were tagged with variants listed in the GWAS catalog v1.0.2 ($R^2$ > 0.8) (*39*). To test whether MEVs detected as ME-eQTLs are more often tagged with GWAS catalog variants than non-ME-eQTL MEVs, we performed permutation analysis. We selected the same number of MEVs from the 9,836 MEVs, which were used in eQTL analysis, in such a way that the distribution of allele frequency and distance from TSS after permutation becomes the same as that of before permutation. To this end, we split the allele frequencies into 50 bins and distances from TSS into 3 bins (1-100kb, 100-500kb, and 500-1000kb). We performed 1,000 permutations and calculated the number of MEVs tagged with GWAS catalog variants. The 1,656 haplotypes from GTEx v8 (828 individuals) were used for calculation of linkage disequilibrium.

### Associations Between MEVs and Traits

**ME-GWAS of 42 diseases in BBJ**

GWAS for 42 diseases were done using 179,660 individuals in BBJ using methods similar to those used in Ishigaki *et al* (*40*). The MEV genotypes in 179,660 individuals were imputed by Minimac3 software using the imputation reference panel generated from the 1000GP GRCh37 datasets. After imputation, variants violating Hardy-Weinburg equilibrium ($P < 1 \times 10^{-6}$), those with Minimac $R^2$ lower than 0.7, and those with a minor allele count lower than 10 were removed. The associations were calculated using a generalized linear mixed model implemented in SAIGE (version 0.44.5 (*41*)) with the leave-one-chromosome-out approach. We used age, sex, and the first 5 genetic pricipal components as covariates. For each disease, we defined a significantly-associated locus as a genomic region within 3 Mb from the lead variants. Based on the methodology used in Ishigaki *et al*, we used $9.58 \times 10^{9}$ as a genome-wide significance threshold and $5 \times 10^{8}$ as a threshold of suggestive association.

**Conditional analysis**

To detect all MEV and disease associations, conditional analysis was performed. All the GWAS peaks with MEVs were conditioned with the lead variants until all the significant associations ($P < 5 \times 10^{-8}$) were conditioned out. For this, the `--condition` option implemented in SAIGE was used.

**Calculation of LD between MEVs and lead variants in UKB**

7221 summary statistics were downloaded from the URLs listed in https://pan-ukb-us-east-1.s3.amazonaws.com/sumstats_release/phenotype_manifest.tsv.bgz (Pan-UKBB). To identify lead variants, we selected variants that have a *P* value for EUR smaller than $5 \times 10^{-8}$ in 3Mb windows; this amounted to 169,822 lead variants across all summary statistics (Table S9). To check for linkage disequilibrium, we calculated $R^2$ between MEVs and SNVs using 5,008 haplotypes in the 1000GP (Table S10).

**Calculation of LD between MEVs and lead variants in BioBank Japan**

The "tophit" variants in BBJ were downloaded from the following URLs: http://jenger.riken.jp:8080/top_hits and https://pheweb.jp. We excluded the results of eQTL mapping and meta-analyses from this analysis. If two lead variants detected by the two GWASs of the same phenotype in the two datasets fall within 1Mb each other, we considered the risk loci to be the same and merged into one locus. In total, the two datasets include 172 GWASs consisting of 4,369 lead variants with *P* value lower than $1 \times 10^{-5}$. The linkage disequilibrium between MEVs and SNVs were calculated using the genotypes identified from 25x WGS of 1,235 Japanese individuals (Table S11).

### Cell-Based Experiments

**Cell lines**

The following LCLs were obtained from the NIGMS Human Genetic Cell Repository at the Coriell Institute for Medical Research: GM12878, GM18954, GM19088, GM20787, GM18944, and GM18999. iPSCs were provided by the RIKEN BRC through the National BioResource Project of the MEXT, Japan. Oligodendroglioma cells were a gift from Dr. Keizo Tomonaga of Kyoto University. NT2/D1 cells were obtained from ATCC (CRL-1973).

**PCR validation of L1-NEDD4**

We checked the presence of the detected *NEDD4* intronic L1 insertion in 70 healthy Japanese by PCR using genomic DNA purified from the iPSCs described above. Out of 70 individuals, 10 and 26 individuals harbored homozygous and heterozygous L1 insertion, respectively (Fig. S41A). The experimental allele frequency in these 70 individuals (32.9%) is consistent with the frequency called by MEGAnE in Japanese (35.4%). Primer sequences used are listed in Table S12.

**Knockout of L1-NEDD4 in iPSCs**

We designed two sgRNAs cleaving upstream and downstream of L1-NEDD4 insertion. To reconstruct the allele without the L1-NEDD4, we amplified the L1-flanking regions (703 bp upstream and 787 bp downstream) and connected them at the TSD using overlap-extension PCR. The connected fragment was used as a template for homology directed repair (HDR). The sgRNA-Cas9 complex and HDR template DNA were transfected to iPSCs derived from a healthy Japanese subject (60s, Male) found to carry two copies of L1-NEDD4 by electroporation using the NEON transfection system. After electroporation, cells were cultured for two weeks and single cell-derived clones were obtained by limiting dilution. Deletion of L1-NEDD4 was checked by the same primers as used for PCR validation in 70 Japanese (Fig. S41B).

**Differentiation of iPSCs into fibroblasts**

iPSC clones were first differentiated to mesenchymal stem cells (MSCs) using STEMdiff Mesenchymal Progenitor Kit according to the manufacturer's protocol. iPSC-derived MSCs were then differentiated to fibroblasts based on the protocol published in Lee *et al* (*42*). MSCs were cultured in DMEM containing 100 ng/ml CTGF, 50 ng/ml ascorbic acid, 1x penicillin/streptomycin, and 10% FBS for at least three weeks. Fully-differentiated fibroblasts were maintained in the same medium used for MSC to fibroblast differentiation.

**qRT-PCR of *NEDD4* transcripts**

To measure the expression levels of *NEDD4* in fibroblasts, we collected L1-NEDD4 KO and WT clones differentiated into fibroblasts and extracted total RNA. Polyadenylated RNA were reverse-transcribed using oligo-dT primer. To measure the expression level of the long transcript variant of *NEDD4*, we designed primers in the long-variant-specific exons (exon 1 and 8). To measure expression of the short transcript variant of *NEDD4*, we designed primers amplifying the junction of the short-variant-specific exon (exon 9) and an exon that are shared in both short and long variants (exon 14), because exon 9 is the only exon that is specific to the short variant. Beta-actin transcript was used as an internal control. We also measured the expression of *GAPDH* and the linearity between beta-actin and *GAPDH* expressions across samples was confirmed. The relative expression levels of the *NEDD4* transcripts were calculated by ΔΔCt method. We serially diluted cDNA to confirm that the qPCR conditions used resulted in exponential amplification. qPCR was performed on ViiA7 Real-Time PCR System using SYBR Green reagent. The sequences of the primers are listed in Table S12.

**Luciferase reporter enhancer assay**

The region downstream of TRIM25 gene, chr17:56870020-56870562 on GRCh38, was amplified from NA18999 by PCR. This individual is heterozygous for the *Alu* insertion in this region. We cloned the alleles with and without *Alu* into a series of luciferase enhancer reporter plasmids used in Andersson *et al* (*43*). We sequenced the cloned fragments and confirmed that the two alleles do not have any sequence difference besides the *Alu* insertion. The plasmids were transfected to Lenti-X 293T cells in 24 well plates using Xfect reagent and luminescence was measured at 24 hours post transfection. A plasmid coding for Renilla luciferase with a CMV promoter was used as a transfection control.

**Luciferase reporter 3'UTR assay**

The 3'UTRs of *HSD17B12* (GRCh38, chr11:43855249-43856615), *EGFR* (GRCh38, chr7:55205618-55211628), *ADIPOQ* (GRCh38, chr3:186854705-186858463), and *MAP3K21* (GRCh38, chr1:233382712-233385148) were amplified by PCR from NA18954, NA19088, NA20787, and NA18944, respectively, and cloned. These individuals carry at least one copy of the 3'UTR with the *Alu* insertion, and colonies were screened for the *Alu*-present allele. Isogenic 3'UTR sequences without *Alu* insertions were generated by deleting the *Alu* insertion using overlap-extension PCR and sequenced to confirm absence of unintended mutations. The 3'UTR sequences were cloned downstream of a firefly luciferase CDS under control of a CMV promoter. The plasmids were transfected into GM12878 LCLs by electroporation using the NEON transfection system, and to OL cells and NT2/D1 cells using Xfect reagent. Luminescence was measured 24 hours post transfection. A plasmid coding for Renilla luciferase with a CMV promoter was used as a transfection control.

**Luciferase reporter 3'UTR assay with transient FAM120A expression**

The coding sequence of *FAM120A* was amplified by PCR from cDNA generated from total RNA purified from GM18954. The coding sequence without the termination codon was cloned to the pTag1-CMV vector (pTag1-CMV-empty) in such a way that the ORF of *FAM120A* is connected with the frame coding Flag tag in the vector (pTag1-CMV-FAM120A-Flag). The plasmids coding for firefly luciferase and 3’UTR of *ADIPOQ* with or without *Alu*, *Renilla* luciferase, and FAM120A-Flag were transfected to GM12878 LCLs by electroporation using the NEON transfection system. To evaluate the effect of FAM120A dosage on reporter activity, 0, 0.4, and 0.8 μg of pTag1-CMV-FAM120A-Flag and 0.8, 0.4, and 0 μg of pTag1-CMV-empty, respectively, were transfected with 0.2 μg of the reporter plasmids.

### Definitions

**Box plot**

Lower whisker represents the lowest datum above Q1 – 1.5 x (Q3-Q1), and upper whisker represents the highest datum below Q3 + 1.5 x (Q3-Q1), where Q1 and Q3 are the first and third quartiles. All box plots were generated by the Python matplotlib module.

#### Software Versions

- Python 3.7.4
- MEGAnE v0.1.1
- pandas 1.3.1
- numpy 1.19.2
- seaborn 0.11.2
- matplotlib 3.3.2
- scipy 1.7.3
- statmodels 0.12.2
- scikit-learn 0.22.1
- pystan 2.19.1.1
- arviz 0.11.2
- cooltools 0.5.0
- hdf5 1.10.2
- biopython 1.74
- pysam 0.15.2
- R 3.6.1
- lme4 1.1.27.1
- mashr 0.2.38
- fgsea 1.19.3
- coloc 4.0.4
- SMR 1.03
- SAIGE 0.44.5
- BWA 0.7.17
- Minimac3 2.0.1
- Minimac4 1.0.2
- SHAPEIT4 4.1.3
- phenogram 1.2.1
- samtools 1.10 and 1.14
- bedtools v2.29.2
- bcftools 1.9
- PLINK v1.90b6.17
- PLINK v2.00a2.3LM
- RepeatMasker 4.0.9
- MELT 2.1.5

### Supplementary Figure Legend

**Fig. S1. Illustration of data processing by MEGAnE**

MEGAnE searches for discordantly mapped reads (chimerically mapped reads, distantly-mapped reads, and unmapped reads) and finds potential breakpoints from clipped reads. It pairs breakpoints likely to represent the upstream and downstream breakpoints of an ME insertion or absence. After removing likely false positives and estimating genotype, it outputs MEVs as VCF format. See Supplementary Materials for more detailed algorithms.

**Fig. S2. Removal of likely false positives by MEGAnE**

(A) Fitting of a Gaussian mixture model to the number of MEI breakpoint-supporting reads. After breakpoint calling and pairing, MEGAnE determines the model, shown by a red line, that best fits the distribution of the number of MEI breakpoint-supporting reads, shown as blue dots. In the case of human samples, there are two peaks representing homozygous and heterozygous MEIs. MEGAnE fits two Gaussian distributions, shown as two gray lines, to the two peaks and determines a cutoff threshold to remove likely false positives which are evidenced by only a few breakpoint-supporting reads. Here, the result of an individual, NA12878, is shown as an example. (B) Cutoff thresholds determined for 2,504 individuals of 1000GP 30x WGS dataset. Cutoff threshold shows correlation with the depth of WGS, showing that appropriate thresholds can be determined based on the characteristics of the input dataset.

**Fig. S3. Criteria used for genotyping MEIs by MEGAnE**

In this figure, results of one individual, NA12878, are shown as an example. (A and B) IGV genome browser view showing increased (A) and decreased (B) read depths at non-reference ME insertion sites. Differential read depth due to TSD (A) or a short deletion (B) at an MEI breakpoint is characteristic of non-reference MEIs (shown with red arrows). MEGAnE uses such read depth information to accurately determine genotypes of MEIs. (C) Distribution of the relative depths at TSDs (depths at each TSD divided by depth at the regions adjacent to each TSD) and the number of breakpoint-spanning reads. (D) Distribution of the relative TSD depths and the number of breakpoint-supporting reads. (E) Distribution of the relative depth at short deletion (depths at short deletions divided by those at regions adjacent to the short deletions) and number of breakpoint-spanning reads. (F) Distribution of the relative depth at short deletion and the number of breakpoint-supporting reads. MEGAnE genotypes MEIs as “0/1” when abundant breakpoint-spanning reads are detected. On the other hand, if there are a few or no breakpoint-spanning reads, but the number of breakpoint-supporting reads and the depth at TSD or short deletion are high relative to the adjacent regions, it genotypes as “1/1.”

**Fig. S4. Criteria used for genotyping ME absences by MEGAnE**

In this figure, results of an individual, NA12878, are shown as examples. (A) IGV genome browser view showing decreased read depth at an absent reference MEs. At the edges of ME absences, there are sharp decreases in read depth, such as ones shown with red arrows. MEGAnE uses this read depth information to determine genotype of ME absences. (B) Distribution of the relative depths (the depth at one edge of a ME absence divided that at the adjacent region) upstream and downstream of edges of ME absences. (C) The distribution of the relative depth at the edge of ME absences either upstream or downstream (whichever is lower), and the distribution of the breakpoint-spanning reads. MEGAnE genotypes MEIs as "0/1" when abundant breakpoint-spanning reads are detected. On the other hand, if there are a few or no breakpoint-spanning reads, but the depths at the edges of MEs are enough low, it genotypes as "1/1."

**Fig. S5. Comparison of MEV discovery between MEGAnE and PAV using all ME-containing SVs**

Here, we used all long-read-resolved SVs identified by PAV that overlap with MEs (*3*) as ground truth. These include variants that are unlikely to be TPRT-mediated insertions. (A and B) True positive rate of MEVs discovered by MEGAnE. MEGAnE was applied to 34 individuals sequenced by long-reads and MEVs were called. Polymorphic MEIs and ME absences are cross-referenced with the SVs identified by PAV and the true positive rate was calculated. (A) The result of the filter-"PASS" variants are shown. (B) The result of the variants labeled as filter not-"PASS" are shown. (C) Sensitivity of MEV discovery by MEGAnE. SVs identified by PAV were grouped based on the genome annotations at the locations of the SVs and evaluated the detection sensitivity. (D) Numbers of MEIs missed by MEGAnE. MEGAnE missed 1,117 SVs harboring ME sequences (C, highlighted by a blue circle). These are grouped by ME family and whether the SVs contain a recently-active MEs. (E) Accuracy of breakpoint positions of MEIs determined by MEGAnE. The MEI breakpoint positions predicted by MEGAnE were compared to those determined by PAV. Relative distances between the breakpoint positions between MEGAnE and PAV are shown as a histogram.

**Fig. S6. Comparison of MEV discovery between MEGAnE and PAV using likely TPRT-mediated insertions**

(A and B) Distribution of lengths of ME absences (A) and insertions (B). The top graphs shows the length distribution of long-read-resolved SVs that overlap with any MEs, while the middle (A) and bottom (B) graphs show the length distribution of SVs that show evidence of TPRT-mediated mobilization (i.e. presence of TSD and polyA tail). The bottom panel of (A) shows the length distribution of ME absences detected by MEGAnE. MEGAnE does not resolve the length of ME insertions. (C and D) Recovery of MEIs (C) and ME absences (D) which are mobilized by TPRT-mediated mechanism. Here, we used SVs identified by PAV (*3*) that have evidence of insertion via TPRT for analysis.

**Fig. S7. Comparison of MEV discovery and genotyping accuracy between MEGAnEand MELT**

We applied MEGAnE and MELTv2 to the 30x WGS of 503 EUR individuals sequenced by the 1000GP. (B-D) We used MEVs that are genotyped by PanGenie in the 503 individuals (*3*) as ground truth. (A) Comparison of the number of MEVs discovered by MEGAnE and MELT. Left and right panels show MEIs and ME absences, respectively. (B and C) We used MEVs that are genotyped by PanGenie in the 503 individuals (*3*) as ground truth. (B) Counts of MEVs that are detected and genotyped by MEGAnE and MELT. Left and right panels show MEIs and ME absences, respectively. The bar plots are stratified by $R^2$ between genotypes from PanGenie and

MEGAnE or MELT. (C and D) Accuracy of MEV genotyping by MEGAnE (C) and MELT (D). Left and right panels show MEIs and ME absences, respectively. To evaluate whether the genotyping accuracy is related to MEV allele frequency (x axis), $R^2$ between genotypes from PanGenie and MEGAnE or MELT (y axis) is plotted.

**Fig. S8. Comparison of MEV genotyping accuracy between MEGAnE and GATK-SV**
We applied MEGAnE to the 30x WGS of 3,202 individuals in the 1000GP. As a GATK-SV result, the callset generated by Ebert *et al.* (*3*) is used. We used MEVs that are genotyped by PanGenie in 3,202 individuals in the 1000GP (*3*) as ground truth. (A and B) Comparison of MEV genotyping between PanGenie and MEGAnE (A) or GATK-SV (B). Left and right panels show MEIs and ME absences, respectively. The color of the dots represents the $R^2$ of genotypes between PanGenie and MEGAnE (A) or GATK-SV (B). (C) Counts of MEVs that are detected and genotyped by MEGAnE and GATK-SV. Left and right panels show MEIs and ME absences, respectively. The bar plots are stratified by $R^2$ between genotypes from PanGenie and MEGAnE or GATK-SV.

**Fig. S9. Mendelian error rate of MEVs detected by MEGAnE**
(A) Mendelian error rate (MEr) of MEIs determined from 602 trios. MEGAnE was applied to the 602 trios sequenced by the 1000GP and MEr was determined. (B) MEr of family-specific MEIs. MEr determined from MEIs that are found in only one trio of the 602 families is shown.

**Fig. S10. Processing of targeted deep sequencing data; quality control**
(A) Illustration showing the experimental design of MEV genotyping by multiples targeted deep sequencing. PCR amplifies 160 target sites enclosing MEVs. We designed the PCR primers in such a way that NGS reads should contain a fragment of ME if the primers amplified an allele with MEI. We used the NGS reads that detected the ME absent allele to genotype MEVs. (B) Number of target-sequencing reads by the 2,221 individuals. Individuals with a read count lower than the cutoff threshold shown as a gray dashed line were removed from analysis. (C) On-target rate of the target-sequencing by the 2,221 individuals. Proportion of on-target (blue) and off-target (orange) are shown. Individuals with on-target rate lower than the cutoff threshold shown as a gray dashed line were removed from analysis. (D) Number of reads mapping to the two control targets, GAPDH and B2M genes. The 2,221 samples are colored by the six PCR plates of which the samples were amplified, showing a plate-level batch effect in the case of GAPDH.

**Fig. S11. Processing of targeted deep sequencing data; target-level normalization and inference of genotypes**
(A) Schematic of target-level normalization, model fitting, and inference of genotypes. (B) One example of target-level normalization, fitting a model, and genotype inference. This example is the target sequencing of a region, GRCh37:chr12:84965132-84965148, which contains a polymorphic L1 insertion. First, the raw read counts for the 2,024 individuals who passed quality-control were normalized by four different methods, and the normalized read counts were visualized (upper 4 panels). By manual checking of the normalized read count distribution, we selected one normalization method which clearly differentiates the individuals with no, heterozygous, and homozygous insertions (the distribution shown with blue dots). Simultaneously,

we manually set a cutoff threshold to remove outlier points (the red vertical line shown in the top fourth panel). Then, the Bayesian model was fit to the selected distribution by MCMC and used to estimate the posterior probability of carrying no or heterozygous insertion. Based on the posterior probability, we inferred the dosage of having ME-absent allele (i.e. pre-integration allele). In the panels labelled "original data with inferred genotypes" and "inferred genotype (dosage)" those genotyped as 0/0, 0/1, and 1/1 of ME-absence alleles are shown as light, intermediate, and dark blue colors. We removed ambiguously genotyped individuals, based on the posterior probabilities (the dots colored with orange in the top 11th and 12th panels represent ambiguously genotyped data points). (C) Maximum read depth at each PCR target across quality-controlled individuals. The PCR targets excluded during target-level normalization, while yet unaware of concordance with MEGAnE genotypes, are shown as blue dots.

**Fig. S12. Validation of MEGAnE genotyping of 25x depth WGS dataset by targeted deep sequencing**

(A) Correlation of MEI genotypes determined by MEGAnE and targeted deep sequencing. Pearson's correlation coefficient between the two techniques are shown. MEIs flagged as filter "PASS" and not-"PASS" are shown as blue and orange bars, respectively. (B) Comparison of allele frequencies of MEIs determined by MEGAnE and deep targeted deep sequencing. Dot color represents ME family. (C) Comparison of correlation of MEI genotypes between MEGAnE and targeted deep sequencing by allele frequency. Dot color represents ME family. (D) Genotyping results of MEGAnE and targeted deep sequencing for all PCR targets. The numbers of individuals with designated genotypes are visualized as heatmaps and with numerical values in the heatmaps. Heatmap color represents ME family.

**Fig. S13. Validation of MEGAnE genotyping result of 15x depth WGS dataset by targeted deep sequencing**

(A) Correlation of MEI genotypes determined by MEGAnE and targeted deep sequencing. Pearson's correlation coefficient between the two techniques are shown. MEIs flagged as filter "PASS" and not-"PASS" are shown as blue and orange bars, respectively. (B) Comparison of allele frequencies of MEIs determined by MEGAnE and targeted deep sequencing. Dot color represents ME family. (C) Comparison of correlation of MEI genotypes between MEGAnE and targeted deep sequencing by allele frequency. Dot color represents ME family. (D) Genotyping results of MEGAnE and targeted deep sequencing for all PCR targets. The numbers of individuals with designated genotypes are visualized as heatmaps and texts in the heatmaps. Heatmap color represents ME family.

**Fig. S14. Validation of MEGAnE genotyping result of joint calling of 25x and 15x depth WGS datasets by targeted sequencing**

(A) Correlation of MEI genotypes determined by MEGAnE and targeted deep sequencing. Pearson's correlation coefficient between the two techniques are shown. MEIs flagged as filter "PASS" and not-"PASS" are shown as blue and orange bars, respectively. (B) Comparison of allele frequencies of MEIs determined by MEGAnE and targeted deep sequencing. Dot color represents ME family. (C) Comparison of correlation of MEI genotypes between MEGAnE and targeted deep sequencing by allele frequency. Dot color represents ME family. (D) Genotyping results of MEGAnE and targeted deep sequencing for all PCR targets. The numbers of individuals with designated genotypes are visualized as heatmaps and texts in the heatmaps. Heatmap color represents ME family.

**Fig. S15. Concordance of genotyping between MEGAnE and deep target-sequencing**
Concordance of genotyping is defined as a proportion of sites which were concordantly genotyped by MEGAnE and deep target-sequencing.

**Fig. S16. Comparison of imputation accuracy based on MEV callsets by MEGAnE and MELT**
We applied MEGAnE and MELTv2 to the 30x WGS of 503 EUR individuals sequenced by the 1000GP. Based on the MEVs detected by MEGAnE and MELT, we performed leave-one-out cross imputation in the 503 EUR (see Supplementary Materials for details). As ground truth, SVs genotyped by PanGenie in the 503 individuals (*3*) were used. (A) Comparison of the numbers of MEVs imputed by the two reference panels containing either the MEV callset from MEGAnE or MELT. Left and right panels show MEIs and ME absences, respectively. The bar plots are stratified by $R^2$ between genotypes from PanGenie and imputation. (B and C) Accuracy of MEV imputation by the reference panel containing MEVs from MEGAnE (B) and MELT (C). Left and right panels show MEIs and ME absences, respectively. To evaluate whether the imputation accuracy is related to MEV allele frequency (x axis), $R^2$ between genotypes from PanGenie and MEGAnE or MELT (y axis) is plotted.

**Fig. S17. Comparison of imputed MEI genotypes and those determined by targeted deep sequencing**
(A) Correlation of the imputed MEI genotypes and those determined by targeted deep sequencing. MEI genotypes were imputed using a reference panel composed of 5,008 haplotypes from the 1000GP dataset. Pearson's correlation coefficient between the two techniques are shown. The color of the bars are separated by $R^2$ of imputed MEIs provided by Minimac4 software. Of note, we used imputed MEIs with an imputation $R^2$ larger than 0.7 for GWAS in this study. (B) Comparison of allele frequencies between the imputed MEIs and deep target-sequencing. Dot color represents ME family. (C) Comparison of correlation of the imputed MEI genotypes and genotypes determined using deep target-sequencing by allele frequency. Dot color represents ME family. (D) The imputed genotypes and those determined by deep target-sequencing for all PCR targets. The numbers of individuals with designated genotypes are visualized as heatmaps and texts in the heatmaps. Heatmap color represents ME family.

**Fig. S18. Computational cost of MEGAnE**
Assessment was performed using the SHIROKANE supercomputer, which is composed of a Intel Xeon Gold 6154 (3.0 GHz) processor, a Lustre parallel distributed file system, and Altair Grid Engine. (A) Computational cost required to analyze one WGS file by MEGAnE step 1. Here, we used a 35x WGS of an individual, NA12878, in the 1000GP. The WGS reads are mapped on GRCh38 and stored in CRAM format. (B) Computational cost required to analyze 2,504 WGS

datasets. First, the human reference genome build, GRCh38, was processed by MEGAnE step 0. Then, MEGAnE step 1 was applied to the 2,504 WGS datasets sequenced at 30x depth by the 1000GP. In this step, 200 threads were allocated for analysis. After MEGAnE step 1, results from step 1 were merged and a joint callset was generated by MEGAnE step 2. (C and D) Wall time (C) and maximum RAM usage (D) required for analysis of each WGS of the 2,504 individuals. Two threads were allocated for analysis of each WGS dataset.

**Fig. S19. Number of MEVs discovered from the 1000GP dataset**
(A) Number of detected MEVs per individual. (B) Number of detected MEVs per individual by the three ME familes. MEGAnE was applied to the 2,504 30x WGS in the 1000GP, and a joint callset was generated. There is one outlier point in AFR, NA20314, an individual reported to have no recent African ancestry (*44*).

**Fig. S20. Comparison of MEV discovery in 1000GP JPT and BBJ**
(A) Venn diagram showing overlap between singletons found in JPT within the 1000GP (104 individuals) and all MEVs, including the filter flag non-"PASS" variants, found in BBJ (4,880 individuals). More than half of the singletons found in JPT were also found in BBJ. (B) Venn diagram showing overlap between JPT-specific non-singleton MEVs found in the 1000GP and all MEVs, including the filter flag non-"PASS" variants, found in BBJ. The vast majority of JPT-specific non-singletons found in 1000GP were also found in BBJ.

**Fig. S21. MEV discovery from the SFARI**
(A) Cumulative plot showing the number and allele frequency of MEVs found in the SFARI. (B) Distribution of the first 2 principal components of MEVs found in the SFARI. Red rectangle shows the 7,642 subjects with PC-inferred European ancestry. The 7,642 subjects were used to compare the insertion preference of MEVs between Europeans and Japanese (BBJ). Kernel density of the distribution is overlaid.

**Fig. S22. Thirteen *de novo* L1HS insertions found in SFARI cohort**
IGV genome browser views showing 13 *de novo* L1 insertion sites. The positions of TSD are shown at the top of each set of images. Chimerically mapped reads are highlighted with non-gray colors. The presence of chimeric reads only in children supports the inference that these are *de novo* insertions.

**Fig. S23. Eighteen examples of *de novo Alu* insertions found in SFARI cohort**
IGV genome browser views showing 18 *de novo Alu* insertion sites. The positions of TSD are shown at the top of each set of images. Chimerically mapped reads are highlighted with non-gray colors. The presence of chimeric reads only in children supports the inference that these are *de novo* insertions.

**Fig. S24. Three *de novo* L1HS insertions shared between two children**
IGV genome browser views showing 3 *de novo* L1HS insertion sites. The positions of TSD are shown at the top of each set of images. Chimerically mapped reads are highlighted with non-gray

colors. The presence of chimeric reads in two children but neither parents supports the inference that these *de novo* insertions are present in a mosaic manner in one of the parents.

**Fig. S25. Three *de novo Alu* insertions shared between two children**
IGV genome browser views showing 3 *de novo Alu* insertion sites. The positions of TSD are shown at the top of each set of images. Chimerically mapped reads are highlighted with non-gray colors. The presence of chimeric reads in two children but not in parents supports the inference that these *de novo* insertions are present in a mosaic manner in one of the parents.

**Fig. S26. Non-reference MER41A insertion presumably originating from a partial deletion of MER41A in the reference allele**
(A) UCSC genome browser view showing the position of a non-reference MER41A insertion (chr13:25878464). This insertion is nested in another copy of MER41A. (B) Dot matrix showing nucleotide similarities between two PacBio reads from an individual, HG00733, and the consensus sequence of MER41A. HG00733 carries heterozygous insertion of MER41A. Top panel shows the PacBio reads coming from the reference allele, while the bottom panel shows the reads from the alternative allele carrying the MER41A insertion. MER41A in the alternative allele is an almost full-length copy, while the reference allele has a partial deletion. (C) Illustration showing the predicted mechanism of the non-reference MER41A insertion. It is estimated that the polymorphic partial deletion of MER41A is present in the reference allele. However, when a polymorphic partial ME deletion is represented by the reference allele, the deletion can sometimes be detected as an insertion in the alternative allele by MEGAnE.

**Fig. S27. *De novo* THE1_I, a MaLR-like endogenous retrovirus element, insertion found in a trio**
(A) IGV browser view showing the *de novo* THE1_I insertion site in a child of a trio in SFARI. Chimeric reads, shown by non-gray colors, are only found in the child. In addition, there is a read depth difference at the TSD. Thus only the child carries the insertion, i.e. it is a *de novo* insertion. (B) Sequences found at the THE1_I insertion breakpoints. The insertion is flanked by an 8-bp TSD. The two breakpoint-spanning reads overlap each other, allowing us to reconstruct the 149-bp inserted sequence, shown by blue characters. (C) Illustration showing the region of THE1_I inserted into the child's genome. The insertion was not a full-length insertion, but only a 149-bp fragment of THE1_I. (D) Alignment between the sequence of the *de novo* THE1_I insertion and the consensus sequence of THE1_I. The *de novo* THE1_I insertion has high similarity to the consensus sequence. (E) Alignment between the sequence of the *de novo* THE1_I insertion and a part of the chromosome 16 of GRCh38. The *de novo* THE1_I insertion has perfect similarity only to this region of GRCh38, suggesting that the origin of the *de novo* insertion is the sequence of chromosome 16.

**Fig. S28. Non-reference LTR8A insertion supported by long-reads**
(A) UCSC genome browser view showing the position of a non-reference LTR8A insertion (GRCh38:chr17:41144157). The insertion is not nested in another LTR8 copy. (B) UCSC genome browser view showing a region of marmoset genome (calJac4) corresponding to the LTR8

polymorphism in the human genome. The LTR8A copy is present in marmoset and macaque genomes, however it is absent in the reference human genome. (C and D) IGV genome browser view showing PacBio long reads (C) and Illumina short-reads (D) from an individual, HG00514, mapping to the LTR8A insertion site. HG00514 carries homozygous LTR8A insertion. Both long and short-reads were mapped chimerically (reads with non-gray colors), showing the presence of insertion. (E) Dot matrix showing the nucleotide similarities between an PacBio read from HG00514 and the human reference genome (top panel) and the consensus sequence of LTR8A (bottom panel). The PacBio read provides evidence of an inserted LTR8A sequence which is not present in the human reference genome, GRCh38.

**Fig. S29. Examples of probable novel LoF ME insertions**
(A and C) UCSC genome browser views showing *Alu* (A) and L1 (C) insertion site in CDS of *ADGRE2* (A) and *DLEC1* (C), respectively. (B and D) IGV genome browser views showing chimerically mapped reads, which are highlighted by non-gray colors, at the insertion breakpoints. (B) The insertions were found in two AFR individuals, NA19312 (LWK) and NA19921 (ASW). (D) The insertion was found in 4 Japanese individuals in BBJ. Here, we show the mapping pattern of one individual.

**Fig. S30. Principal component analysis using SNVs and MEVs**
(A) Pearson correlation coefficients between principal components (PCs) of SNVs and MEVs. (B) Comparison between PCs of SNVs and MEVs. PCs of SNVs and MEs that have absolute Pearson correlation coefficients larger than 0.5 were used for analysis.

**Fig. S31. Super-population- and population-specific MEIs found in the 1000GP dataset**
(A and B) Proportion of the ME families among super-population- and population-specific MEIs found in the 1000GP dataset. MEVs were identified from 2,504 individuals in the 1000GP dataset, and the proportions of each ME family are shown as pie charts.

**Fig. S32. Insertion distribution of singletons and family-specific heritable insertions**
(A and B) Correlations between genome features and singletons found in 1000GP GRCh38 dataset, BBJ, and SFARI (A) or family-specific heritable insertions. Spearman correlation coefficients between the number of super-population-specific ME insertions in non-overlapping 1Mb windows and genome features of H1-hESC cells were calculated. Dendrograms show results of hierarchical clustering.

**Fig. S33. Correlation between super-population-specific *Alu* insertion and genome features**
(A) Correlations between super-population-specific *Alu* insertions and genome features. MEVs were identified from the 1000GP dataset. Spearman correlation coefficients between the number of super-population-specific *Alu* insertions in non-overlapping 1Mb windows and genome features of H1-hESC cells were calculated. (B) Distribution of the super-population-specific *Alu* polymorphisms and replication timing. The counts of super-population-specific *Alu* insertions and replication timing of non-overlapping 5Mb windows are shown. (C) Correlation between super-

population-specific *Alu* insertions and CpG methylation levels. Spearman correlation coefficients between the number of super-population-specific *Alu* insertions and the CpG methylation levels of non-overlapping 1Mb windows were calculated. The CpG methylation levels measured in multiple iPSCs and ESCs established from Japanese and Caucasian individuals were used.

**Fig. S34. Motifs detected at ME insertion breakpoints**
To detect motifs characteristic at ME insertion breakpoints, 4bp upstream and 10bp downstream of ME insertion breakpoints found in the 1000GP, SFARI, and BBJ datasets are used. Enrichment of motifs compared to the random sequences of the human genome build, GRCh38, was tested by HOMER software. "Possible false positive" flags for HERV-K motifs reflect the output from HOMER.

**Fig. S35. Preferential insertion of MEs in non-transcribed strand of genes**
(A) The number of family-specific inherited MEIs found in genes. MEIs of all three active ME families are enriched in the non-transcribed strand of genes. *P* value of binomial test between the number of MEIs in transcribed and non-transcribed strands of genes are shown at the top of the panels. (B) Enrichment of family-specific inherited *Alu* and L1HS in genes expressed in spermatids. (C) Enrichment of *de novo Alu* in genes expressed in spermatids. (B and C) Wilcoxson rank-sum statistics of the expressions of the genes with MEIs compared to those of the genes without MEI are shown. SPG: spermatogonia, SPC: spermatocyte, ST: spermatids.

**Fig. S36. Tissue-sharing and distribution of ME-eQTLs**
(A) Histogram of the numbers of tissues in which a given ME-eQTL was detected. (B) Proportion of tissue-specific and multi-tissue ME-eQTLs. The numbers shown in the bars represent the actual number of ME-eQTLs used for visualization. (C) Distribution of ME-eQTLs across chromosomes. If an ME-eQTL is detected in multiple tissues, mean effect size is used for visualization.

**Fig. S37. Genome features associated with ME-eQTLs**
(A) Raw number of ME-eQTLs (left) and significant (LFSR $< 0.05$) ME-eGene pairs (right) detected in early-replicating domains. *P* values of Fisher exact tests are shown. (B) Odds ratios of being detected as a significant (LFSR $< 0.05$) ME-eGene pair in a genome region with the designated features. Red and blue points are significant enrichments or depletions, having odds ratios significantly different from zero ($P < 0.05$ after Bonferroni correction). Error bars show 95% confidence intervals. RT, replication timing.

**Fig. S38. Three examples of MEIs in gene regulatory elements**
UCSC genome browser view showing gene models and epigenetic states near three *Alu* insertions. All the three *Alu* insertions are detected as ME-eQTLs associating with a decrease in expression of an adjacent gene. The insertion sites of *Alu* are shown as vertical red lines. The eGenes associated with the *Alu* variants are highlighted with arrows at the top of the plots.

**Fig. S39. Weak association between an *Alu* insertion in the 3'UTR of *EGFR* gene and asthma**

Regional association plot showing the weak association between the *Alu* insertion shown as a purple plus mark and asthma. MEVs and SNVs are shown as plus marks and circles, respectively. The *Alu* insertion is highlighted with a red arrow.

**Fig. S40. Phenograms showing linkage between MEVs associated with traits**
(A) LD ($R^2 > 0.8$) between 43 MEVs and sentinel variants in 73 GWASs in Pan-UKB. (B) LD ($R^2 > 0.8$) between 42 MEVs and sentinel variants in 54 GWASs in BBJ.

**Fig. S41. PCR validation of L1-NEDD4 allele frequency in Japanese and CRISPR-Cas9 knockout in iPSCs**
(A) L1-NEDD4 was genotyped in 70 iPSCs established from healthy Japanese individuals. (B) Nine L1-NEDD4 KO and 11 WT iPSC clones. The sgRNA-Cas9 complex and HDR template were transfected to iPSCs derived from a subject carrying two copies of L1-NEDD4. The KO and WT clones were obtained by limiting dilution.

**Fig. S42. Conditioning of NEDD4-eQTL and exon-eQTL analysis**
Regional association plot showing the three haplotype blocks independently associated with *NEDD4* gene expression. MEVs and SNVs are shown as plus marks and circles, respectively. The L1-NEDD4 is highlighted with a red arrow. (B) Heatmap showing the effect size of L1-NEDD4 for expression of three exons in 49 tissues. Color bar at the top of the heatmap corresponds to tissue. Significant associations (FDR < 0.05) are marked with asterisks. L1-NEDD4 associates with increased expression of exon 9 and 34 in fibroblasts.

**Fig. S43. T2D and prostate cancer GWAS detect associations with MEVs**
Manhattan plots and regional association plots showing the results of T2D and prostate cancer GWAS. MEVs and SNVs are shown as plus marks and circles, respectively. MEVs are highlighted with red arrows. (A) An *Alu* insertion associated with T2D. (B) An *Alu* insertion associated with prostate cancer. The *Alu* is tagged with a sentinel variant detected after conditioning with the two SNVs (8:128103969:C:T and 8:128047278:G:C) serving as sentinels for the two most associated haplotypes.

**Fig. S44. Odds ratios of carrying L1-NEDD4 by body part affected by keloid**
Red and dark red points show odds ratios significantly greater than one ($P < 0.05$ after Bonferroni correction accounting for additional tests shown in the main figure).

### Fig. S1

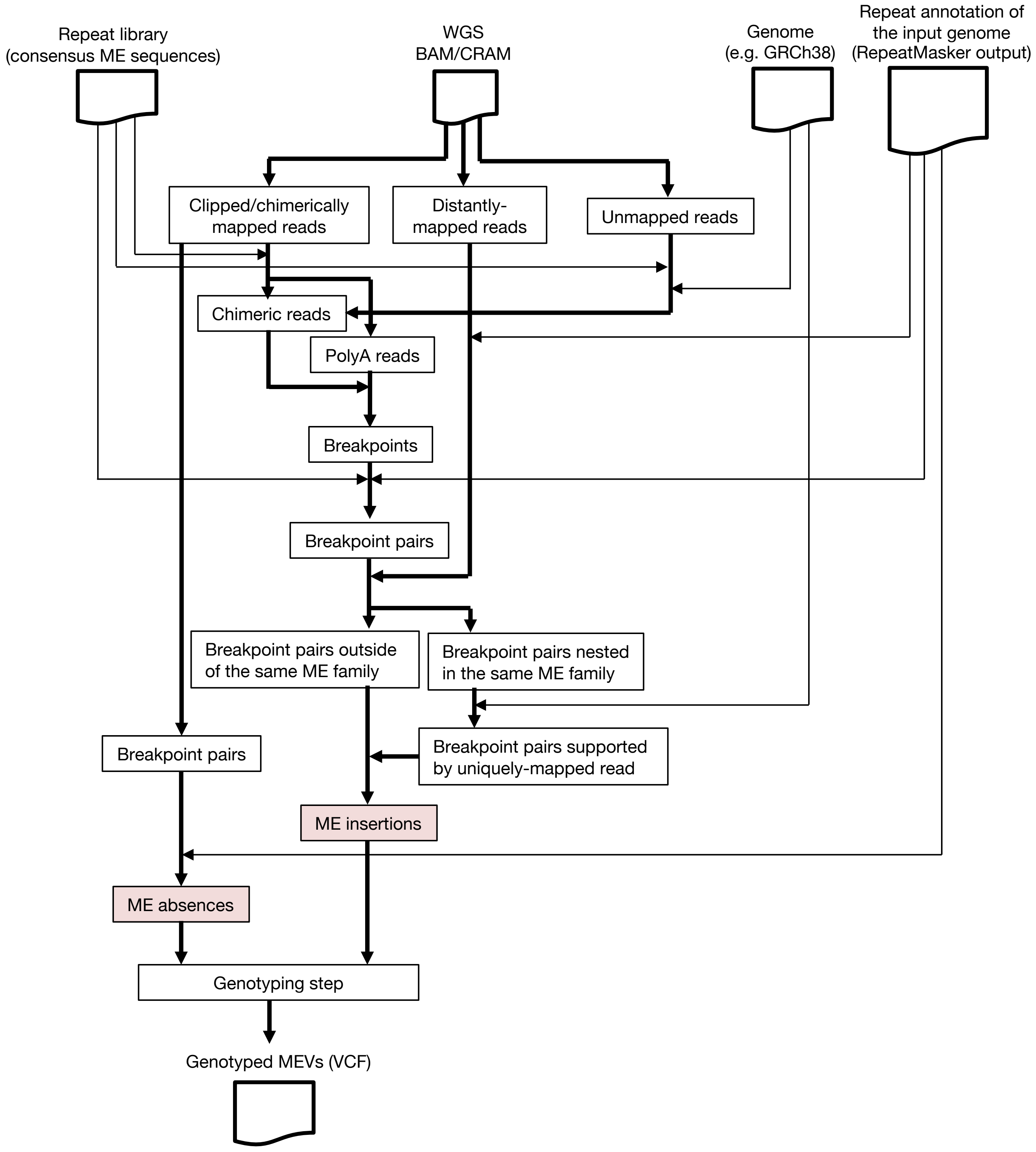

Repeat library
(consensus ME sequences)
WGS
BAM/CRAM
Genome
(e.g. GRCh38)
Repeat annotation of
the input genome
(RepeatMasker output)
Clipped/chimerically
mapped reads
Distantly-
mapped reads
Unmapped reads
Chimeric reads
PolyA reads
Breakpoints
Breakpoint pairs
Breakpoint pairs outside
of the same ME family
Breakpoint pairs nested
in the same ME family
Breakpoint pairs
Breakpoint pairs supported
by uniquely-mapped read
ME insertions
ME absences
Genotyping step
Genotyped MEVs (VCF)

### Fig. S2

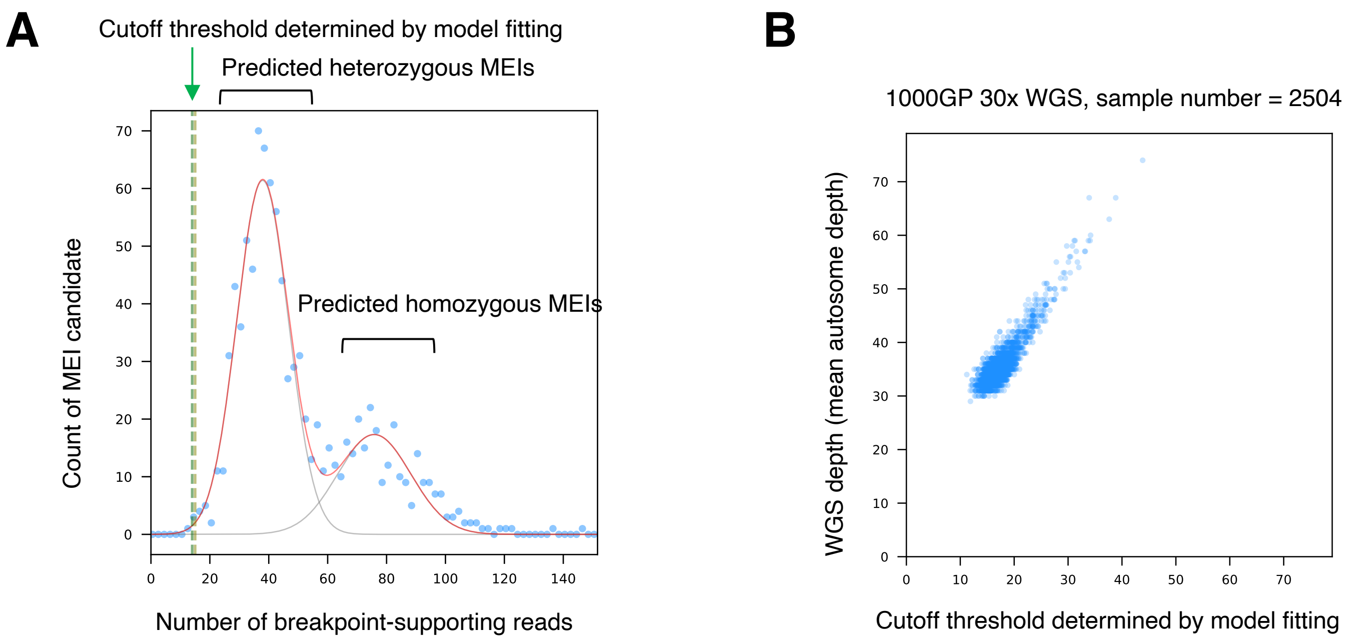
A
Cutoff threshold determined by model fitting
Predicted heterozygous MEIs
Predicted homozygous MEIs
Count of MEI candidate
Number of breakpoint-supporting reads
B
1000GP 30x WGS, sample number = 2504
WGS depth (mean autosome depth)
Cutoff threshold determined by model fitting

### Fig. S3

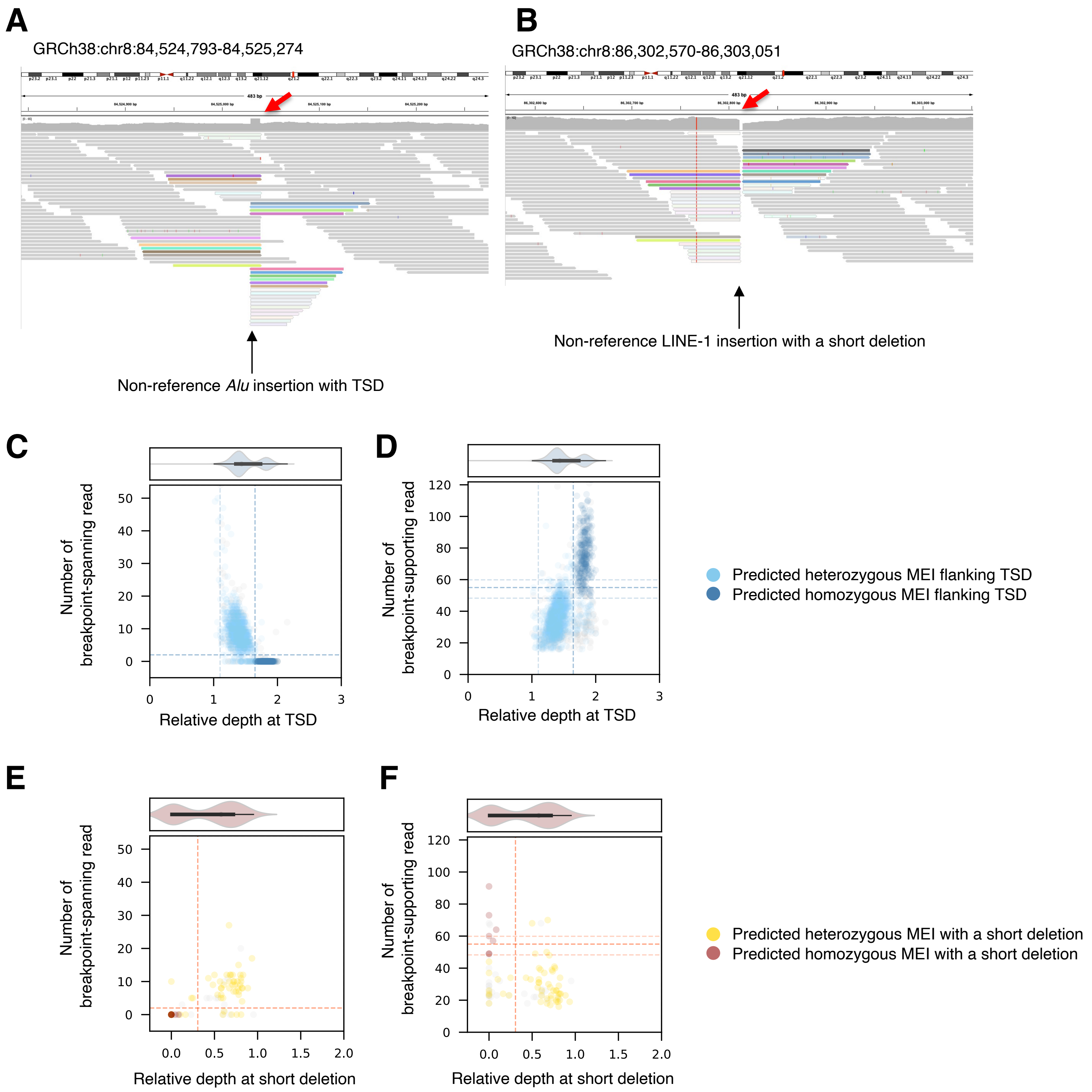
A
GRCh38:chr8:84,524,793-84,525,274
Non-reference Alu insertion with TSD
B
GRCh38:chr8:86,302,570-86,303,051
Non-reference LINE-1 insertion with a short deletion
C
Number of breakpoint-spanning read
Relative depth at TSD
D
Number of breakpoint-supporting read
Relative depth at TSD
Predicted heterozygous MEI flanking TSD
Predicted homozygous MEI flanking TSD
E
Number of breakpoint-spanning read
Relative depth at short deletion
F
Number of breakpoint-supporting read
Relative depth at short deletion
Predicted heterozygous MEI with a short deletion
Predicted homozygous MEI with a short deletion

### Fig. S4

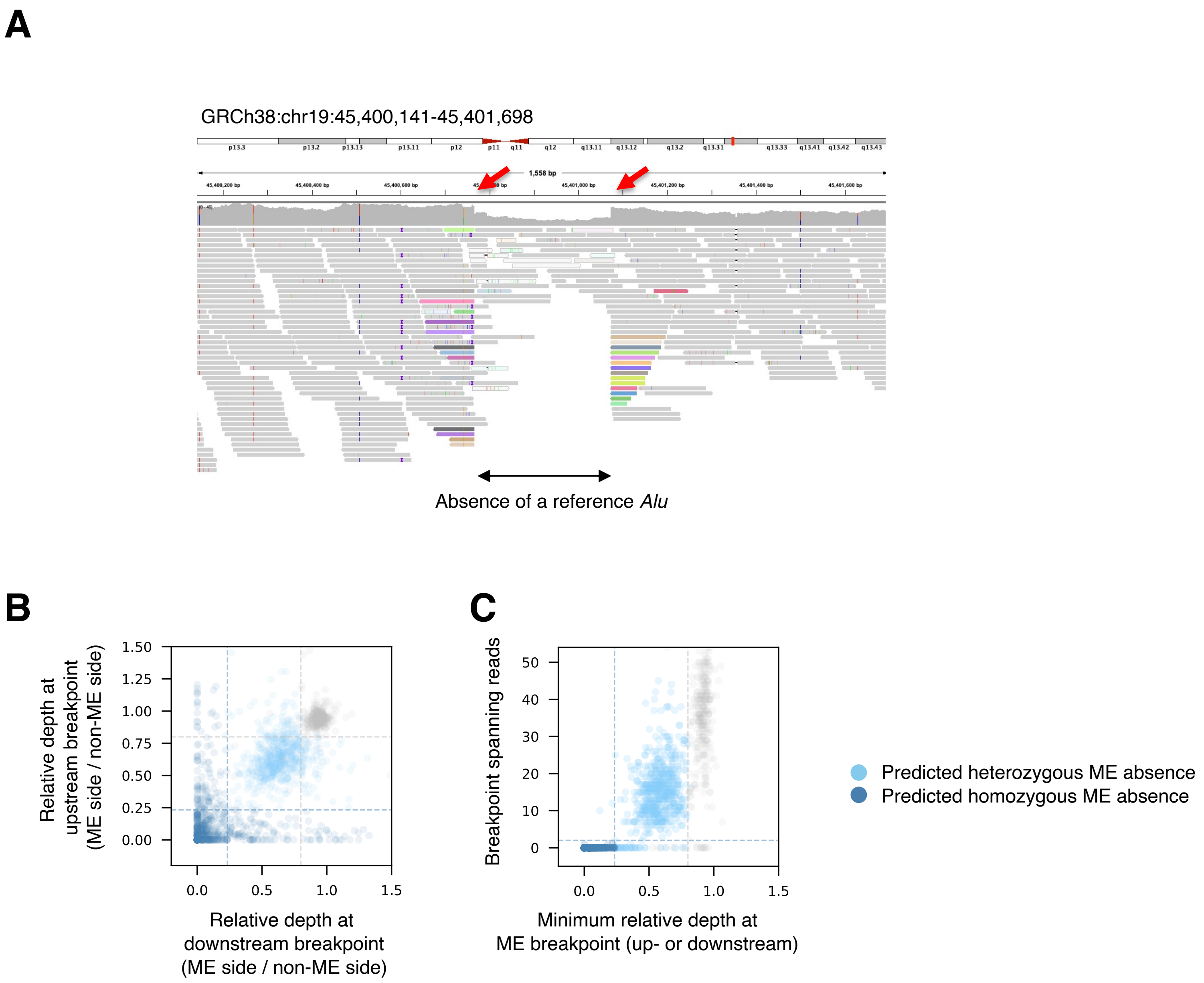
A
GRCh38:chr19:45,400,141-45,401,698
1,558 bp
Absence of a reference *Alu*
B
Relative depth at upstream breakpoint (ME side / non-ME side)
1.50
1.25
1.00
0.75
0.50
0.25
0.00
0.0
0.5
1.0
1.5
Relative depth at downstream breakpoint (ME side / non-ME side)
C
Breakpoint spanning reads
50
40
30
20
10
0
0.0
0.5
1.0
1.5
Minimum relative depth at ME breakpoint (up- or downstream)
Predicted heterozygous ME absence
Predicted homozygous ME absence

### Fig. S5

**A**

| Polymorphism type | Number of MEVs identified by PAV | Number of MEGAnE calls (filter PASS) | Number of true positive MEGAnE calls (filter PASS) | True positive rate |
|---|---|---|---|---|
| MEI | 14691 | 7599 | 7367 | 97% |
| ME absence | 9831 | 2131 | 1997 | 94% |

**B**

| Polymorphism type | Number of MEVs identified by PAV | Number of MEGAnE calls (filter not-PASS) | Number of true positive MEGAnE calls (filter not-PASS) | True positive rate |
|---|---|---|---|---|
| MEI | 14691 | 3198 | 1016 | 32% |
| ME absence | 9831 | 164 | 42 | 26% |

**C**

| MEI location | Number of MEVs identified by PAV | Number of MEIs detected by MEGAnE (%total) | Number of MEIs missed by MEGAnE |
|---|---|---|---|
| Nested in another ME belonging to the same ME family | 5255 | 1067 (20%) | 4158 |
| Nested in another ME belonging to different ME family | 4407 | 3536 (80%) | 871 |
| Nested in simple repeat | 198 | 66 (33%) | 132 |
| None of above | 4831 | 3714 (77%) | 1117 |
| Total | 14691 | 8383 (57%) | 6308 |

**D**

| MEI family | Number of MEIs missed by MEGAnE | Recently active ME subfamily (Number of missed by MEGAnE) |
|---|---|---|
| *Alu* | 420 | *Alu*Y (n = 235) |
| LINE-1 | 423 | L1HS (n = 61) |
| SVA | 66 | SVA_E (n = 15); SVA_F (n = 15) |
| ERV | 139 | LTR5_Hs (n = 7) |
| DNA transposon | 69 | N.A. |

**E**

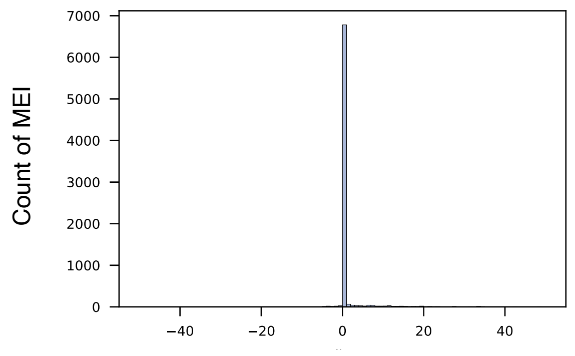

Relative MEI breakpoint position
(Distance between positions determined by PAV and MEGAnE)

### Fig. S6

**A**

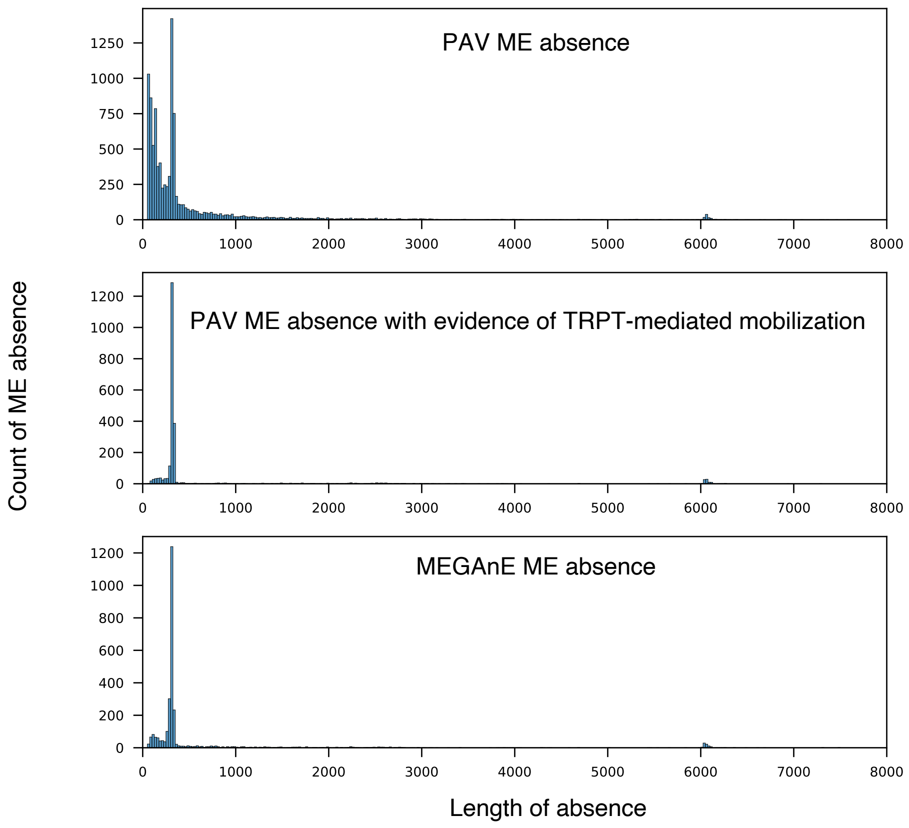

**B**

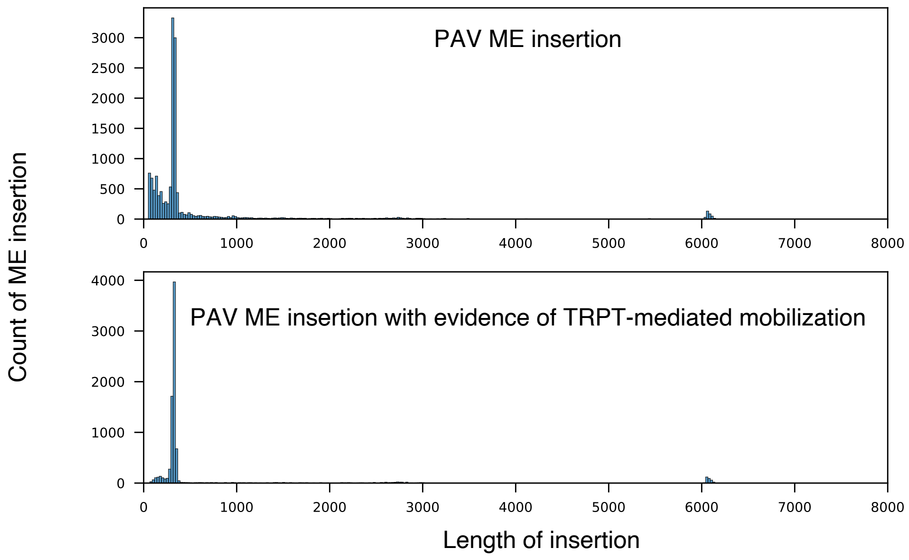

**C**

| Polymorphism type | Number of MEVs identified by PAV | Number of MEVs identified by PAV which have evidence of TPRT-mediated insertion (TPRT-variants) | Number of TPRT-variants detected by MEGAnE (Filter "PASS" variants) | Recovery of TPRT-variants by MEGAnE |
|---|---|---|---|---|
| MEI | 14691 | 8362 | 6829 | 81.7% |
| ME absence | 9831 | 2210 | 1769 | 80.0% |

**D**

| Polymorphism type | Number of MEVs identified by PAV | Number of MEVs identified by PAV which have evidence of TPRT-mediated insertion (TPRT-variants) | Number of TPRT-variants detected by MEGAnE (Filter not-"PASS" variants) | Recovery of TPRT-variants by MEGAnE |
|---|---|---|---|---|
| MEI | 14691 | 8362 | 708 | 7.3% |
| ME absence | 9831 | 2210 | 67 | 3.1% |

### Fig. S7

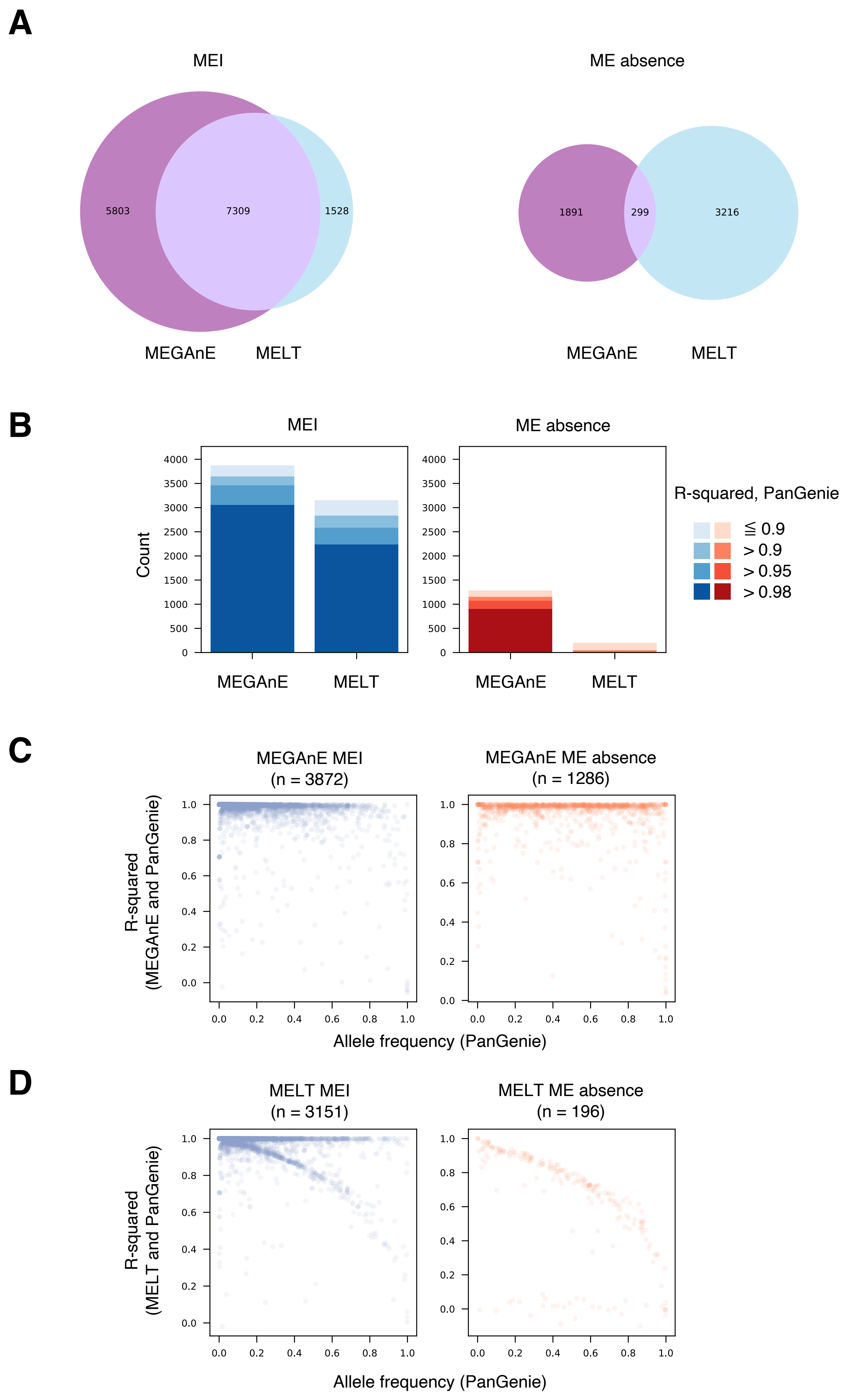
A
MEI
5803
7309
1528
MEGAnE
MELT
ME absence
1891
299
3216
MEGAnE
MELT
B
MEI
ME absence
Count
MEGAnE
MELT
MEGAnE
MELT
R-squared, PanGenie
≦ 0.9
> 0.9
> 0.95
> 0.98
C
MEGAnE MEI
(n = 3872)
MEGAnE ME absence
(n = 1286)
R-squared
(MEGAnE and PanGenie)
Allele frequency (PanGenie)
D
MELT MEI
(n = 3151)
MELT ME absence
(n = 196)
R-squared
(MELT and PanGenie)
Allele frequency (PanGenie)

### Fig. S8

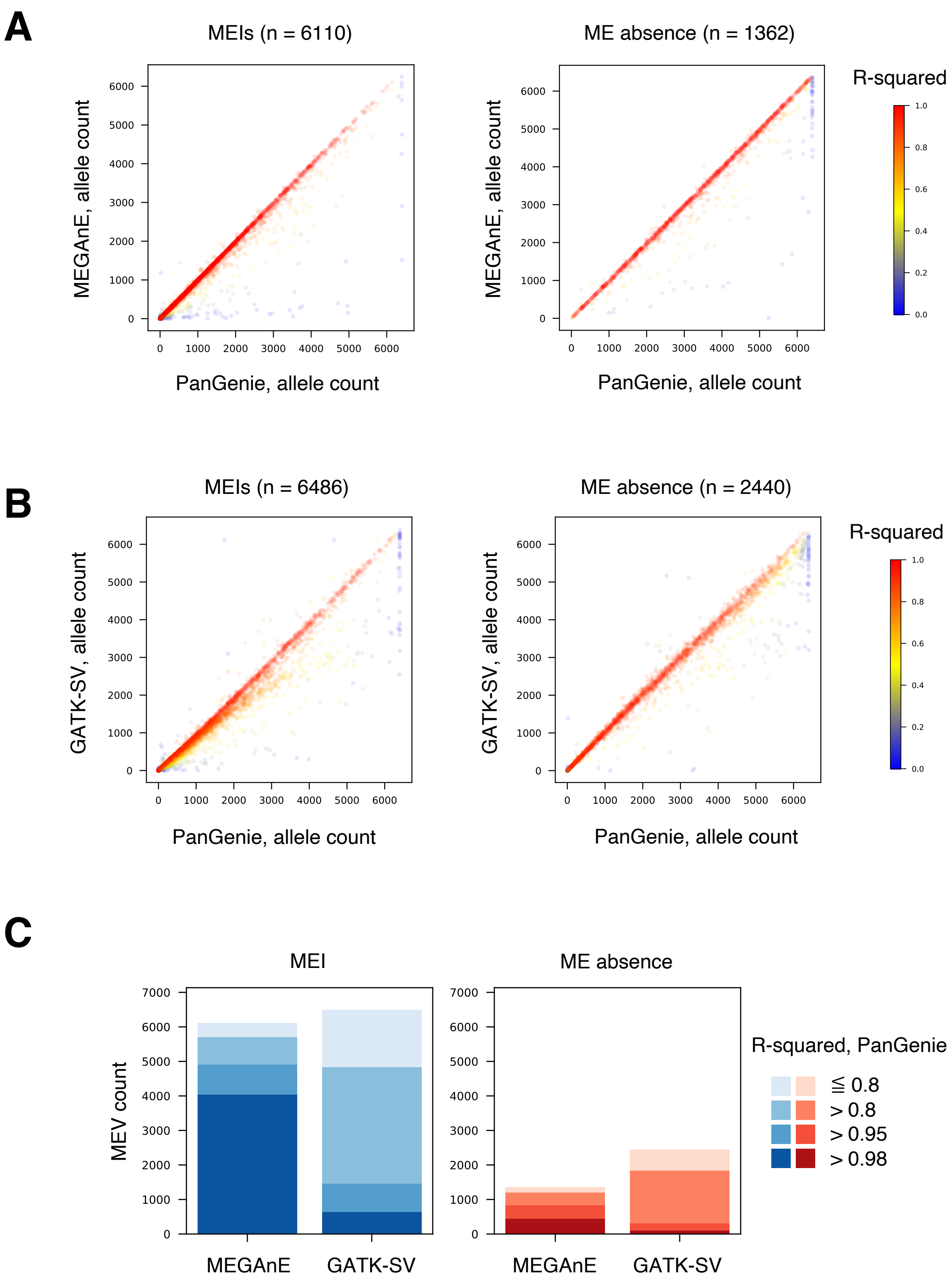
A
MEIs (n = 6110)
ME absence (n = 1362)
MEGAnE, allele count
PanGenie, allele count
R-squared
B
MEIs (n = 6486)
ME absence (n = 2440)
GATK-SV, allele count
PanGenie, allele count
R-squared
C
MEI
ME absence
MEV count
MEGAnE
GATK-SV
R-squared, PanGenie
≦ 0.8
> 0.8
> 0.95
> 0.98

### Fig. S9

## A

| Mean MEI count per individual | Total MEr | MEr | | | | |
|---|---|---|---|---|---|---|
| | | Child = 0<br>Father = 2<br>Mother = 2 | Child = 1<br>Father = 0<br>Mother = 0<br>(Family-specific) | Child = 1<br>Father = 0<br>Mother = 0<br>(Observed in >1 family) | Child = 1<br>Father = 2<br>Mother = 2 | Child = 2<br>Father = 0<br>Mother = 0 |
| 1416 | 1.75% | 0.25% | 0.03% | 1.02% | 0.46% | 0.24% |

## B

| Mean MEI count per individual | Mean family-specific MEI per individual | % family-specific MEI with MEr | % family-specific MEI concordant with Mendelian inheritance |
|---|---|---|---|
| 1416 | 10.4 | 4.7% | 95.3% |

### Fig. S10

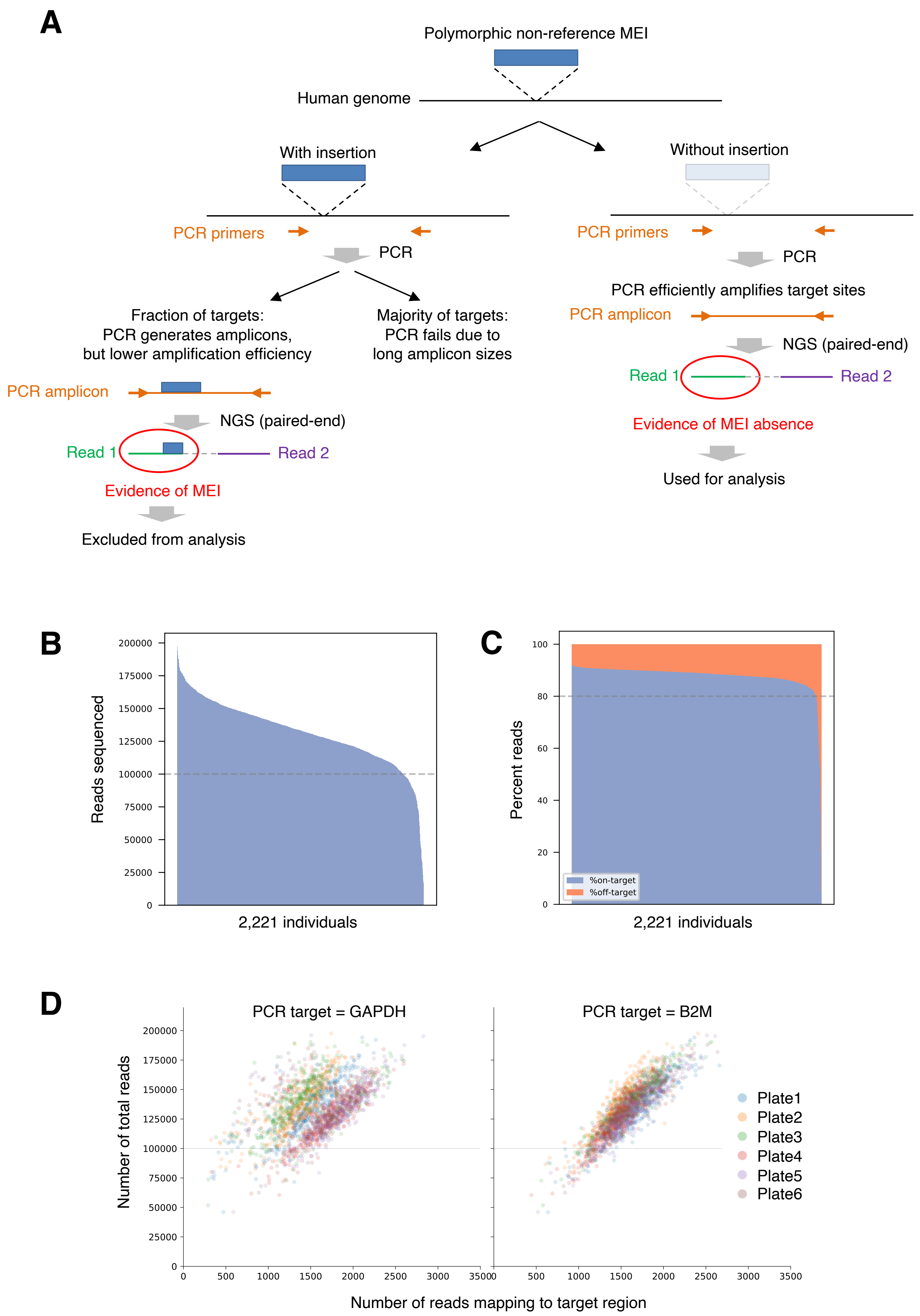

A
Polymorphic non-reference MEI
Human genome
With insertion
Without insertion
PCR primers
PCR
Fraction of targets:
PCR generates amplicons,
but lower amplification efficiency
Majority of targets:
PCR fails due to
long amplicon sizes
PCR amplicon
NGS (paired-end)
Read 1
Read 2
Evidence of MEI
Excluded from analysis
PCR primers
PCR
PCR efficiently amplifies target sites
PCR amplicon
NGS (paired-end)
Read 1
Read 2
Evidence of MEI absence
Used for analysis
B
Reads sequenced
2,221 individuals
C
Percent reads
%on-target
%off-target
2,221 individuals
D
PCR target = GAPDH
PCR target = B2M
Number of total reads
Plate1
Plate2
Plate3
Plate4
Plate5
Plate6
Number of reads mapping to target region

### Fig. S11

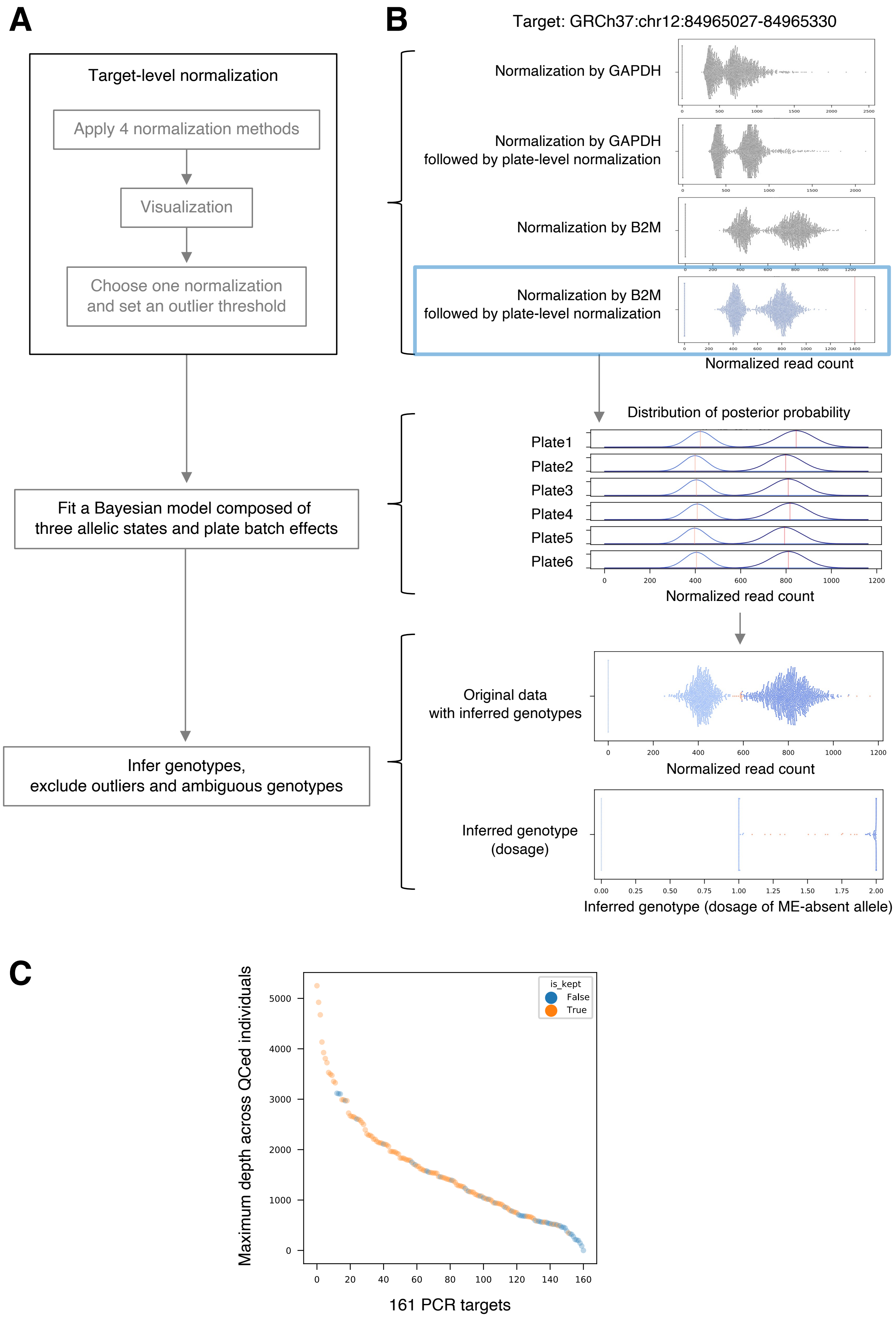
A
Target-level normalization
Apply 4 normalization methods
Visualization
Choose one normalization and set an outlier threshold
Fit a Bayesian model composed of three allelic states and plate batch effects
Infer genotypes, exclude outliers and ambiguous genotypes
B
Target: GRCh37:chr12:84965027-84965330
Normalization by GAPDH
Normalization by GAPDH followed by plate-level normalization
Normalization by B2M
Normalization by B2M followed by plate-level normalization
Normalized read count
Distribution of posterior probability
Plate1
Plate2
Plate3
Plate4
Plate5
Plate6
Normalized read count
Original data with inferred genotypes
Normalized read count
Inferred genotype (dosage)
Inferred genotype (dosage of ME-absent allele)
C
Maximum depth across QCed individuals
is_kept
False
True
161 PCR targets

### Fig. S12

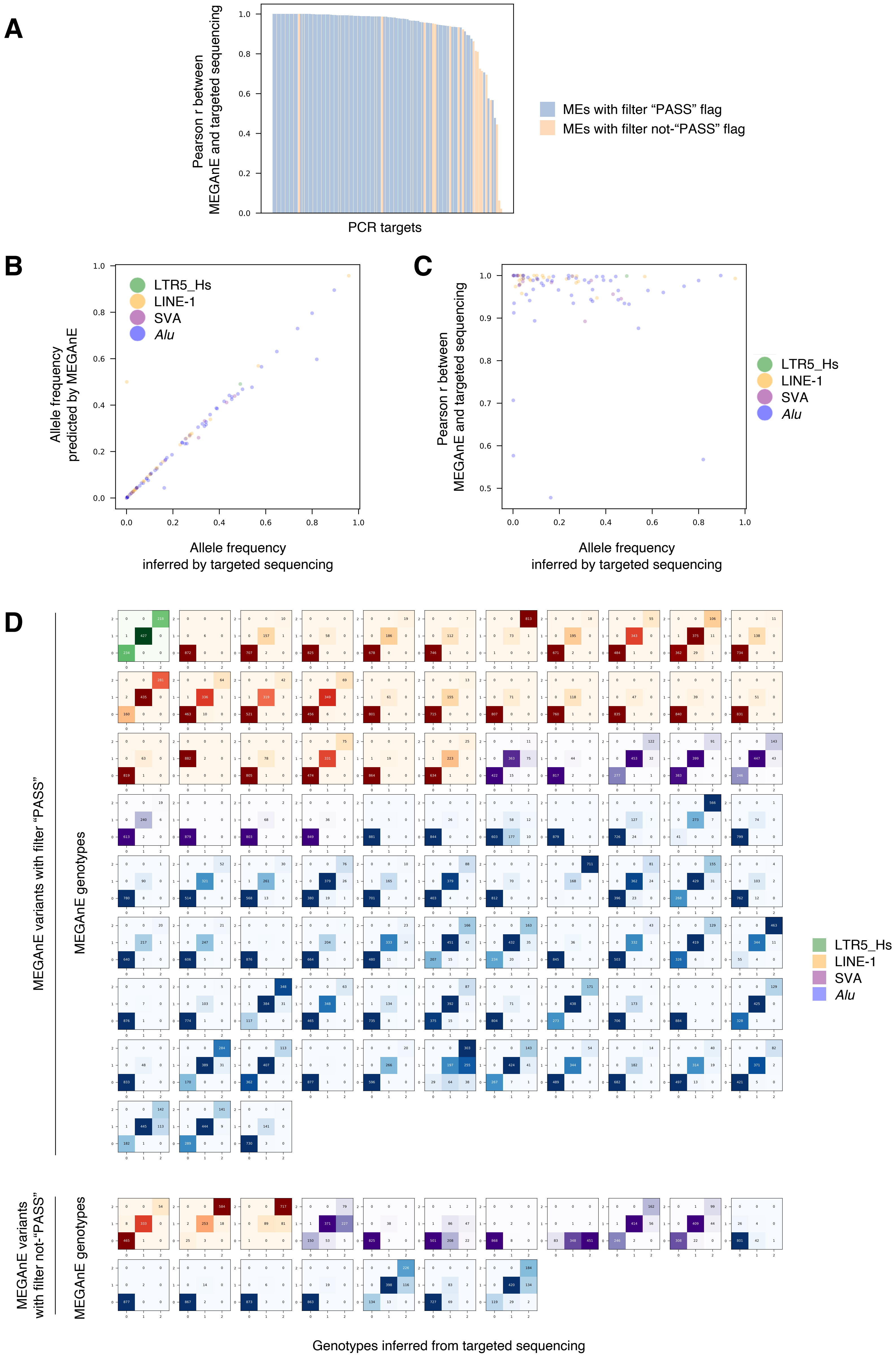
A
Pearson r between MEGAnE and targeted sequencing
PCR targets
MEs with filter “PASS” flag
MEs with filter not-“PASS” flag
B
Allele frequency predicted by MEGAnE
Allele frequency inferred by targeted sequencing
LTR5_Hs
LINE-1
SVA
Alu
C
Pearson r between MEGAnE and targeted sequencing
Allele frequency inferred by targeted sequencing
LTR5_Hs
LINE-1
SVA
Alu
D
MEGAnE variants with filter “PASS”
MEGAnE genotypes
MEGAnE variants with filter not-“PASS”
MEGAnE genotypes
Genotypes inferred from targeted sequencing
LTR5_Hs
LINE-1
SVA
Alu

### Fig. S13

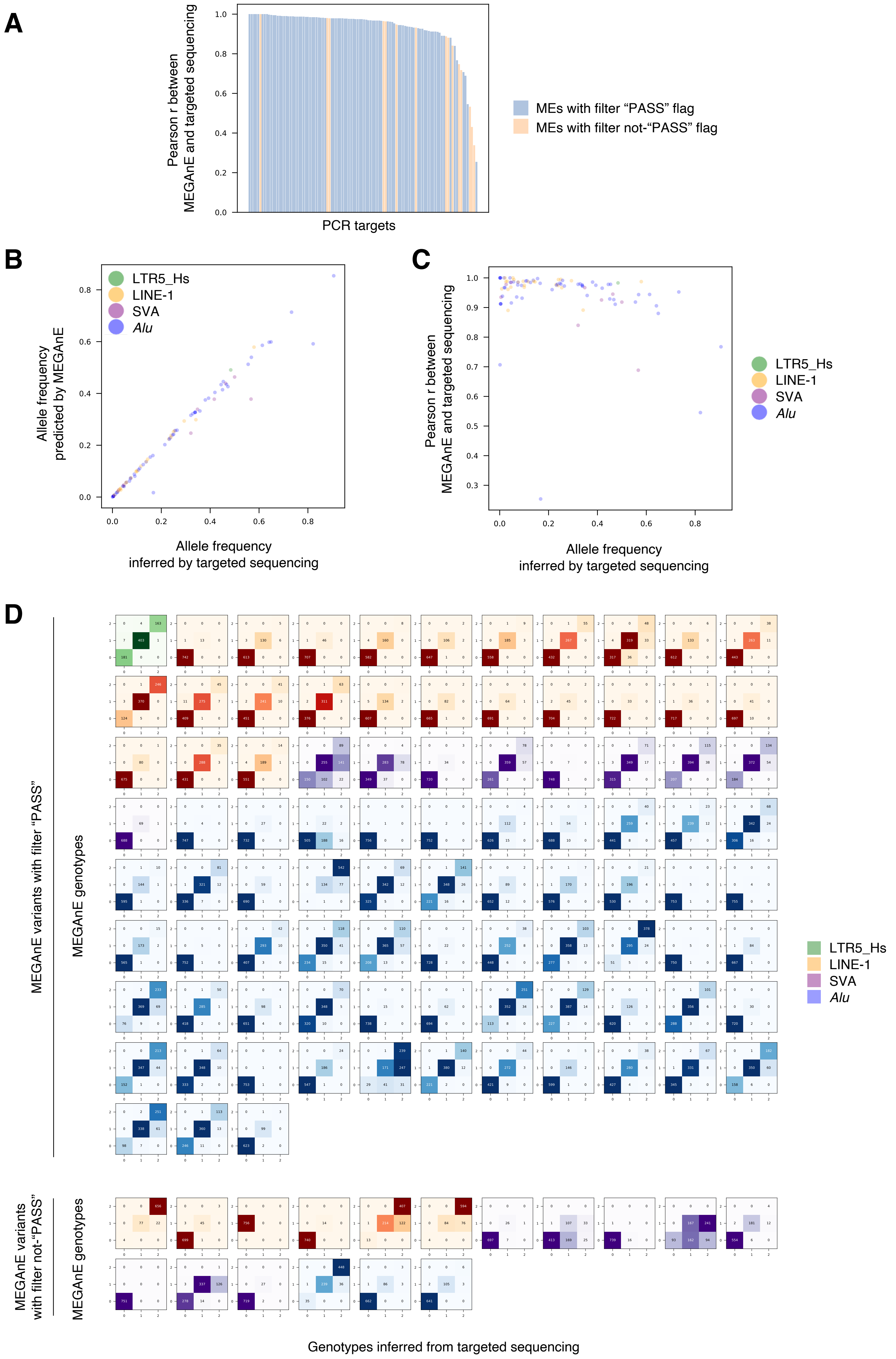
A
Pearson r between
MEGAnE and targeted sequencing
PCR targets
MEs with filter "PASS" flag
MEs with filter not-"PASS" flag
B
Allele frequency
predicted by MEGAnE
Allele frequency
inferred by targeted sequencing
LTR5_Hs
LINE-1
SVA
Alu
C
Pearson r between
MEGAnE and targeted sequencing
Allele frequency
inferred by targeted sequencing
LTR5_Hs
LINE-1
SVA
Alu
D
MEGAnE variants with filter "PASS"
MEGAnE genotypes
MEGAnE variants
with filter not-"PASS"
MEGAnE genotypes
Genotypes inferred from targeted sequencing
LTR5_Hs
LINE-1
SVA
Alu

### Fig. S14

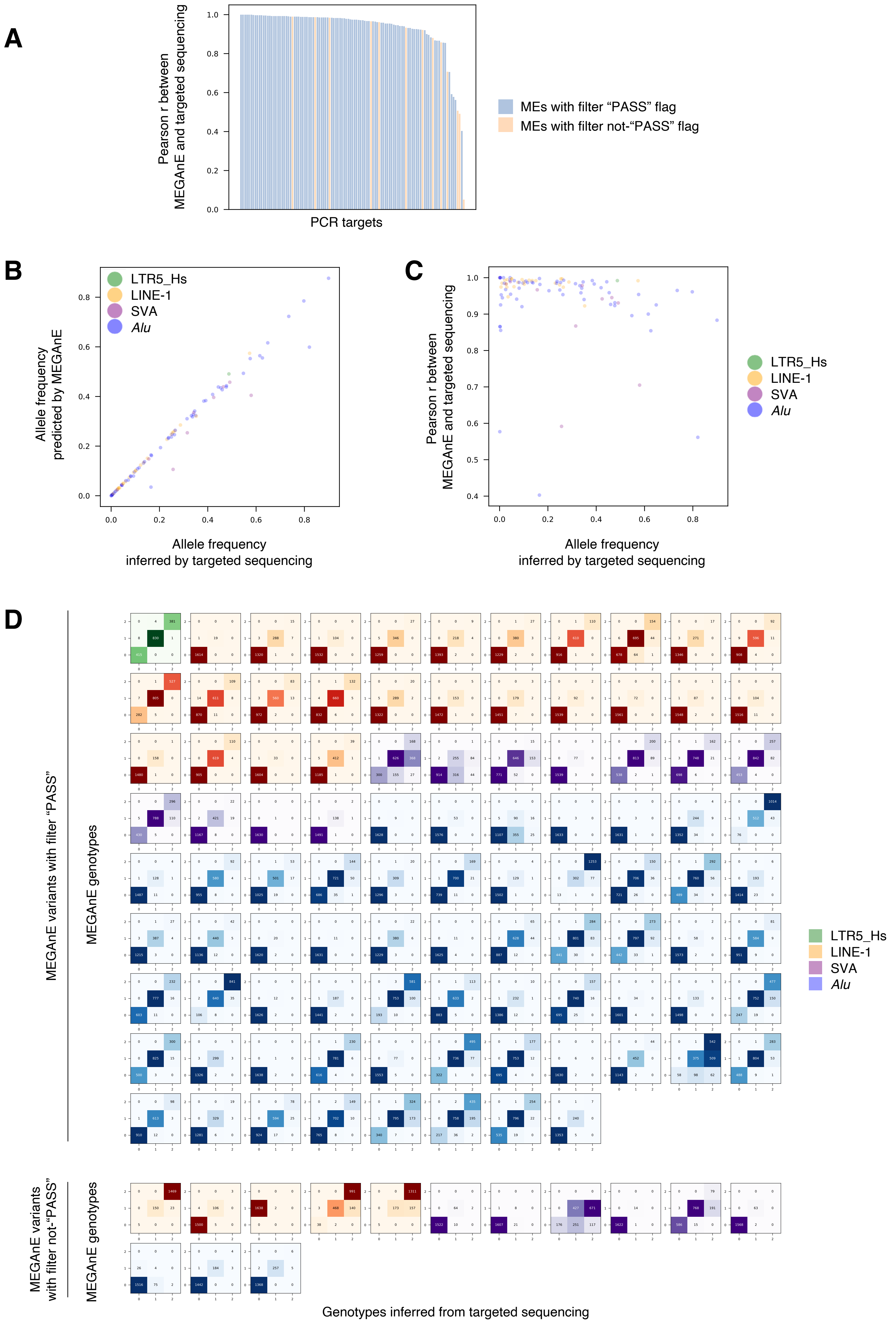
A
Pearson r between MEGAnE and targeted sequencing
PCR targets
MEs with filter "PASS" flag
MEs with filter not-"PASS" flag
B
Allele frequency predicted by MEGAnE
Allele frequency inferred by targeted sequencing
LTR5_Hs
LINE-1
SVA
Alu
C
Pearson r between MEGAnE and targeted sequencing
Allele frequency inferred by targeted sequencing
LTR5_Hs
LINE-1
SVA
Alu
D
MEGAnE variants with filter "PASS"
MEGAnE genotypes
MEGAnE variants with filter not-"PASS"
MEGAnE genotypes
Genotypes inferred from targeted sequencing
LTR5_Hs
LINE-1
SVA
Alu

### Fig. S15

| WGS depth | Number of individuals | MEGAnE filter | Number of sites genotyped by targeted seq | Number of sites concordant with targeted seq | % concordance |
|---:|---:|---:|---:|---:|---:|
| 25x | 888 | PASS | 80235 | 77725 | 97% |
| 25x | 888 | Not-PASS | 15840 | 13782 | 87% |
| 15x | 759 | PASS | 68684 | 66403 | 97% |
| 15x | 759 | Not-PASS | 12818 | 10876 | 85% |
| 25x or 15x | 1647 | PASS | 157084 | 152020 | 97% |
| 25x or 15x | 1647 | Not-PASS | 22877 | 19506 | 85% |

### Fig. S16

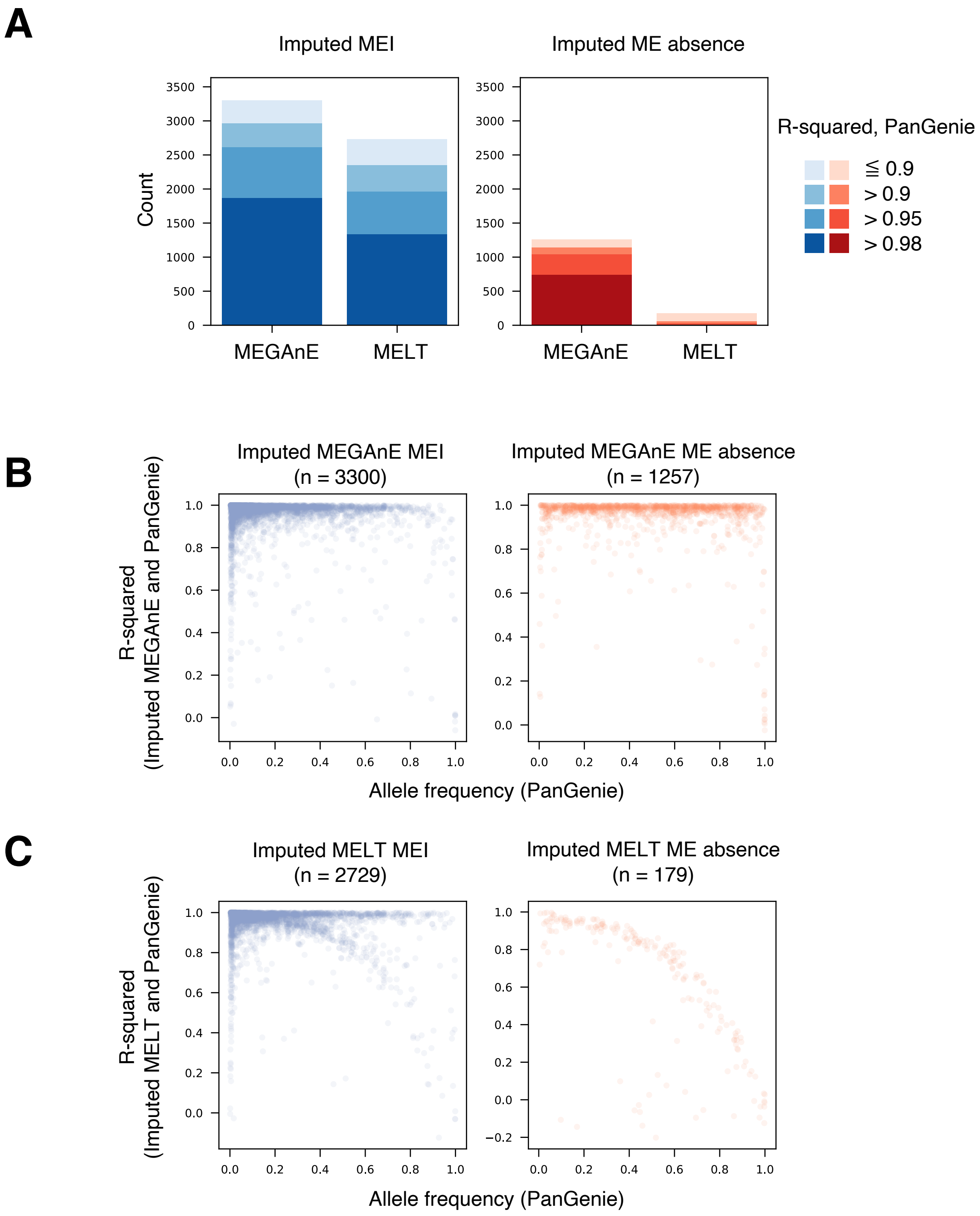
A
Imputed MEI
Imputed ME absence
Count
MEGAnE
MELT
R-squared, PanGenie
≦ 0.9
> 0.9
> 0.95
> 0.98
B
Imputed MEGAnE MEI
(n = 3300)
Imputed MEGAnE ME absence
(n = 1257)
R-squared
(Imputed MEGAnE and PanGenie)
Allele frequency (PanGenie)
C
Imputed MELT MEI
(n = 2729)
Imputed MELT ME absence
(n = 179)
R-squared
(Imputed MELT and PanGenie)
Allele frequency (PanGenie)

### Fig. S17

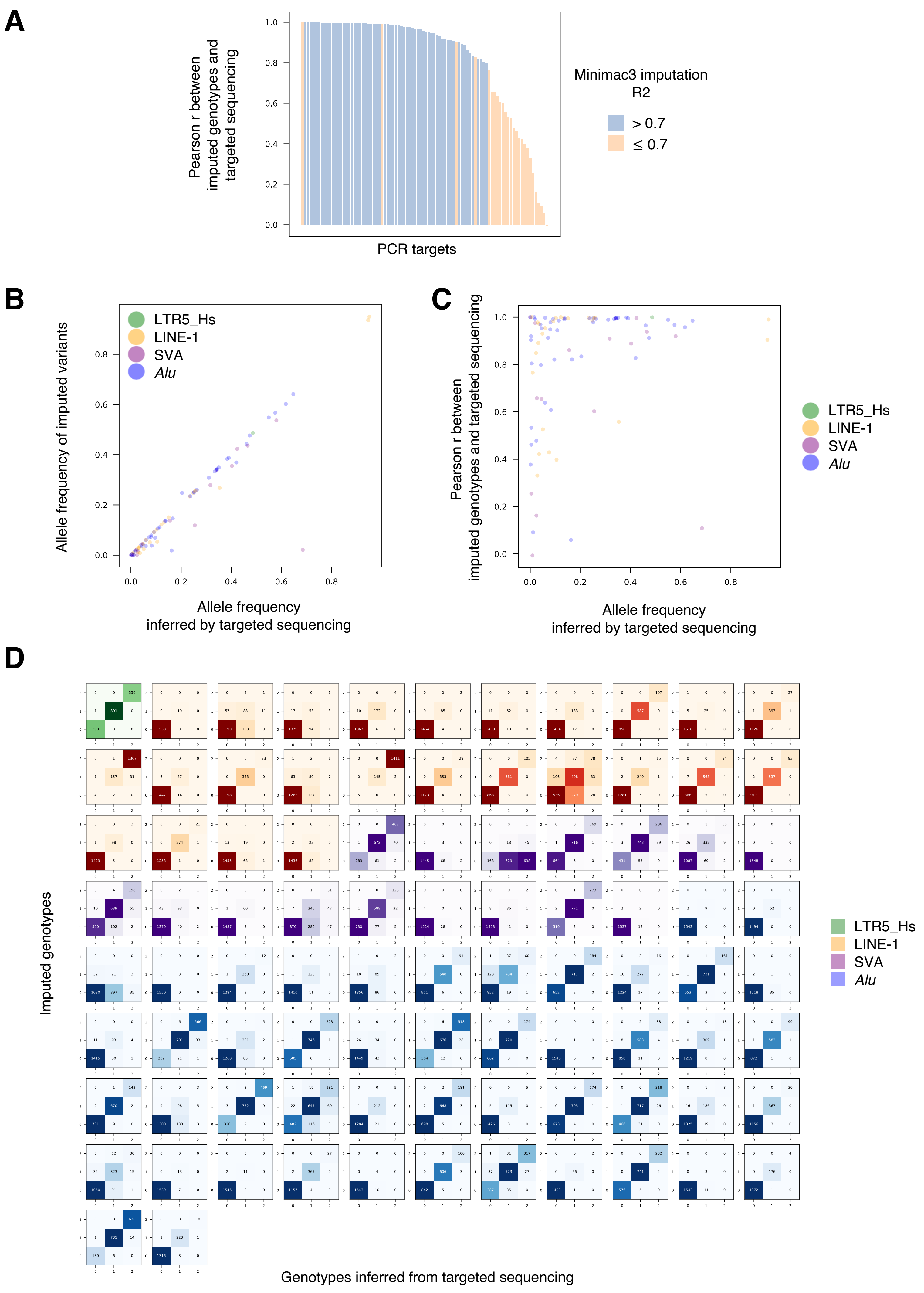
A
Pearson r between imputed genotypes and targeted sequencing
PCR targets
Minimac3 imputation R2
> 0.7
≤ 0.7
B
Allele frequency of imputed variants
Allele frequency inferred by targeted sequencing
LTR5_Hs
LINE-1
SVA
Alu
C
Pearson r between imputed genotypes and targeted sequencing
Allele frequency inferred by targeted sequencing
LTR5_Hs
LINE-1
SVA
Alu
D
Imputed genotypes
Genotypes inferred from targeted sequencing
LTR5_Hs
LINE-1
SVA
Alu

### Fig. S18

**A**

| MEGAnE Step | Thread number | Wall time | Total CPU time | Max memory |
|---|---|---|---|---|
| Step 1 | 1 | 1 h 56 min | 1 h 55 min | 5.0 GB |
| Step 1 | 2 | 1 h 13 min | 1 h 55 min | 5.1 GB |
| Step 1 | 4 | 53 min 16 sec | 2 h 5 min | 8.3 GB |
| Step 1 | 8 | 40 min 49 sec | 2 h 6 min | 17.0 GB |
| Step 1 | 12 | 35 min 48 sec | 2 h 10 min | 27.9 GB |

**B**

| MEGAnE Step | Thread number | Wall time | Summed wall time of 2,504 samples | Max memory |
|---|---|---|---|---|
| Step 0 | 1 | 13 min 6 sec | NA | 49.3 GB |
| Step 1 | 200 threads (2 threads x 100 jobs) | < 2 days | 179 days | 11.4GB / sample |
| Step 2 | 1 | 2 h 33 min | NA | 8.9 GB |

**C**

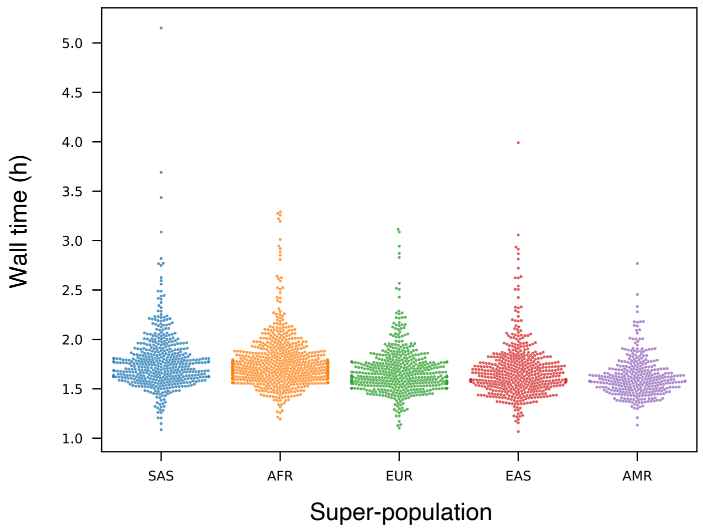

**D**

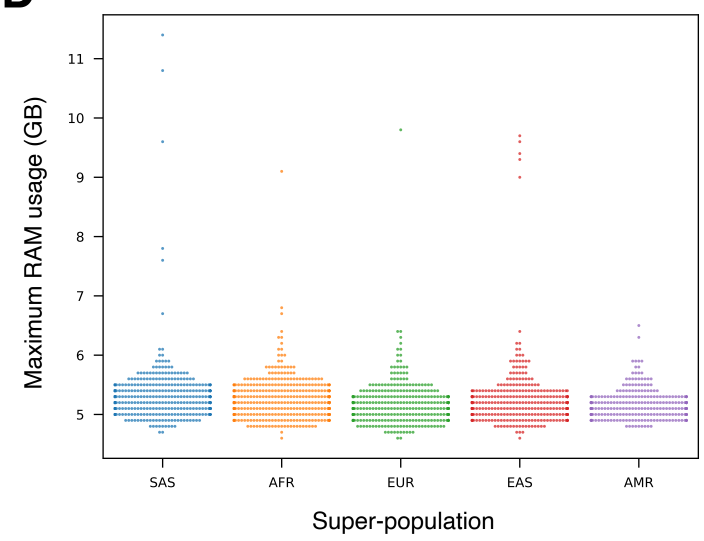

### Fig. S19

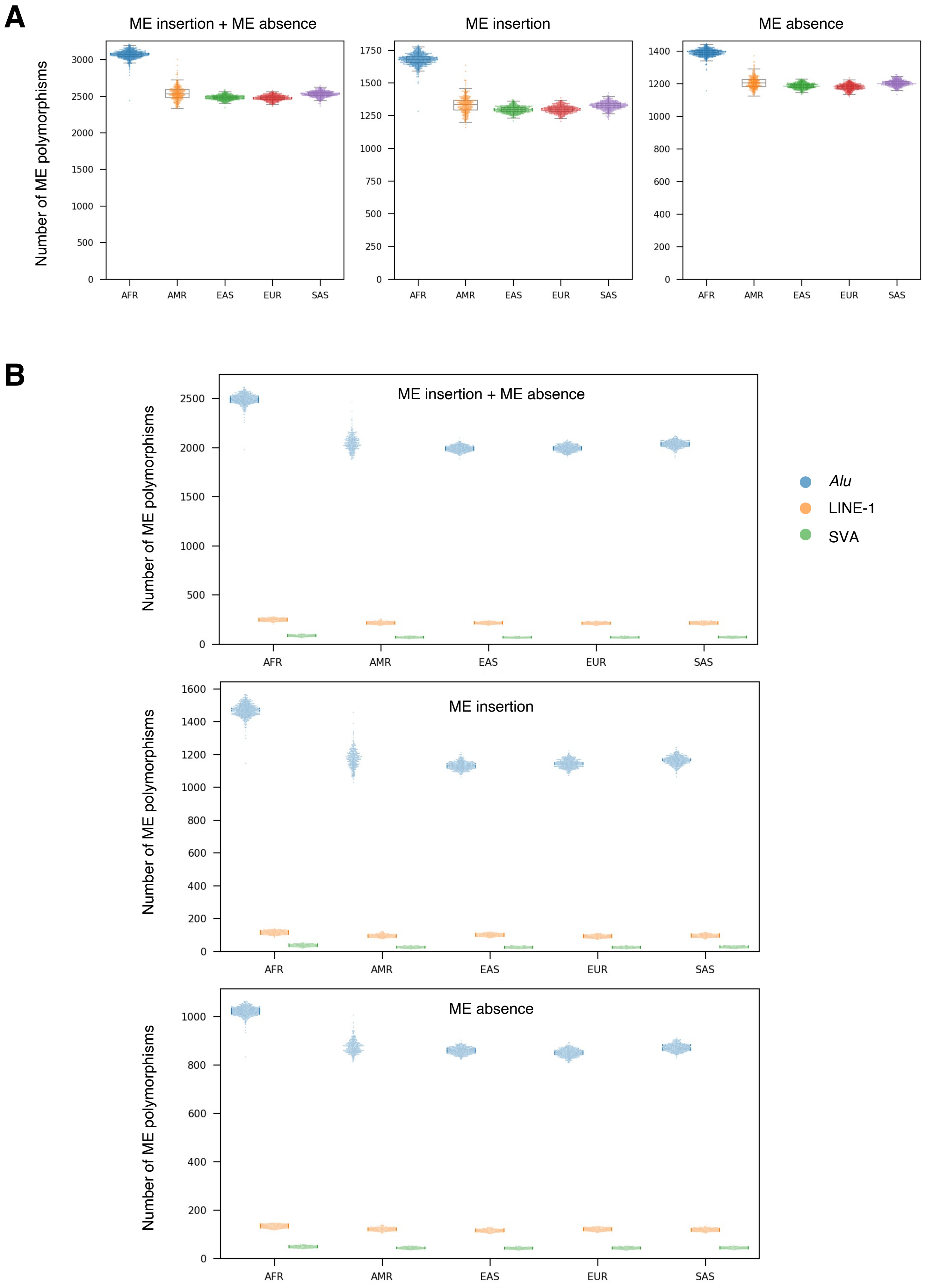
A
ME insertion + ME absence
ME insertion
ME absence
Number of ME polymorphisms
AFR
AMR
EAS
EUR
SAS
B
ME insertion + ME absence
ME insertion
ME absence
Number of ME polymorphisms
Alu
LINE-1
SVA

### Fig. S20

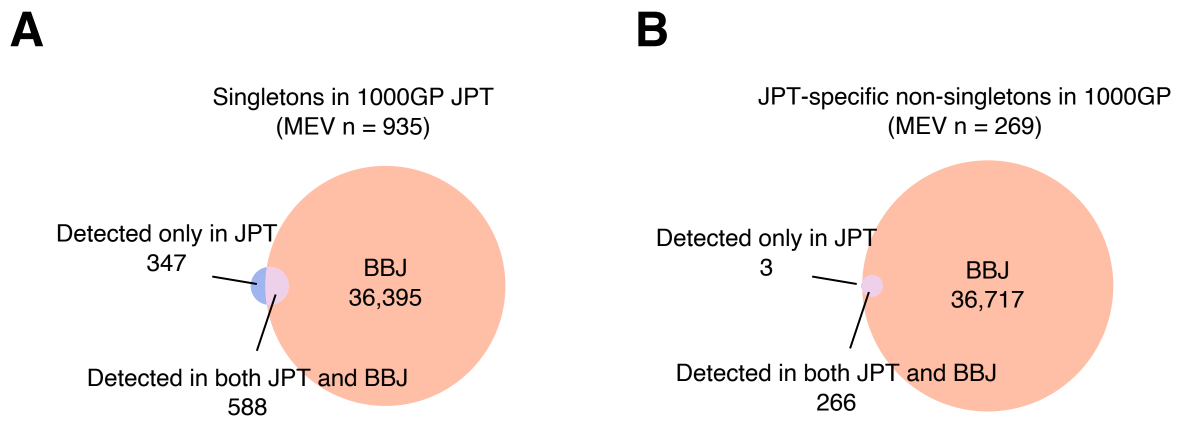
A
Singletons in 1000GP JPT
(MEV n = 935)
Detected only in JPT
347
BBJ
36,395
Detected in both JPT and BBJ
588
B
JPT-specific non-singletons in 1000GP
(MEV n = 269)
Detected only in JPT
3
BBJ
36,717
Detected in both JPT and BBJ
266

### Fig. S21

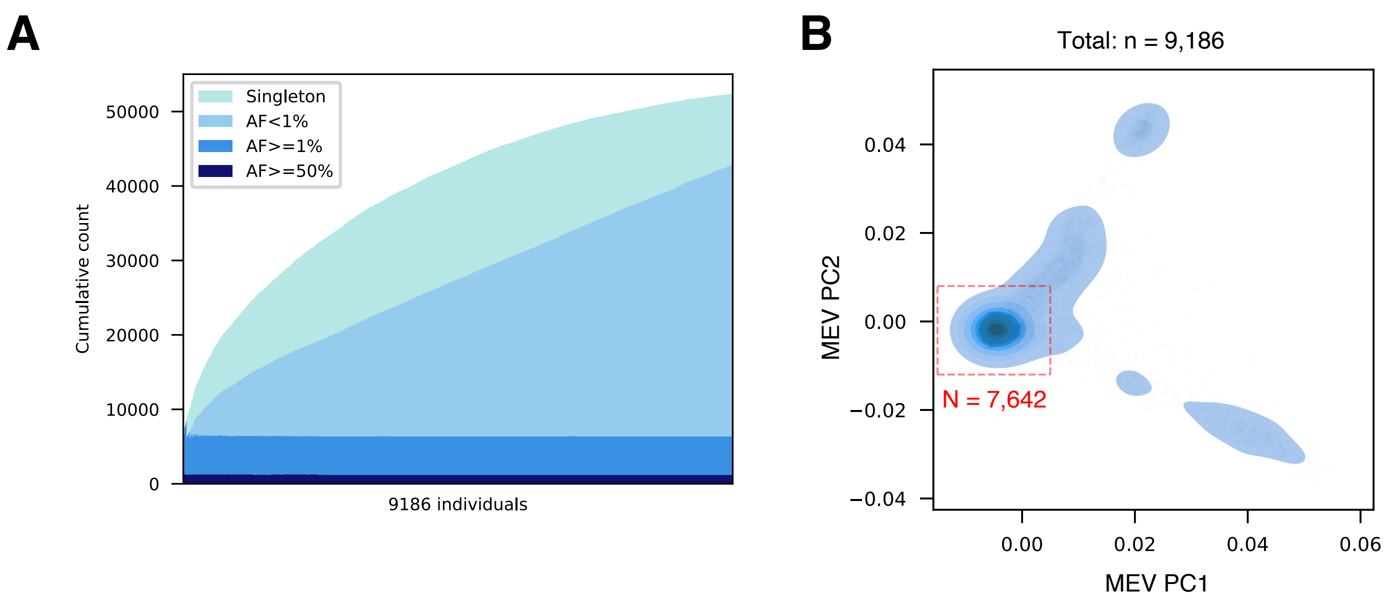
A
Singleton
AF<1%
AF>=1%
AF>=50%
Cumulative count
50000
40000
30000
20000
10000
0
9186 individuals
B
Total: n = 9,186
MEV PC2
0.04
0.02
0.00
−0.02
−0.04
N = 7,642
0.00
0.02
0.04
0.06
MEV PC1

### Fig. S22

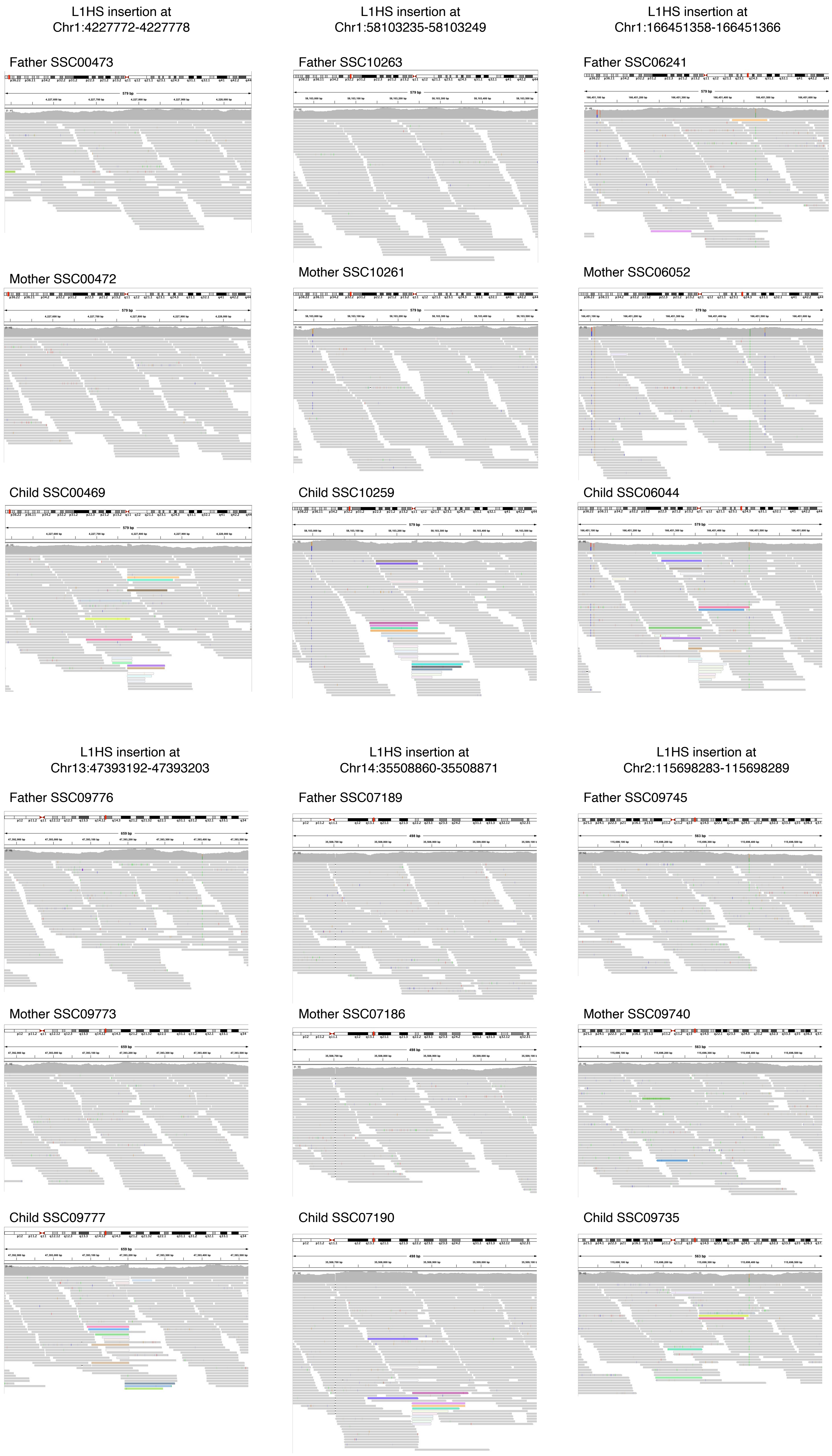
L1HS insertion at
Chr1:4227772-4227778
L1HS insertion at
Chr1:58103235-58103249
L1HS insertion at
Chr1:166451358-166451366
Father SSC00473
Father SSC10263
Father SSC06241
Mother SSC00472
Mother SSC10261
Mother SSC06052
Child SSC00469
Child SSC10259
Child SSC06044
L1HS insertion at
Chr13:47393192-47393203
L1HS insertion at
Chr14:35508860-35508871
L1HS insertion at
Chr2:115698283-115698289
Father SSC09776
Father SSC07189
Father SSC09745
Mother SSC09773
Mother SSC07186
Mother SSC09740
Child SSC09777
Child SSC07190
Child SSC09735

### Fig. S22, continued

L1HS insertion at
Chr2:194830322-194830336

Father SSC12045

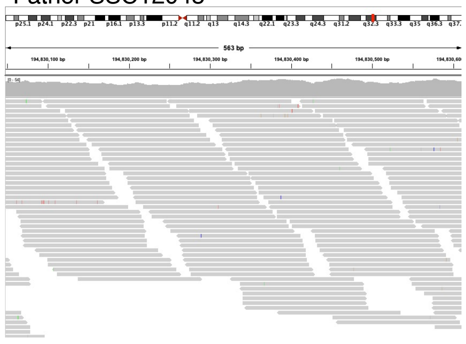

Mother SSC12039

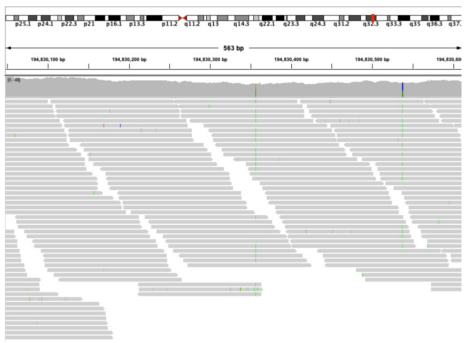

Child SSC12044

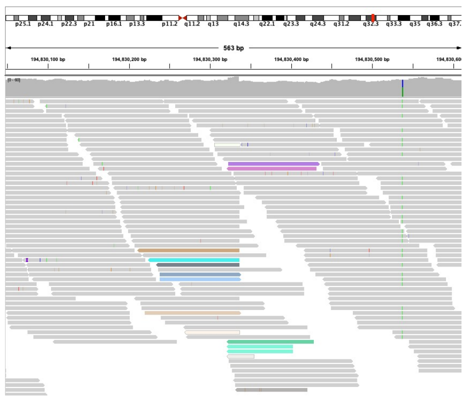

L1HS insertion at
Chr21:25428297-25428314

Father SSC08661

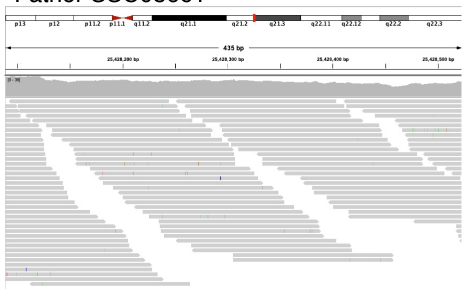

Mother SSC08660

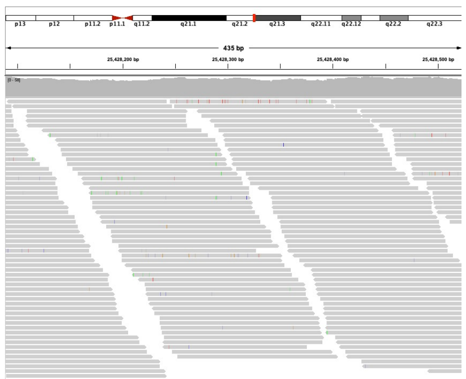

Child SSC08659

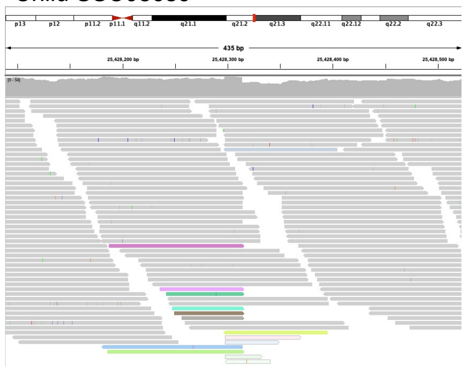

L1HS insertion at
Chr3:9093236-9093247

Father SSC01102

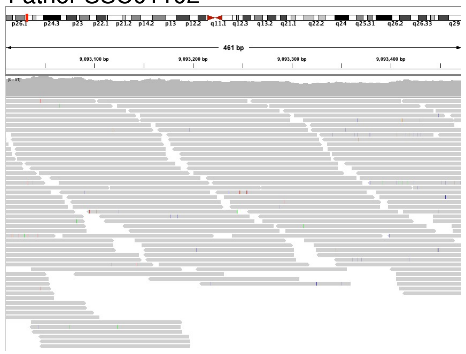

Mother SSC01117

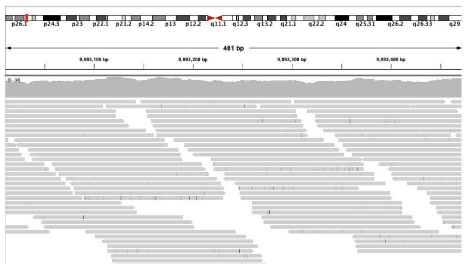

Child SSC01113

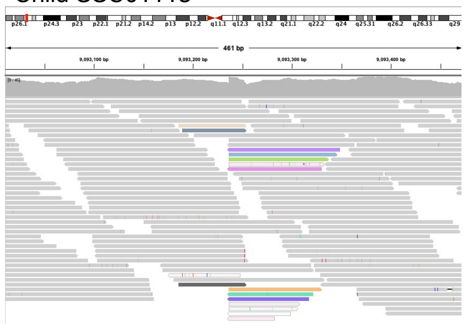

L1HS insertion at
Chr6:8063530-8063545

Father SSC12787

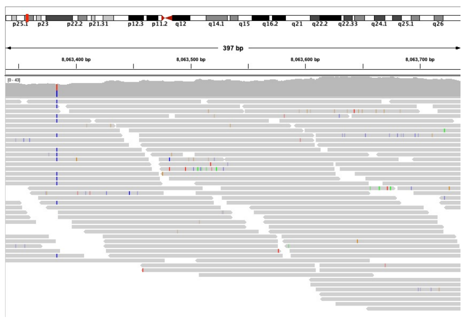

Mother SSC12783

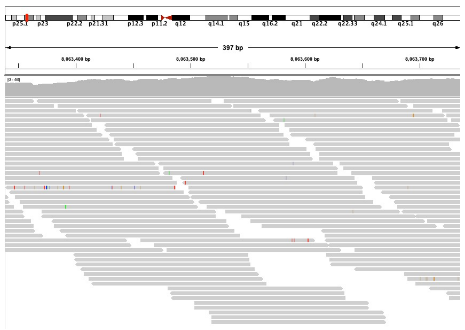

Child SSC12870

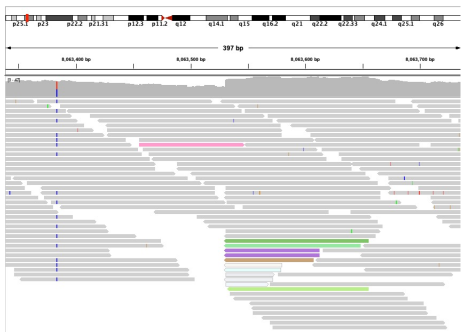

L1HS insertion at
Chr8:6366924-6366940

Father SSC00276

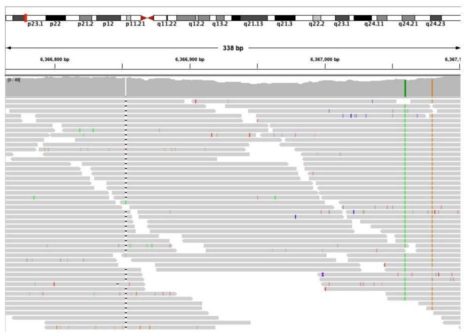

Mother SSC12064

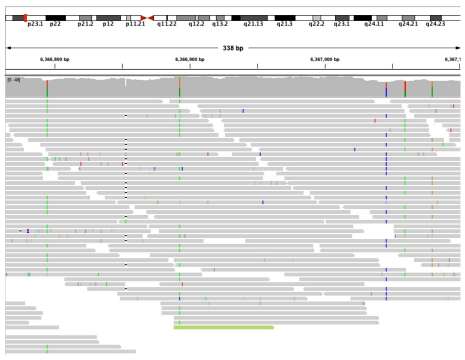

Child SSC00269

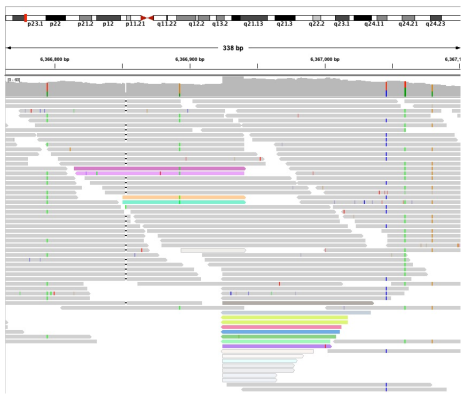

L1HS insertion at
Chr8:17458428-17458446

Father SSC06760

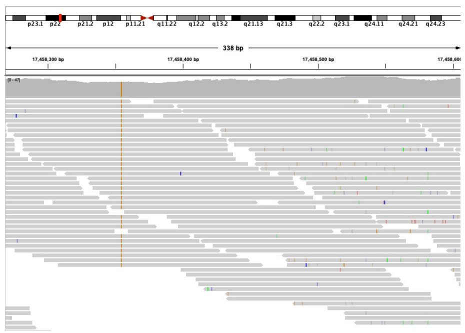

Mother SSC06757

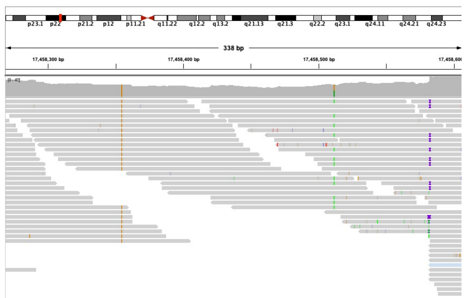

Child SSC06754

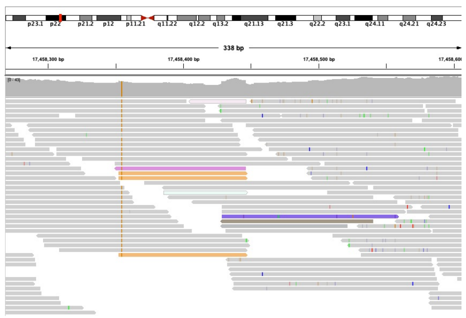

### Fig. S22, continued

L1HS insertion at
Chr9:74570049-74570058

Father SSC11336

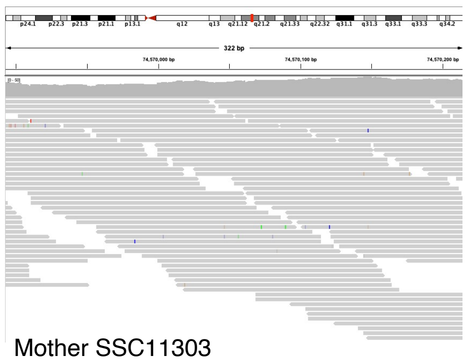

Mother SSC11303

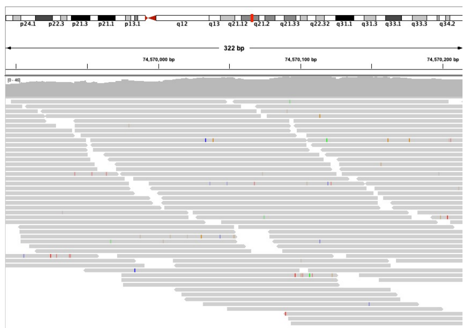

Child SSC11300

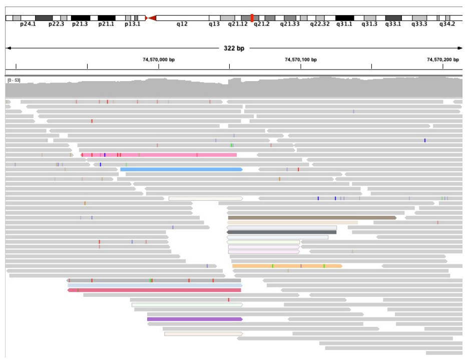

### Fig. S23

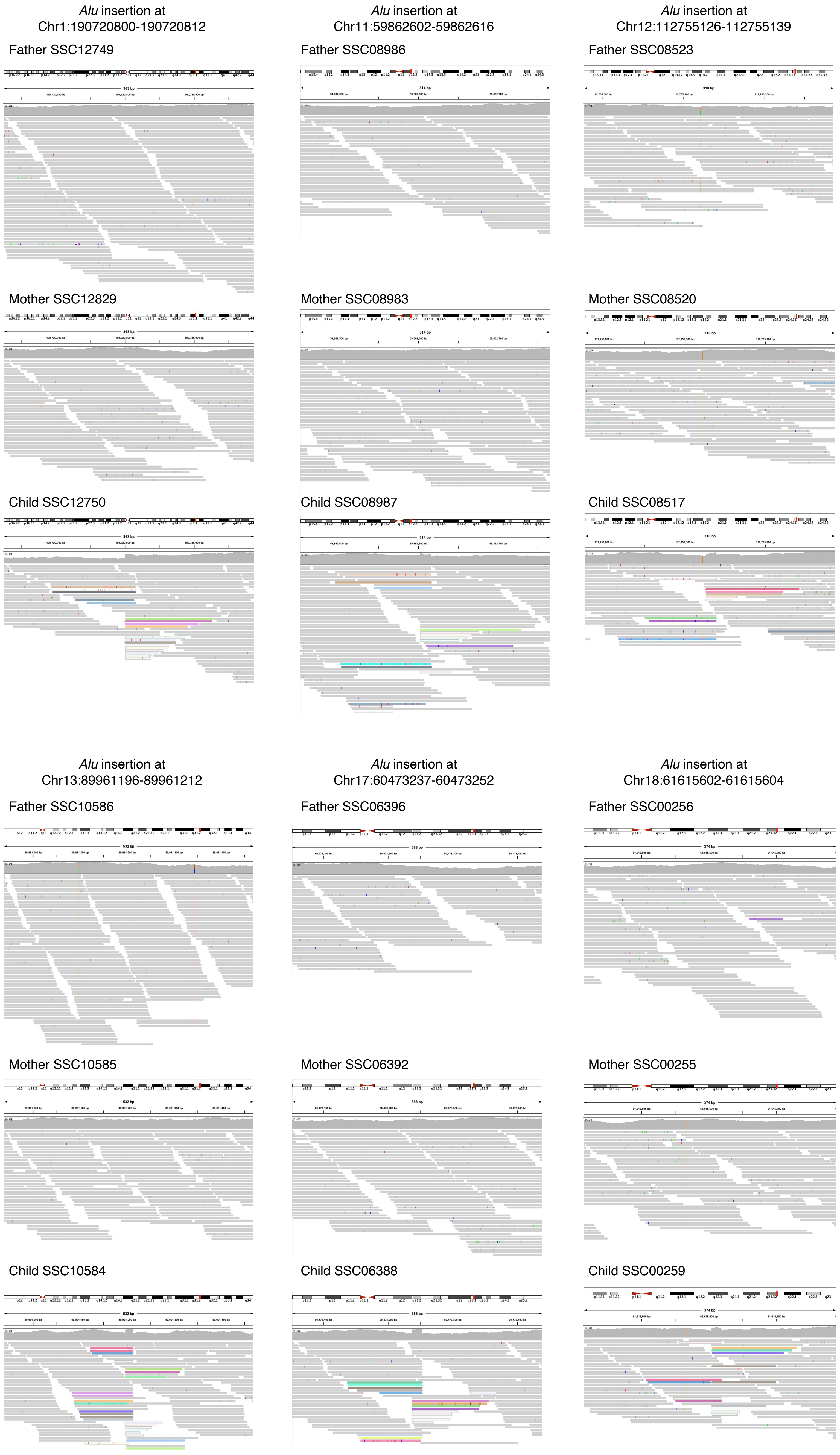
*Alu* insertion at Chr1:190720800-190720812
*Alu* insertion at Chr11:59862602-59862616
*Alu* insertion at Chr12:112755126-112755139
Father SSC12749
Father SSC08986
Father SSC08523
Mother SSC12829
Mother SSC08983
Mother SSC08520
Child SSC12750
Child SSC08987
Child SSC08517
*Alu* insertion at Chr13:89961196-89961212
*Alu* insertion at Chr17:60473237-60473252
*Alu* insertion at Chr18:61615602-61615604
Father SSC10586
Father SSC06396
Father SSC00256
Mother SSC10585
Mother SSC06392
Mother SSC00255
Child SSC10584
Child SSC06388
Child SSC00259

### Fig. S23, continued

*Alu* insertion at Chr18:76688653-76688666

Father SSC00892

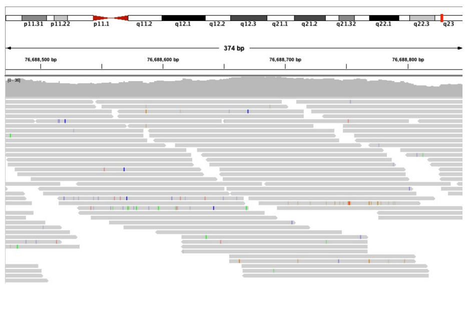

Mother SSC00893

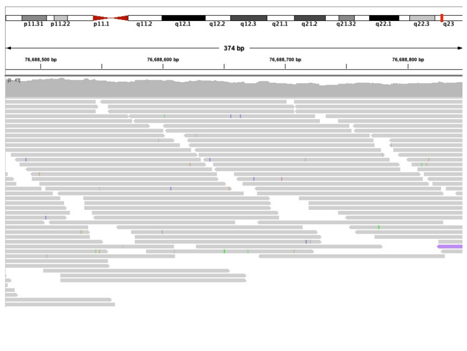

Child SSC00894

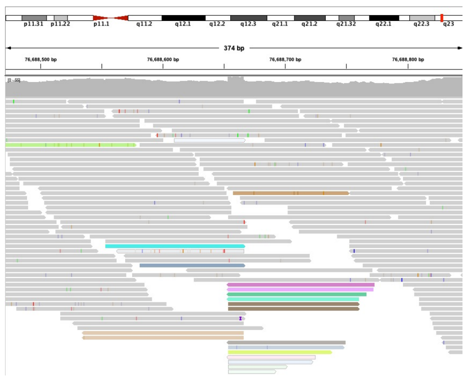

*Alu* insertion at Chr19:30950094-30950109

Father SSC05039

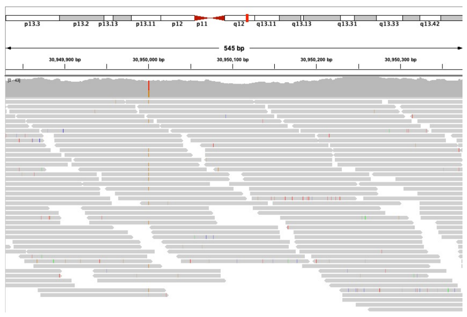

Mother SSC05032

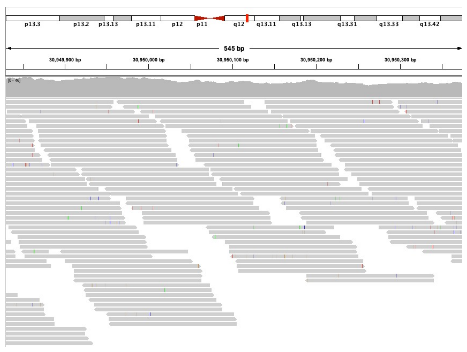

Child SSC05040

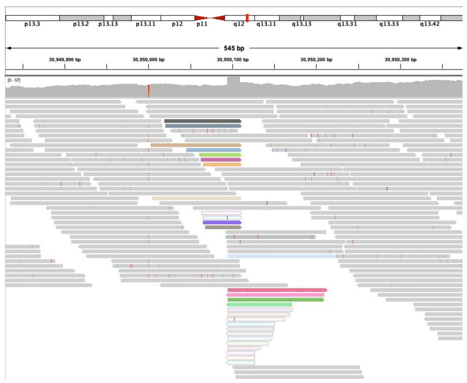

*Alu* insertion at Chr3:137876063-137876078

Father SSC04389

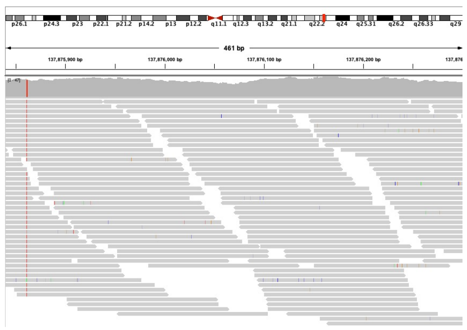

Mother SSC04385

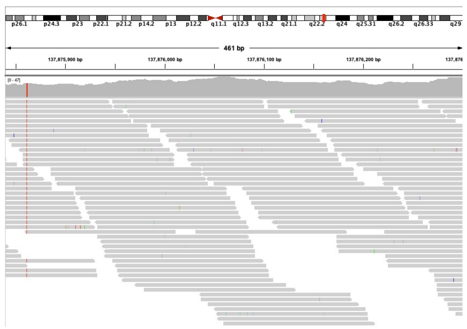

Child SSC04382

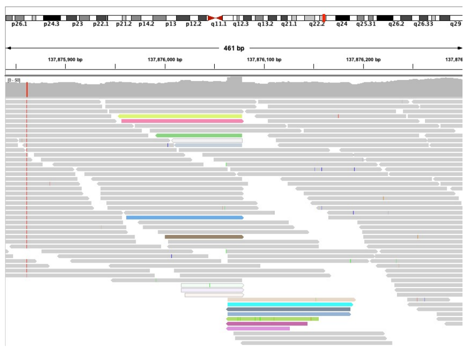

*Alu* insertion at Chr3:188327270-188327280

Father SSC06896

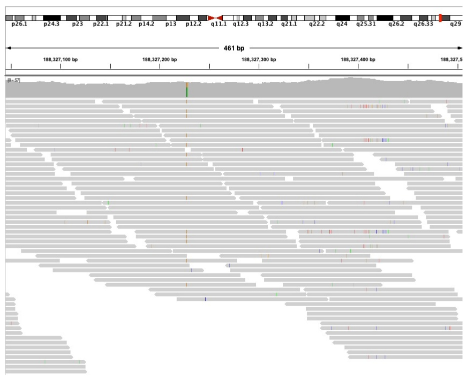

Mother SSC06890

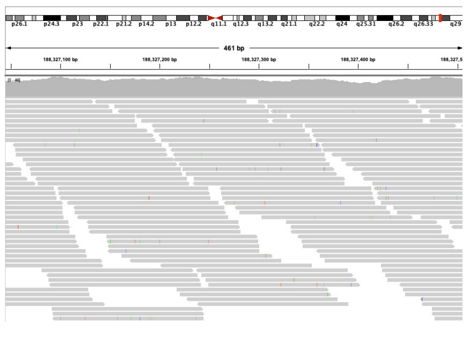

Child SSC06897

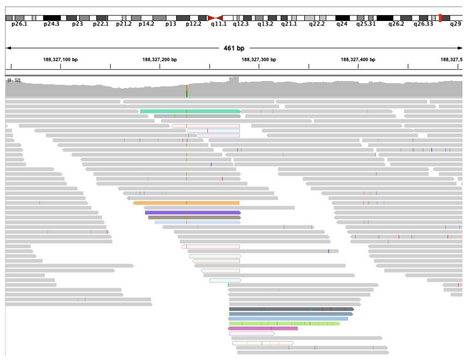

*Alu* insertion at Chr4:155581528-155581538

Father SSC09007

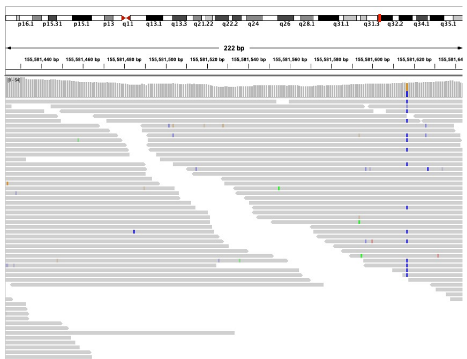

Mother SSC08999

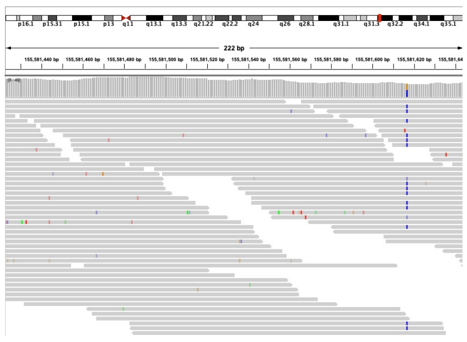

Child SSC09008

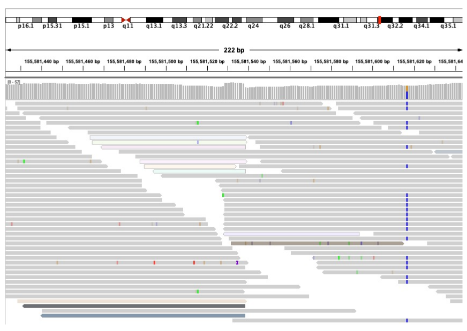

*Alu* insertion at Chr4:160407103-160407109

Father SSC02711

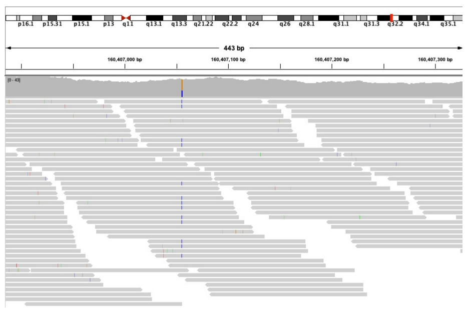

Mother SSC02693

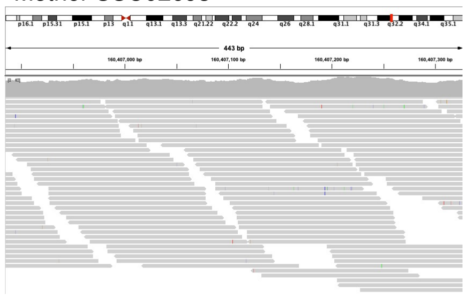

Child SSC02682

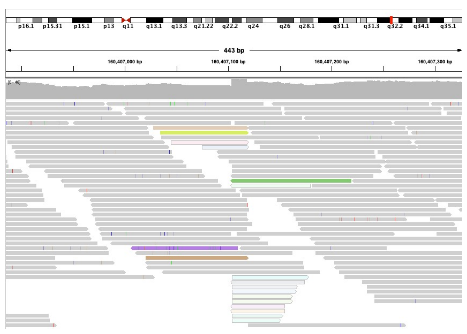

### Fig. S23, continued

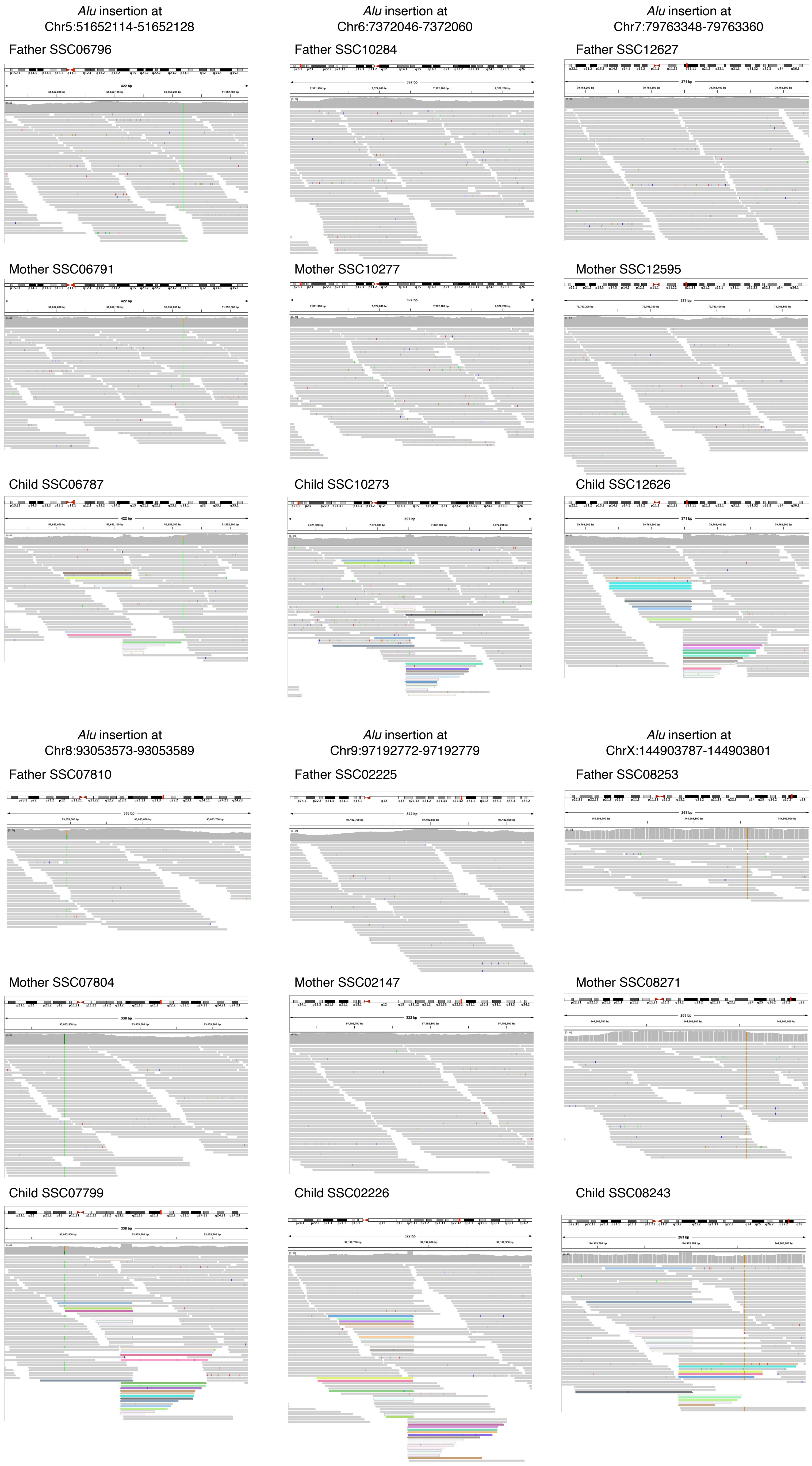

### Fig. S24

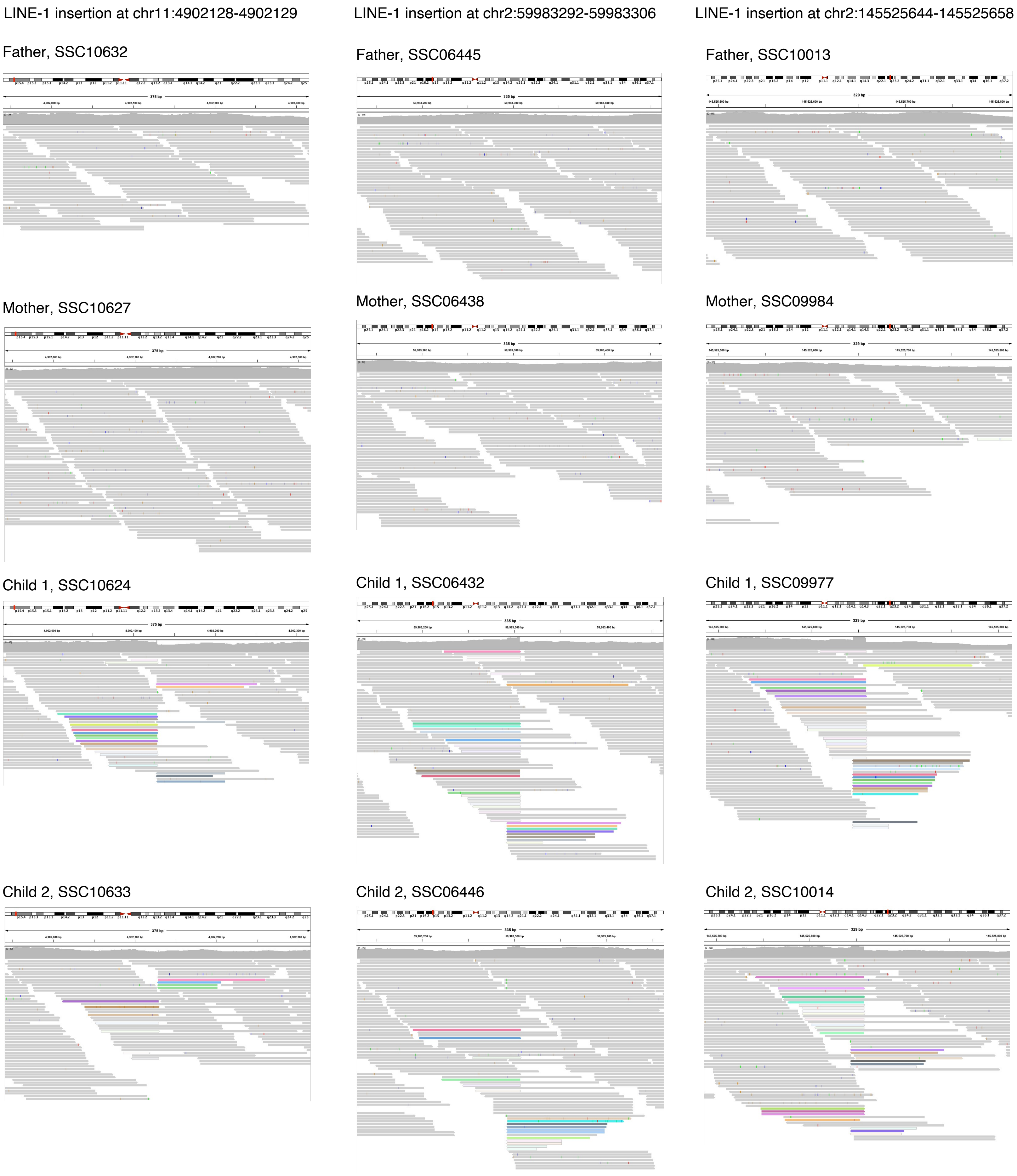
LINE-1 insertion at chr11:4902128-4902129
LINE-1 insertion at chr2:59983292-59983306
LINE-1 insertion at chr2:145525644-145525658
Father, SSC10632
Father, SSC06445
Father, SSC10013
Mother, SSC10627
Mother, SSC06438
Mother, SSC09984
Child 1, SSC10624
Child 1, SSC06432
Child 1, SSC09977
Child 2, SSC10633
Child 2, SSC06446
Child 2, SSC10014

### Fig. S25

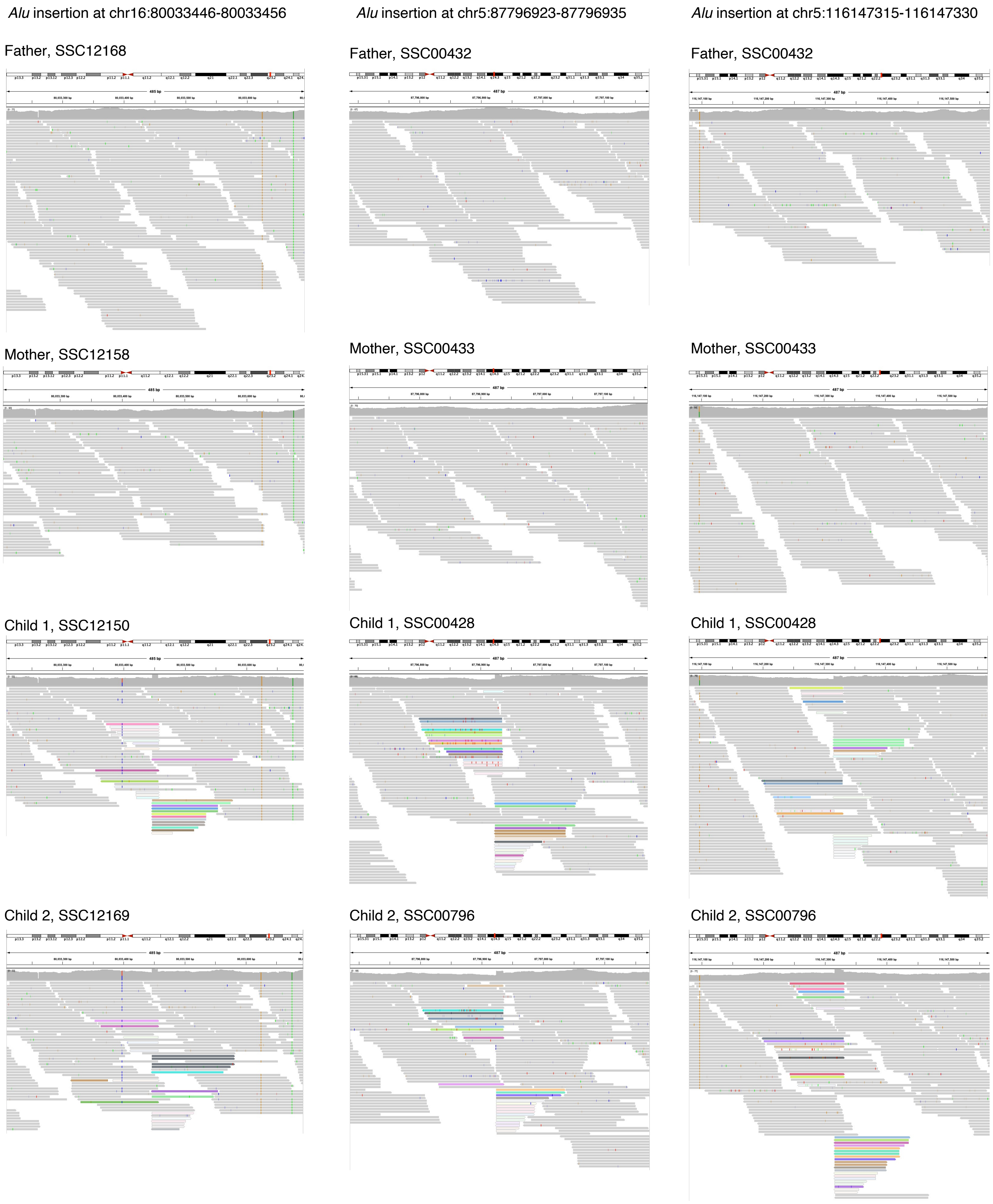

Alu insertion at chr16:80033446-80033456
Alu insertion at chr5:87796923-87796935
Alu insertion at chr5:116147315-116147330
Father, SSC12168
Father, SSC00432
Father, SSC00432
Mother, SSC12158
Mother, SSC00433
Mother, SSC00433
Child 1, SSC12150
Child 1, SSC00428
Child 1, SSC00428
Child 2, SSC12169
Child 2, SSC00796
Child 2, SSC00796

### Fig. S26

A

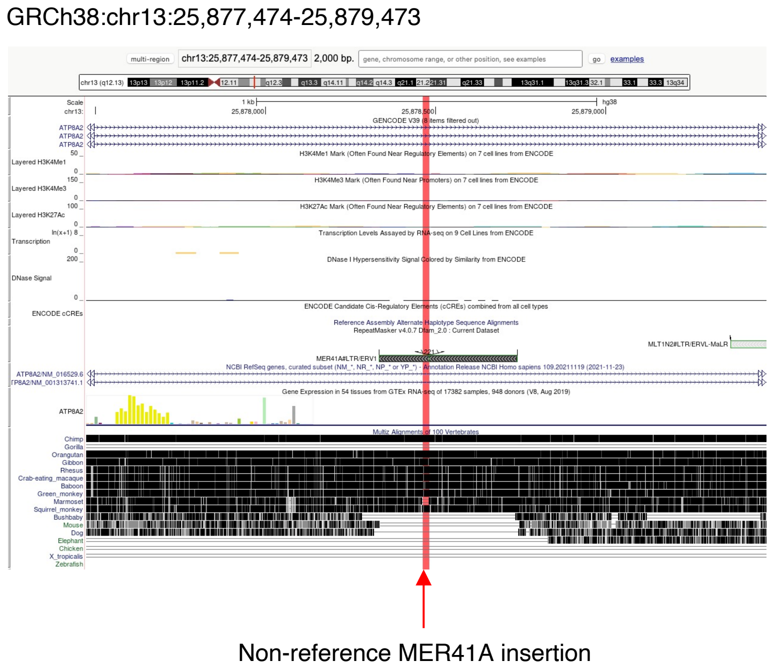

GRCh38:chr13:25,877,474-25,879,473
Non-reference MER41A insertion

B

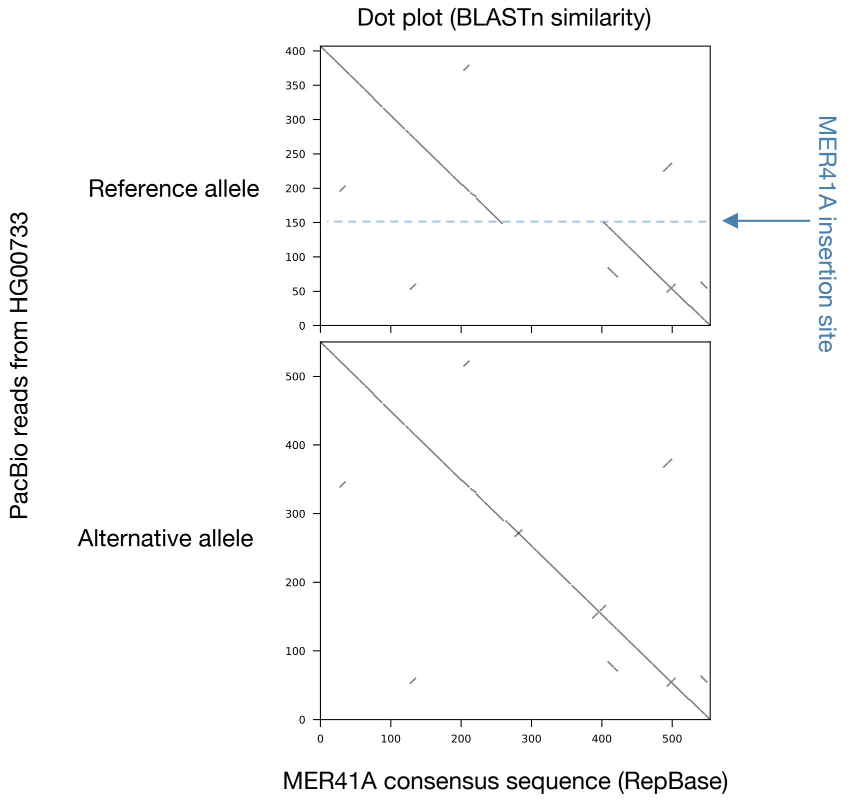

Dot plot (BLASTn similarity)
PacBio reads from HG00733
Reference allele
Alternative allele
MER41A insertion site
MER41A consensus sequence (RepBase)

C

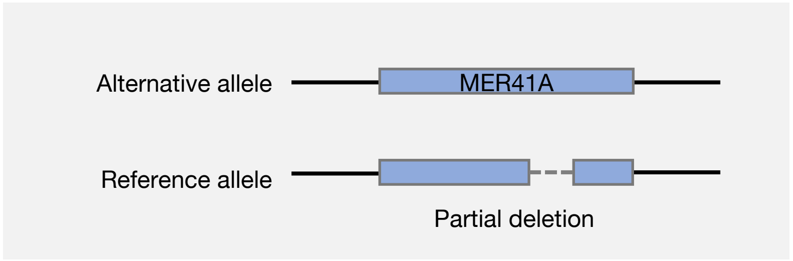

Alternative allele
MER41A
Reference allele
Partial deletion

### Fig. S27

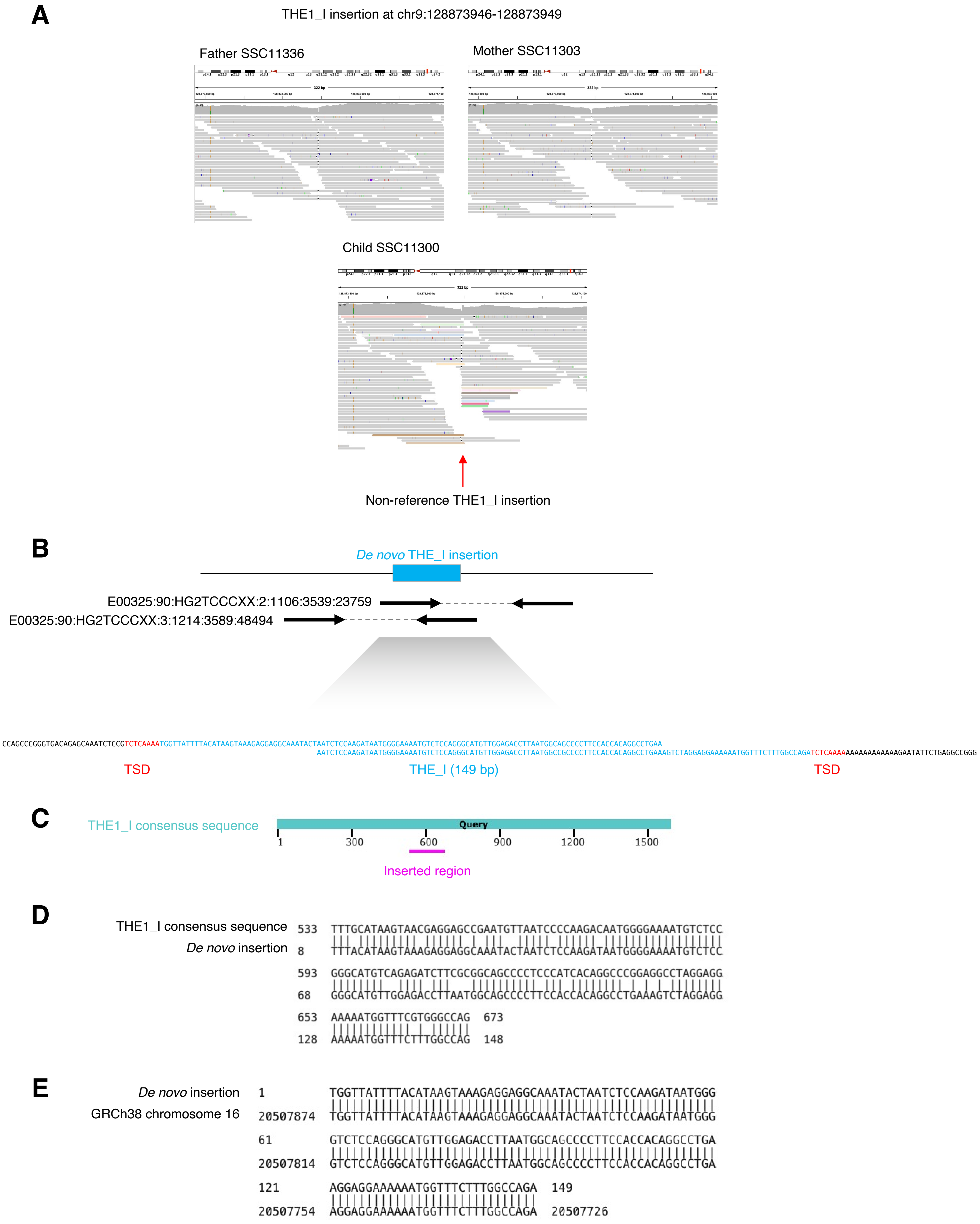
A
THE1_I insertion at chr9:128873946-128873949
Father SSC11336
Mother SSC11303
Child SSC11300
Non-reference THE1_I insertion
B
De novo THE_I insertion
E00325:90:HG2TCCCXX:2:1106:3539:23759
E00325:90:HG2TCCCXX:3:1214:3589:48494
CCAGCCCGGGTGACAGAGCAAATCTCCGTCTCAAAATGGTTATTTTACATAAGTAAAGAGGAGGCAAATACTAATCTCCAAGATAATGGGGAAAATGTCTCCAGGGCATGTTGGAGACCTTAATGGCAGCCCCTTCCACCACAGGCCTGAA
AATCTCCAAGATAATGGGGAAAATGTCTCCAGGGCATGTTGGAGACCTTAATGGCCGCCCCTTCCACCACAGGCCTGAAAGTCTAGGAGGAAAAAATGGTTTCTTTGGCCAGATCTCAAAAAAAAAAAAAAAAGAATATTCTGAGGCCGGG
TSD
THE_I (149 bp)
TSD
C
THE1_I consensus sequence
Query
1
300
600
900
1200
1500
Inserted region
D
THE1_I consensus sequence 533 TTTGCATAAGTAACGAGGAGCCGAATGTTAATCCCCAAGACAATGGGGAAAATGTCTCC
De novo insertion 8 TTTACATAAGTAAAGAGGAGGCAAATACTAATCTCCAAGATAATGGGGAAAATGTCTCC
593 GGGCATGTCAGAGATCTTCGCGGCAGCCCCTCCCATCACAGGCCCGGAGGCCTAGGAGG
68 GGGCATGTTGGAGACCTTAATGGCAGCCCCTTCCACCACAGGCCTGAAAGTCTAGGAGG
653 AAAAATGGTTTCGTGGGCCAG 673
128 AAAAATGGTTTCTTTGGCCAG 148
E
De novo insertion 1 TGGTTATTTTACATAAGTAAAGAGGAGGCAAATACTAATCTCCAAGATAATGGG
GRCh38 chromosome 16 20507874 TGGTTATTTTACATAAGTAAAGAGGAGGCAAATACTAATCTCCAAGATAATGGG
61 GTCTCCAGGGCATGTTGGAGACCTTAATGGCAGCCCCTTCCACCACAGGCCTGA
20507814 GTCTCCAGGGCATGTTGGAGACCTTAATGGCAGCCCCTTCCACCACAGGCCTGA
121 AGGAGGAAAAAATGGTTTCTTTGGCCAGA 149
20507754 AGGAGGAAAAAATGGTTTCTTTGGCCAGA 20507726

### Fig. S28

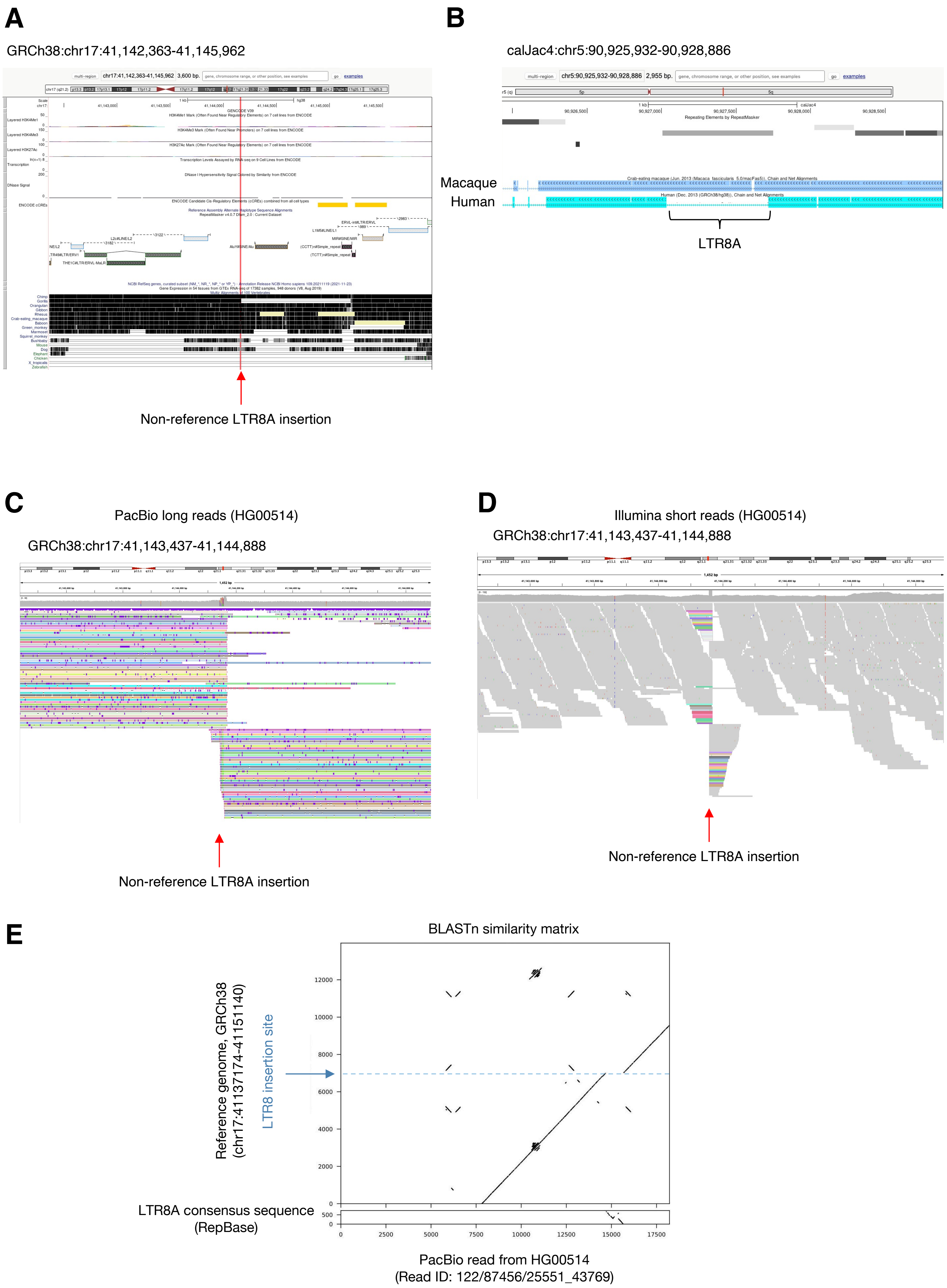
A
GRCh38:chr17:41,142,363-41,145,962
Non-reference LTR8A insertion
B
calJac4:chr5:90,925,932-90,928,886
Macaque
Human
LTR8A
C
PacBio long reads (HG00514)
GRCh38:chr17:41,143,437-41,144,888
Non-reference LTR8A insertion
D
Illumina short reads (HG00514)
GRCh38:chr17:41,143,437-41,144,888
Non-reference LTR8A insertion
E
BLASTn similarity matrix
Reference genome, GRCh38
(chr17:41137174-41151140)
LTR8 insertion site
LTR8A consensus sequence
(RepBase)
PacBio read from HG00514
(Read ID: 122/87456/25551_43769)

### Fig. S29

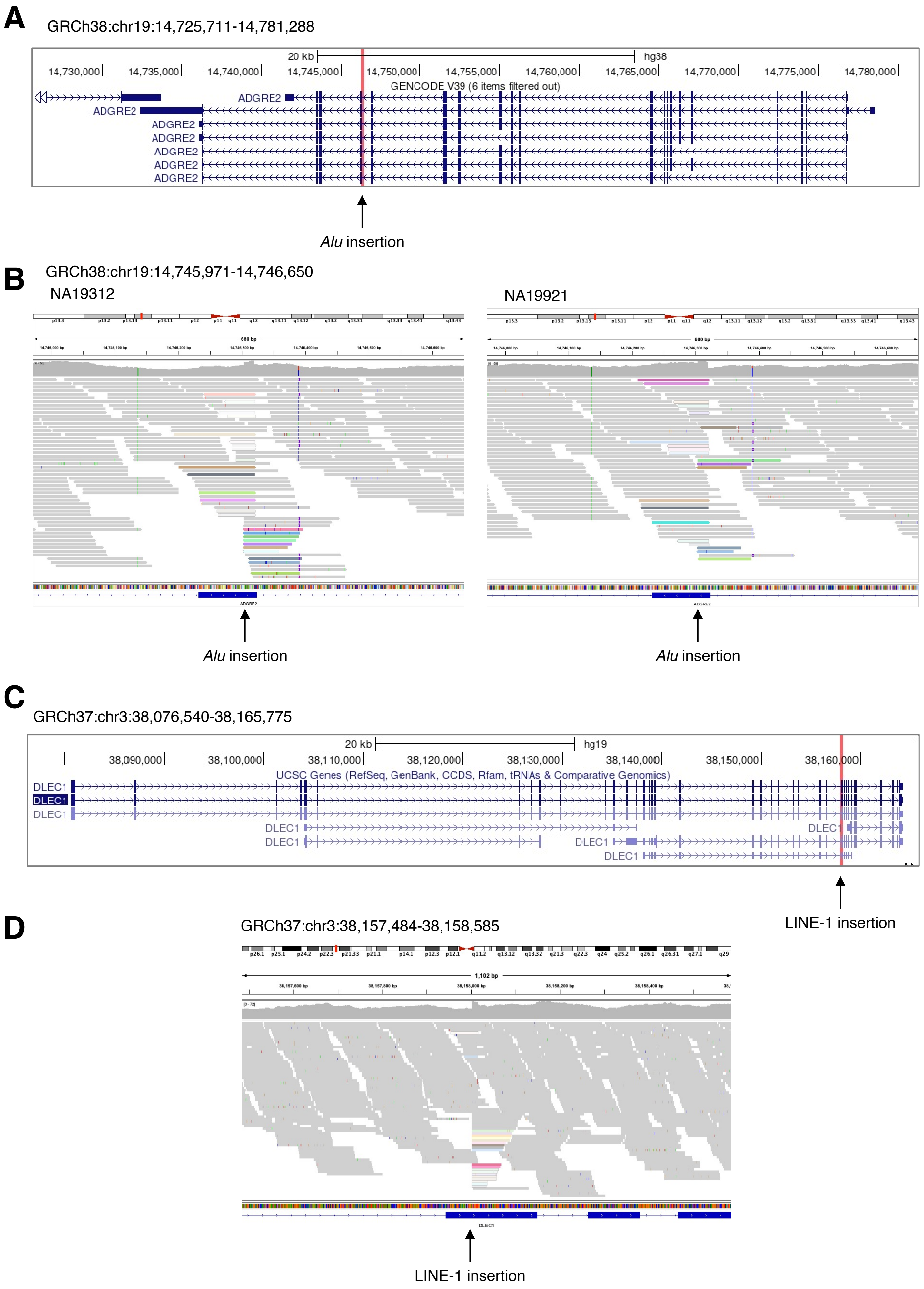
A
GRCh38:chr19:14,725,711-14,781,288
20 kb
hg38
14,730,000
14,735,000
14,740,000
14,745,000
14,750,000
14,755,000
14,760,000
14,765,000
14,770,000
14,775,000
14,780,000
GENCODE V39 (6 items filtered out)
ADGRE2
ADGRE2
ADGRE2
ADGRE2
ADGRE2
ADGRE2
ADGRE2
Alu insertion
B
GRCh38:chr19:14,745,971-14,746,650
NA19312
NA19921
Alu insertion
Alu insertion
C
GRCh37:chr3:38,076,540-38,165,775
20 kb
hg19
38,090,000
38,100,000
38,110,000
38,120,000
38,130,000
38,140,000
38,150,000
38,160,000
UCSC Genes (RefSeq, GenBank, CCDS, Rfam, tRNAs & Comparative Genomics)
DLEC1
DLEC1
DLEC1
DLEC1
DLEC1
DLEC1
DLEC1
LINE-1 insertion
D
GRCh37:chr3:38,157,484-38,158,585
LINE-1 insertion

### Fig. S30

**A**

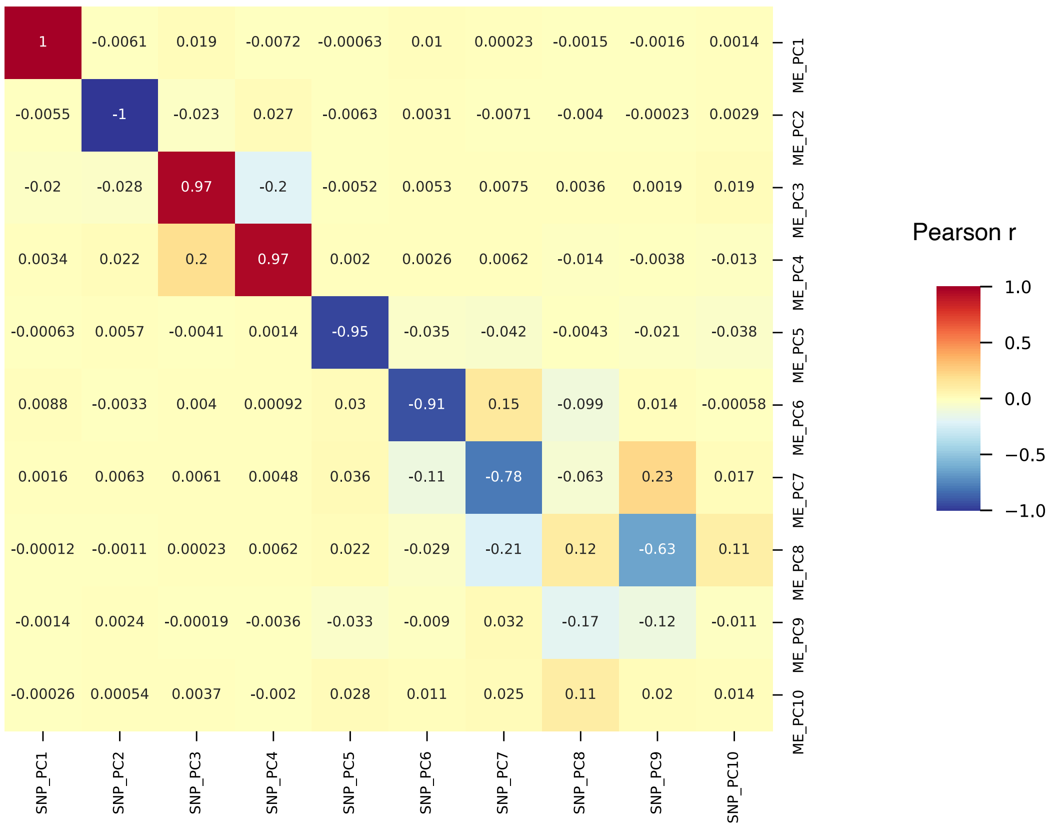
1 -0.0061 0.019 -0.0072 -0.00063 0.01 0.00023 -0.0015 -0.0016 0.0014
-0.0055 -1 -0.023 0.027 -0.0063 0.0031 -0.0071 -0.004 -0.00023 0.0029
-0.02 -0.028 0.97 -0.2 -0.0052 0.0053 0.0075 0.0036 0.0019 0.019
0.0034 0.022 0.2 0.97 0.002 0.0026 0.0062 -0.014 -0.0038 -0.013
-0.00063 0.0057 -0.0041 0.0014 -0.95 -0.035 -0.042 -0.0043 -0.021 -0.038
0.0088 -0.0033 0.004 0.00092 0.03 -0.91 0.15 -0.099 0.014 -0.00058
0.0016 0.0063 0.0061 0.0048 0.036 -0.11 -0.78 -0.063 0.23 0.017
-0.00012 -0.0011 0.00023 0.0062 0.022 -0.029 -0.21 0.12 -0.63 0.11
-0.0014 0.0024 -0.00019 -0.0036 -0.033 -0.009 0.032 -0.17 -0.12 -0.011
-0.00026 0.00054 0.0037 -0.002 0.028 0.011 0.025 0.11 0.02 0.014
ME_PC1
ME_PC2
ME_PC3
ME_PC4
ME_PC5
ME_PC6
ME_PC7
ME_PC8
ME_PC9
ME_PC10
SNP_PC1
SNP_PC2
SNP_PC3
SNP_PC4
SNP_PC5
SNP_PC6
SNP_PC7
SNP_PC8
SNP_PC9
SNP_PC10
Pearson r
1.0
0.5
0.0
−0.5
−1.0

**B**

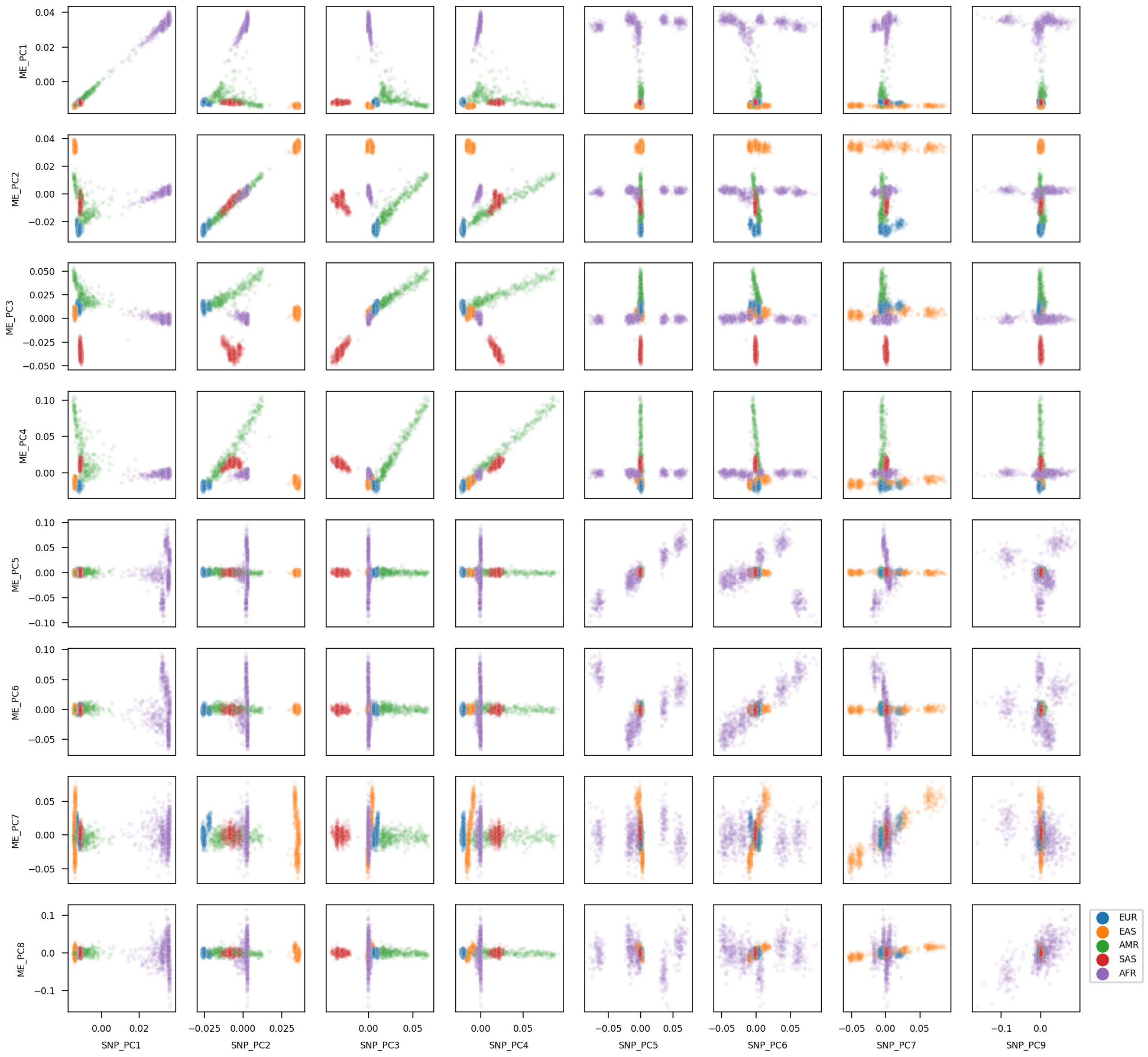
ME_PC1
ME_PC2
ME_PC3
ME_PC4
ME_PC5
ME_PC6
ME_PC7
ME_PC8
SNP_PC1
SNP_PC2
SNP_PC3
SNP_PC4
SNP_PC5
SNP_PC6
SNP_PC7
SNP_PC9
EUR
EAS
AMR
SAS
AFR

Fig. S31

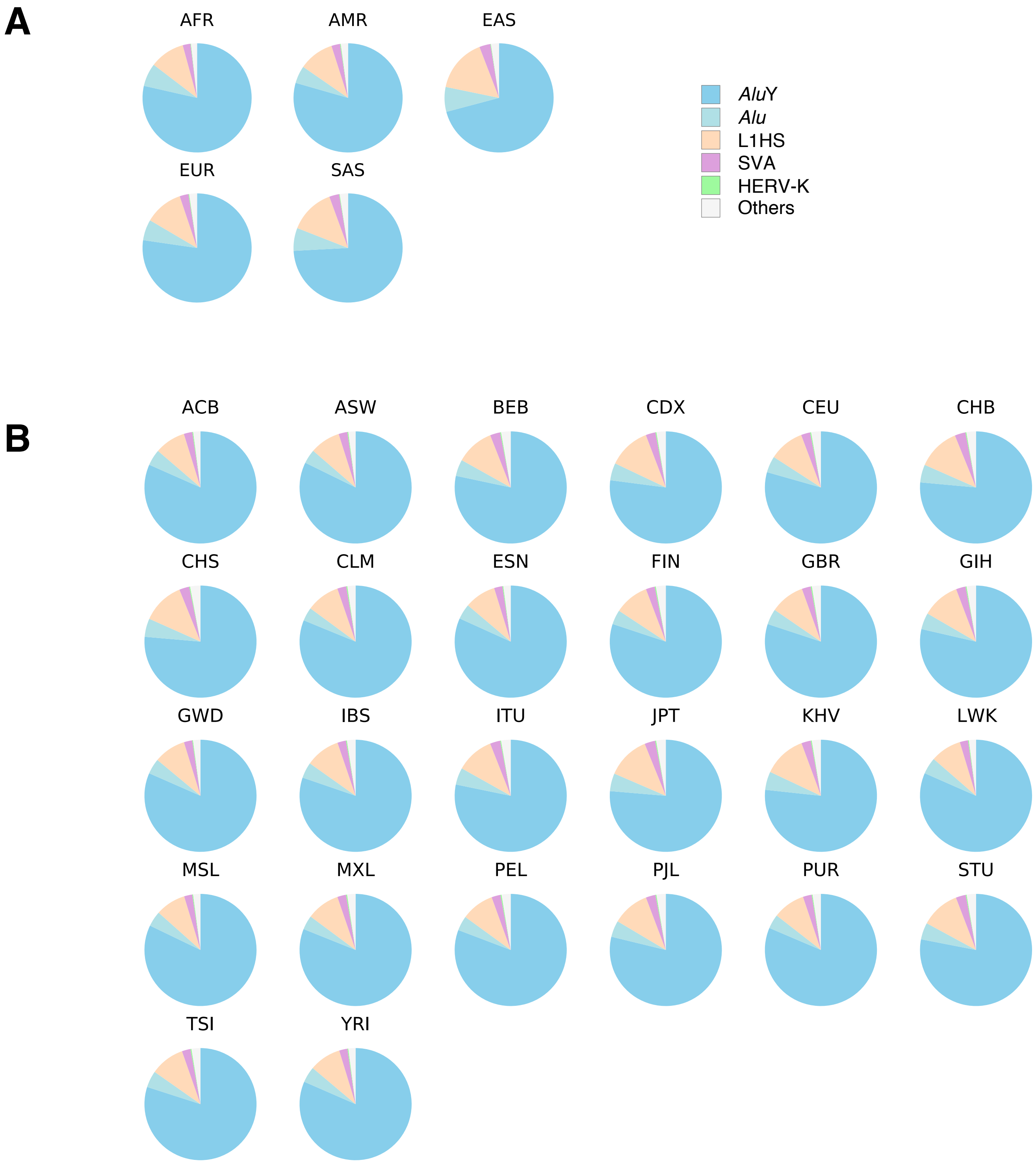
A
AFR
AMR
EAS
EUR
SAS
AluY
Alu
L1HS
SVA
HERV-K
Others
B
ACB
ASW
BEB
CDX
CEU
CHB
CHS
CLM
ESN
FIN
GBR
GIH
GWD
IBS
ITU
JPT
KHV
LWK
MSL
MXL
PEL
PJL
PUR
STU
TSI
YRI

### Fig. S32

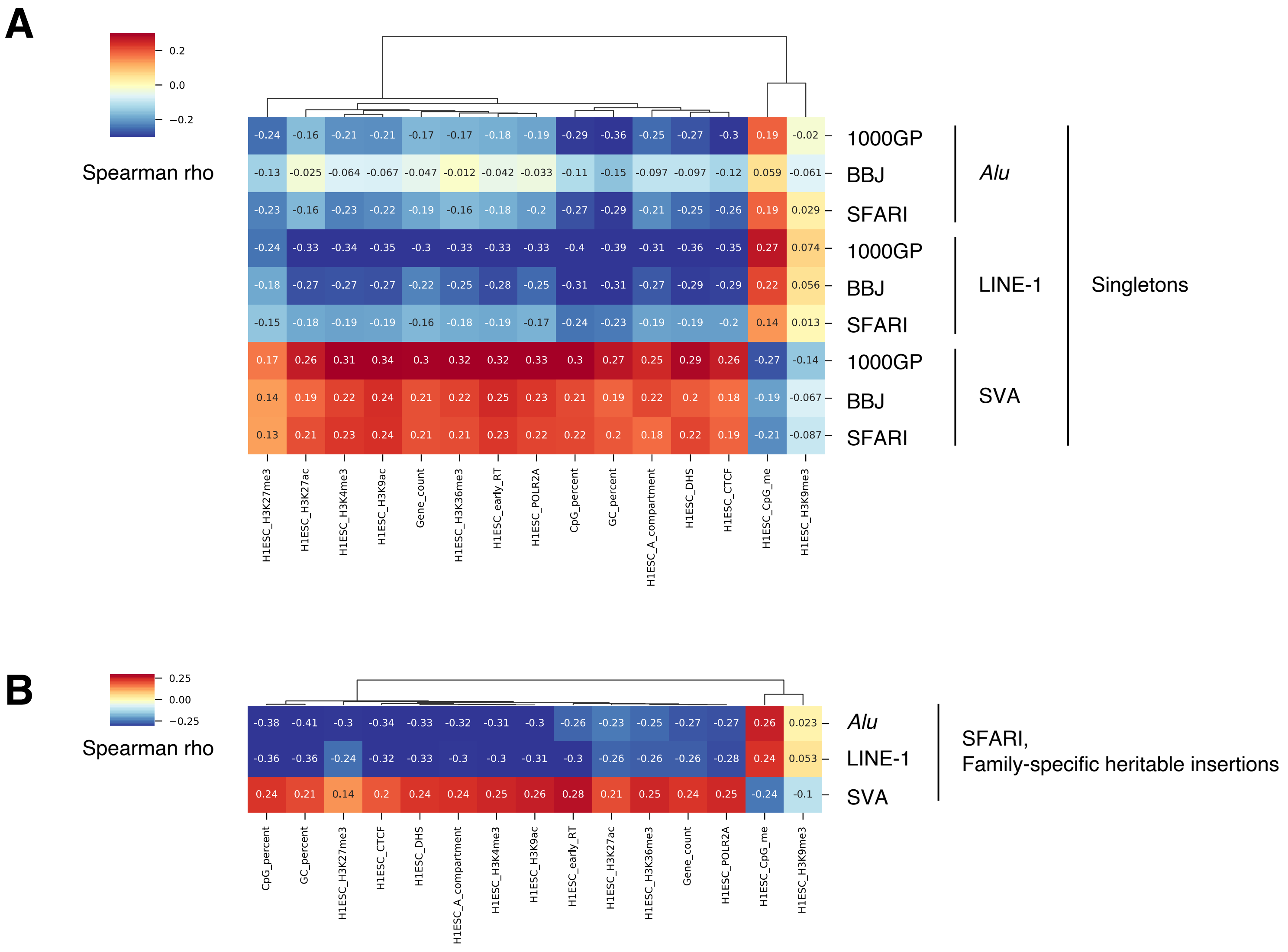
A
Spearman rho
0.2
0.0
−0.2
1000GP
BBJ
SFARI
Alu
LINE-1
SVA
Singletons
H1ESC_H3K27me3
H1ESC_H3K27ac
H1ESC_H3K4me3
H1ESC_H3K9ac
Gene_count
H1ESC_H3K36me3
H1ESC_early_RT
H1ESC_POLR2A
CpG_percent
GC_percent
H1ESC_A_compartment
H1ESC_DHS
H1ESC_CTCF
H1ESC_CpG_me
H1ESC_H3K9me3
B
Spearman rho
0.25
0.00
−0.25
Alu
LINE-1
SVA
SFARI,
Family-specific heritable insertions

### Fig. S33

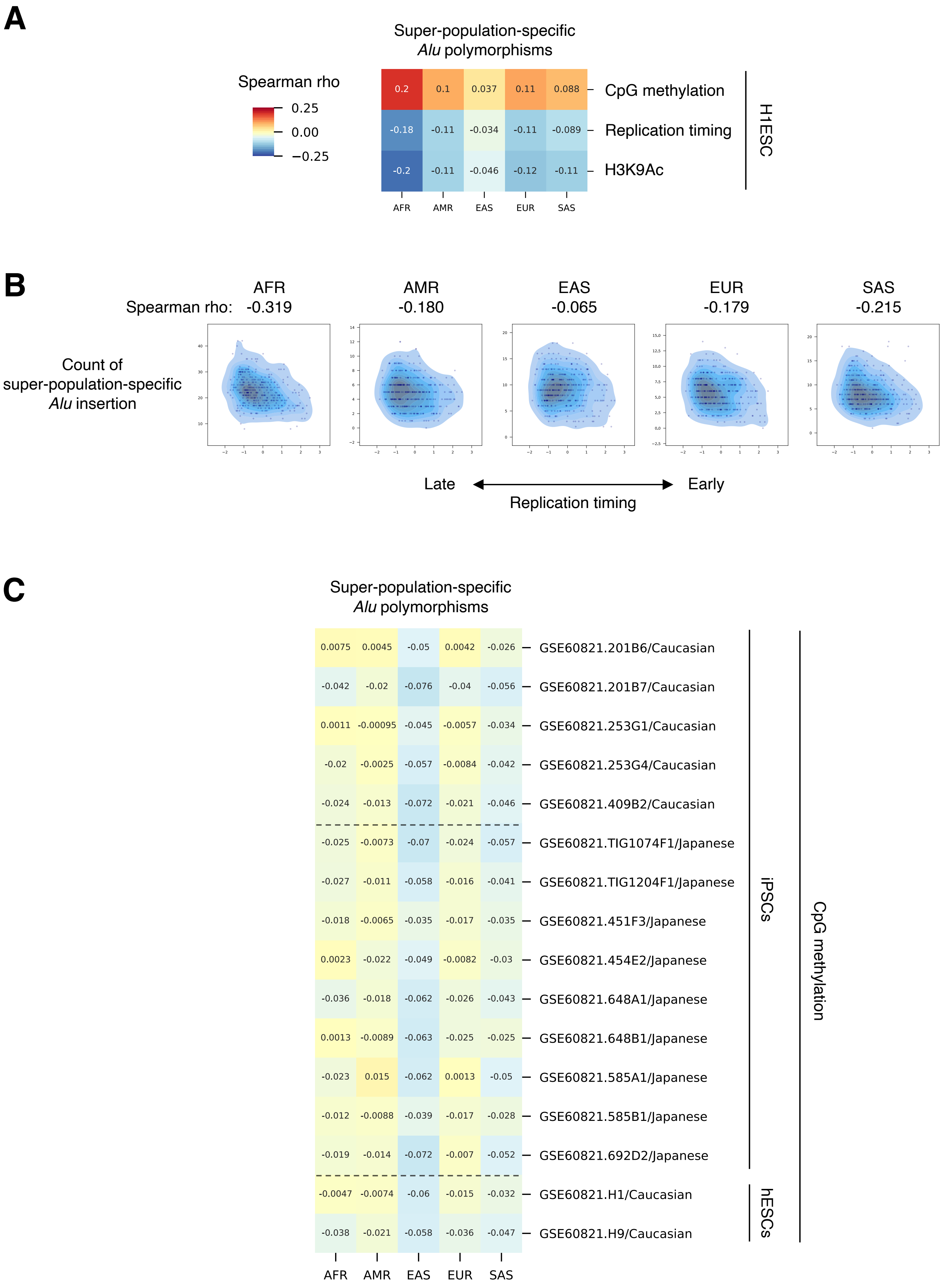
A
Super-population-specific
Alu polymorphisms
Spearman rho
0.25
0.00
−0.25
0.2 0.1 0.037 0.11 0.088 CpG methylation
-0.18 -0.11 -0.034 -0.11 -0.089 Replication timing
-0.2 -0.11 -0.046 -0.12 -0.11 H3K9Ac
H1ESC
AFR AMR EAS EUR SAS
B
Spearman rho:
AFR -0.319
AMR -0.180
EAS -0.065
EUR -0.179
SAS -0.215
Count of
super-population-specific
Alu insertion
Late
Early
Replication timing
C
Super-population-specific
Alu polymorphisms
0.0075 0.0045 -0.05 0.0042 -0.026 GSE60821.201B6/Caucasian
-0.042 -0.02 -0.076 -0.04 -0.056 GSE60821.201B7/Caucasian
0.0011 -0.00095 -0.045 -0.0057 -0.034 GSE60821.253G1/Caucasian
-0.02 -0.0025 -0.057 -0.0084 -0.042 GSE60821.253G4/Caucasian
-0.024 -0.013 -0.072 -0.021 -0.046 GSE60821.409B2/Caucasian
-0.025 -0.0073 -0.07 -0.024 -0.057 GSE60821.TIG1074F1/Japanese
-0.027 -0.011 -0.058 -0.016 -0.041 GSE60821.TIG1204F1/Japanese
-0.018 -0.0065 -0.035 -0.017 -0.035 GSE60821.451F3/Japanese
0.0023 -0.022 -0.049 -0.0082 -0.03 GSE60821.454E2/Japanese
-0.036 -0.018 -0.062 -0.026 -0.043 GSE60821.648A1/Japanese
0.0013 -0.0089 -0.063 -0.025 -0.025 GSE60821.648B1/Japanese
-0.023 0.015 -0.062 0.0013 -0.05 GSE60821.585A1/Japanese
-0.012 -0.0088 -0.039 -0.017 -0.028 GSE60821.585B1/Japanese
-0.019 -0.014 -0.072 -0.007 -0.052 GSE60821.692D2/Japanese
-0.0047 -0.0074 -0.06 -0.015 -0.032 GSE60821.H1/Caucasian
-0.038 -0.021 -0.058 -0.036 -0.047 GSE60821.H9/Caucasian
iPSCs
hESCs
CpG methylation
AFR AMR EAS EUR SAS

### Fig. S34

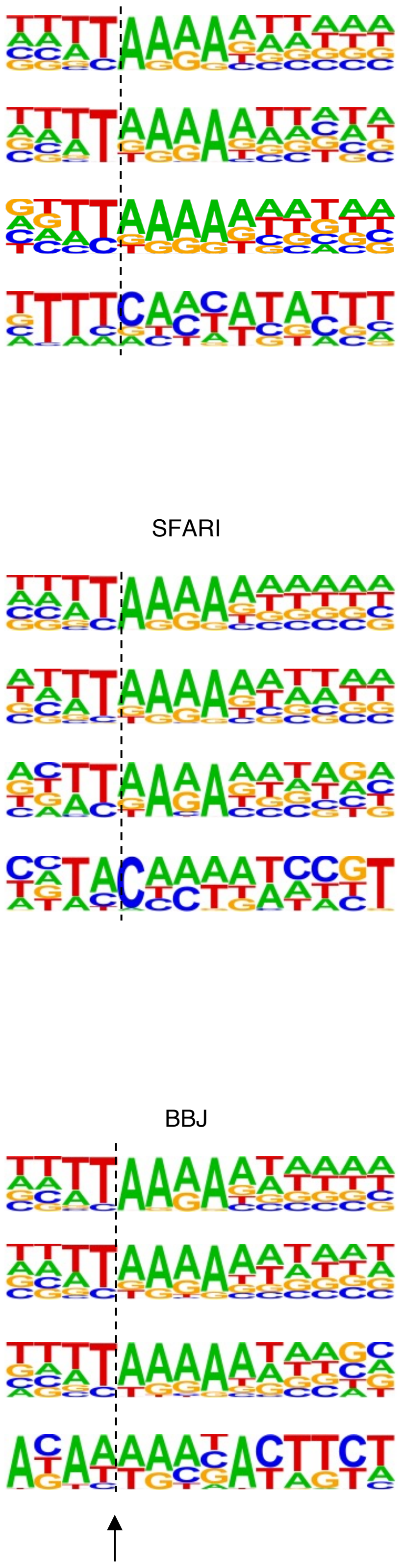

ME insertion breakpoint detected by MEGAnE

**1000GP**

| | Enrichment $P$ | % of target | % of background |
|---|---|---|---|
| *Alu* | 1e-6406 | 40.62 | 5.56 |
| LINE-1 | 1e-552 | 34.33 | 6.53 |
| SVA | 1e-198 | 33.83 | 5.46 |
| HERV-K | 1e-6 (possible false positive) | 34.48 | 2.99 |

**SFARI**

| | Enrichment $P$ | % of target | % of background |
|---|---|---|---|
| *Alu* | 1e-6789 | 39.90 | 5.39 |
| LINE-1 | 1e-631 | 34.78 | 6.40 |
| SVA | 1e-208 | 31.79 | 5.30 |
| HERV-K | 1e-6 (possible false positive) | 22.86 | 0.65 |

**BBJ**

| | Enrichment $P$ | % of target | % of background |
|---|---|---|---|
| *Alu* | 1e-2216 | 34.59 | 4.44 |
| LINE-1 | 1e-404 | 42.13 | 10.09 |
| SVA | 1e-92 | 35.33 | 5.03 |
| HERV-K | 1e-5 (possible false positive) | 33.33 | 0.00 |

### Fig. S35

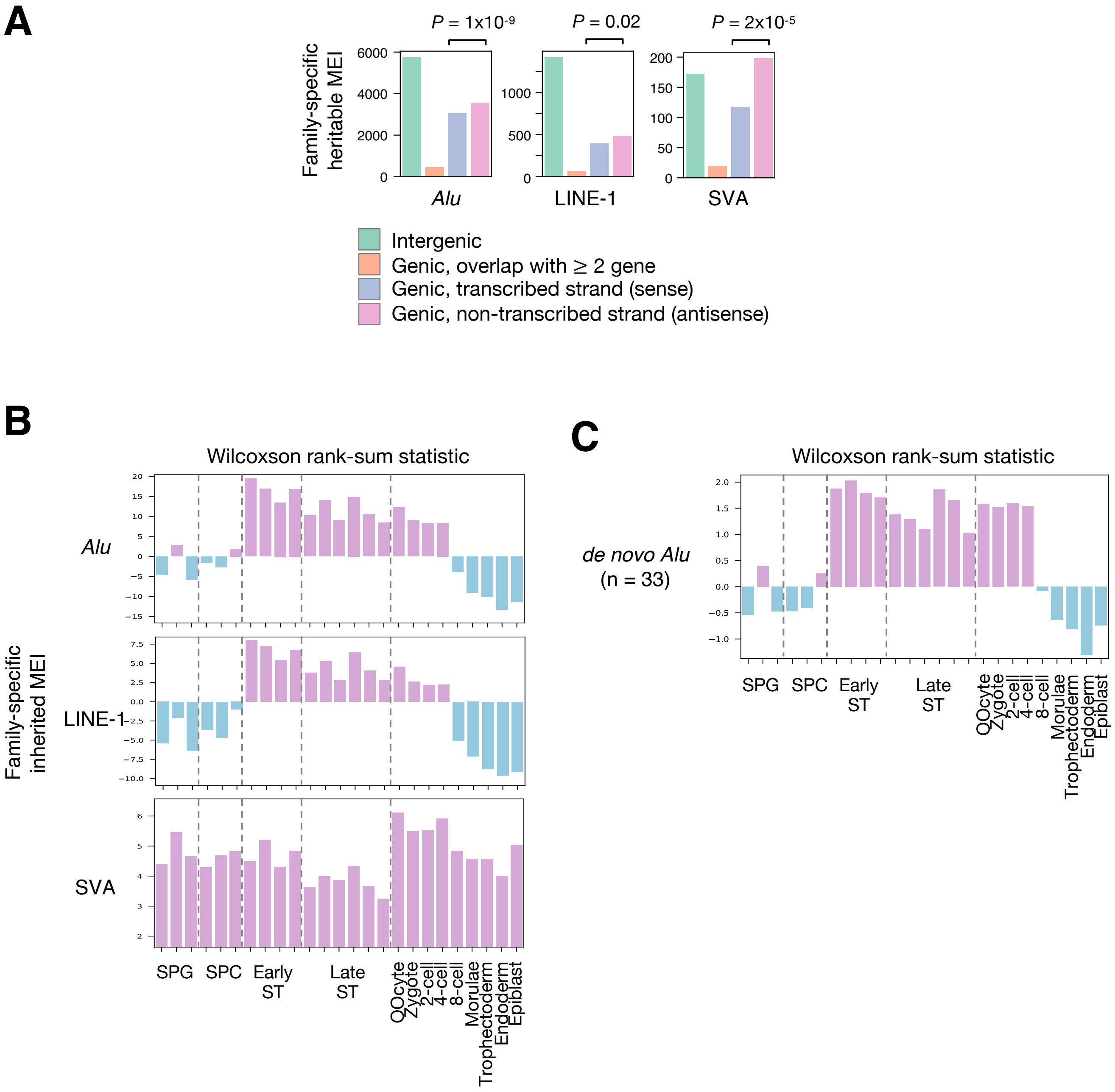

A
P = 1x10-9
P = 0.02
P = 2x10-5
Family-specific heritable MEI
Alu
LINE-1
SVA
Intergenic
Genic, overlap with ≥ 2 gene
Genic, transcribed strand (sense)
Genic, non-transcribed strand (antisense)
B
Wilcoxson rank-sum statistic
Family-specific inherited MEI
Alu
LINE-1
SVA
SPG
SPC
Early ST
Late ST
OOcyte
Zygote
2-cell
4-cell
8-cell
Morulae
Trophectoderm
Endoderm
Epiblast
C
Wilcoxson rank-sum statistic
de novo Alu (n = 33)

### Fig. S36

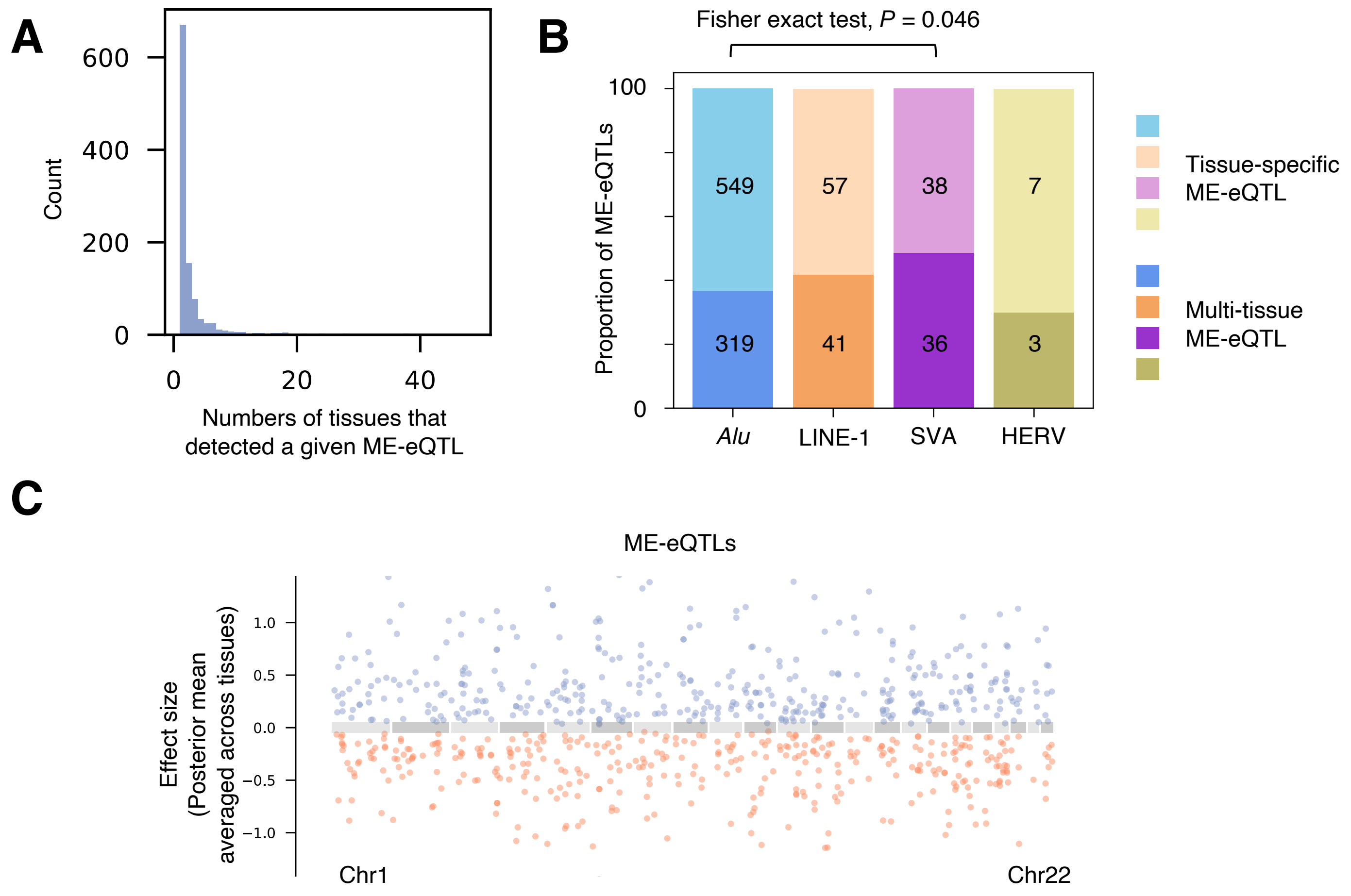
A
Count
600
400
200
0
0
20
40
Numbers of tissues that detected a given ME-eQTL
B
Fisher exact test, P = 0.046
Proportion of ME-eQTLs
100
0
549
57
38
7
319
41
36
3
Alu
LINE-1
SVA
HERV
Tissue-specific ME-eQTL
Multi-tissue ME-eQTL
C
ME-eQTLs
Effect size (Posterior mean averaged across tissues)
1.0
0.5
0.0
−0.5
−1.0
Chr1
Chr22

### Fig. S37

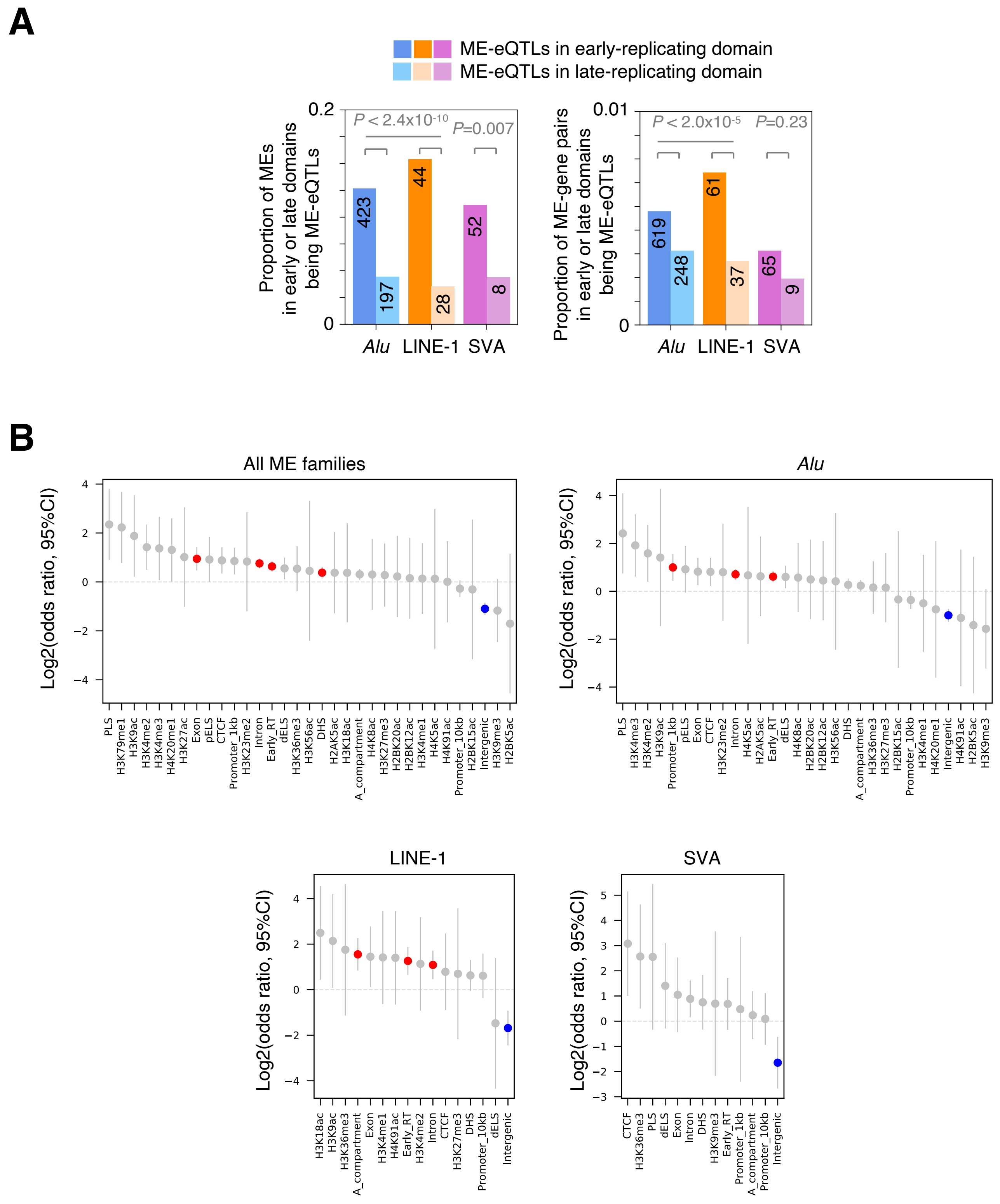

A
ME-eQTLs in early-replicating domain
ME-eQTLs in late-replicating domain
Proportion of MEs in early or late domains being ME-eQTLs
P < 2.4x10-10
P=0.007
423
197
44
28
52
8
Alu
LINE-1
SVA
Proportion of ME-gene pairs in early or late domains being ME-eQTLs
P < 2.0x10-5
P=0.23
619
248
61
37
65
9
B
All ME families
Alu
LINE-1
SVA
Log2(odds ratio, 95%CI)

### Fig. S38

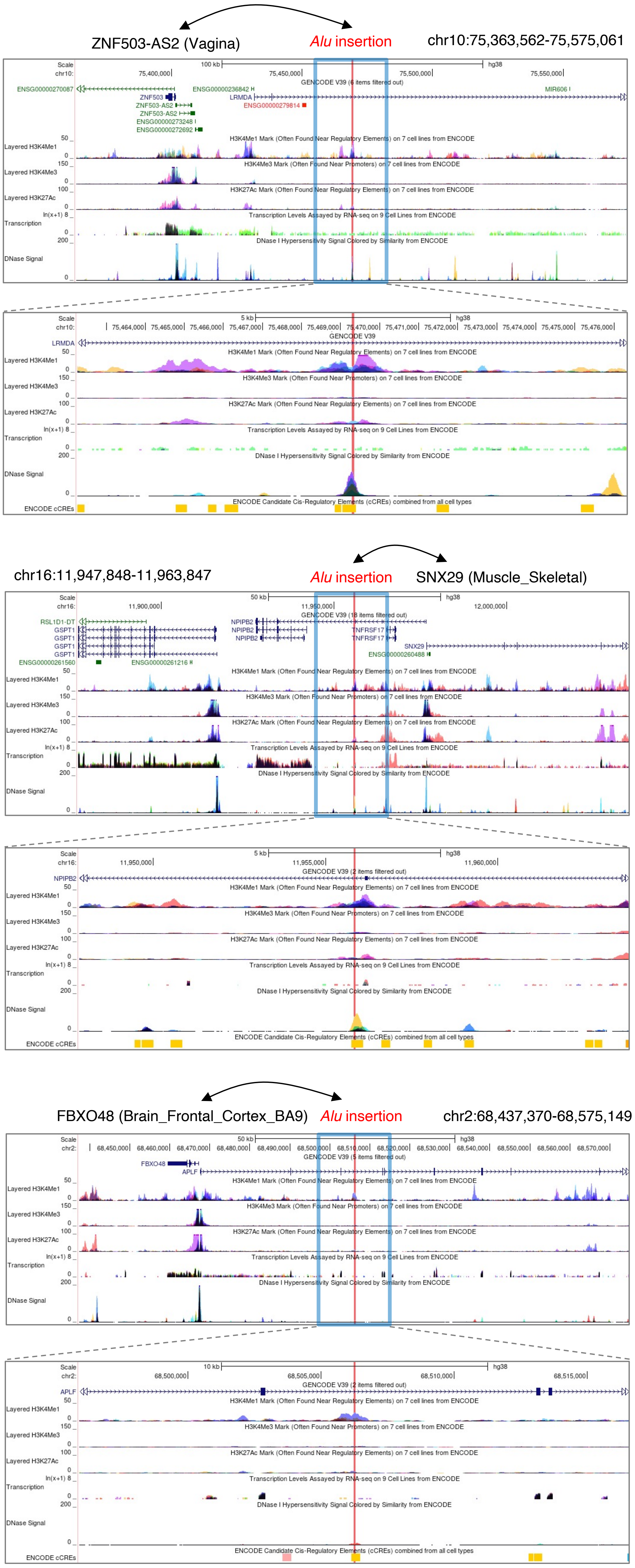

ZNF503-AS2 (Vagina)
Alu insertion
chr10:75,363,562-75,575,061
chr16:11,947,848-11,963,847
Alu insertion
SNX29 (Muscle_Skeletal)
FBXO48 (Brain_Frontal_Cortex_BA9)
Alu insertion
chr2:68,437,370-68,575,149

### Fig. S39

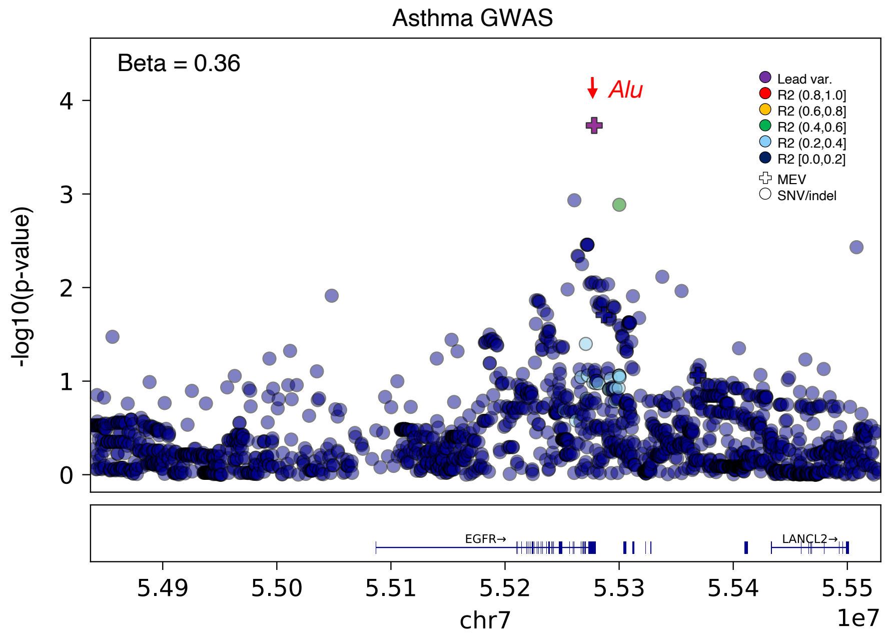
Asthma GWAS
Beta = 0.36
Alu
Lead var.
R2 (0.8,1.0]
R2 (0.6,0.8]
R2 (0.4,0.6]
R2 (0.2,0.4]
R2 [0.0,0.2]
MEV
SNV/indel
-log10(p-value)
0
1
2
3
4
EGFR→
LANCL2→
5.49
5.50
5.51
5.52
5.53
5.54
5.55
1e7
chr7

### Fig. S40

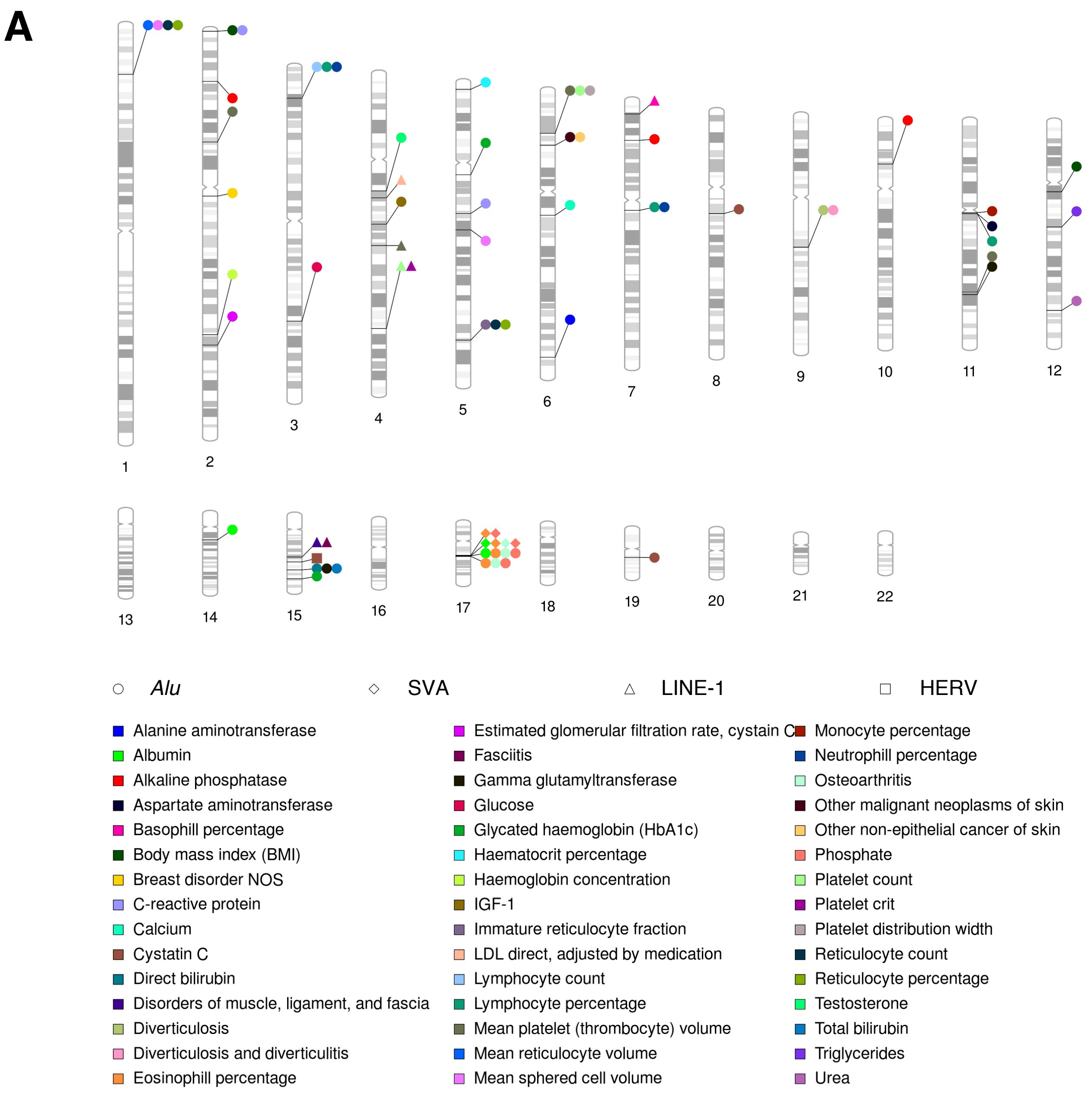
A
Alu
SVA
LINE-1
HERV
Alanine aminotransferase
Albumin
Alkaline phosphatase
Aspartate aminotransferase
Basophill percentage
Body mass index (BMI)
Breast disorder NOS
C-reactive protein
Calcium
Cystatin C
Direct bilirubin
Disorders of muscle, ligament, and fascia
Diverticulosis
Diverticulosis and diverticulitis
Eosinophill percentage
Estimated glomerular filtration rate, cystain C
Fasciitis
Gamma glutamyltransferase
Glucose
Glycated haemoglobin (HbA1c)
Haematocrit percentage
Haemoglobin concentration
IGF-1
Immature reticulocyte fraction
LDL direct, adjusted by medication
Lymphocyte count
Lymphocyte percentage
Mean platelet (thrombocyte) volume
Mean reticulocyte volume
Mean sphered cell volume
Monocyte percentage
Neutrophill percentage
Osteoarthritis
Other malignant neoplasms of skin
Other non-epithelial cancer of skin
Phosphate
Platelet count
Platelet crit
Platelet distribution width
Reticulocyte count
Reticulocyte percentage
Testosterone
Total bilirubin
Triglycerides
Urea

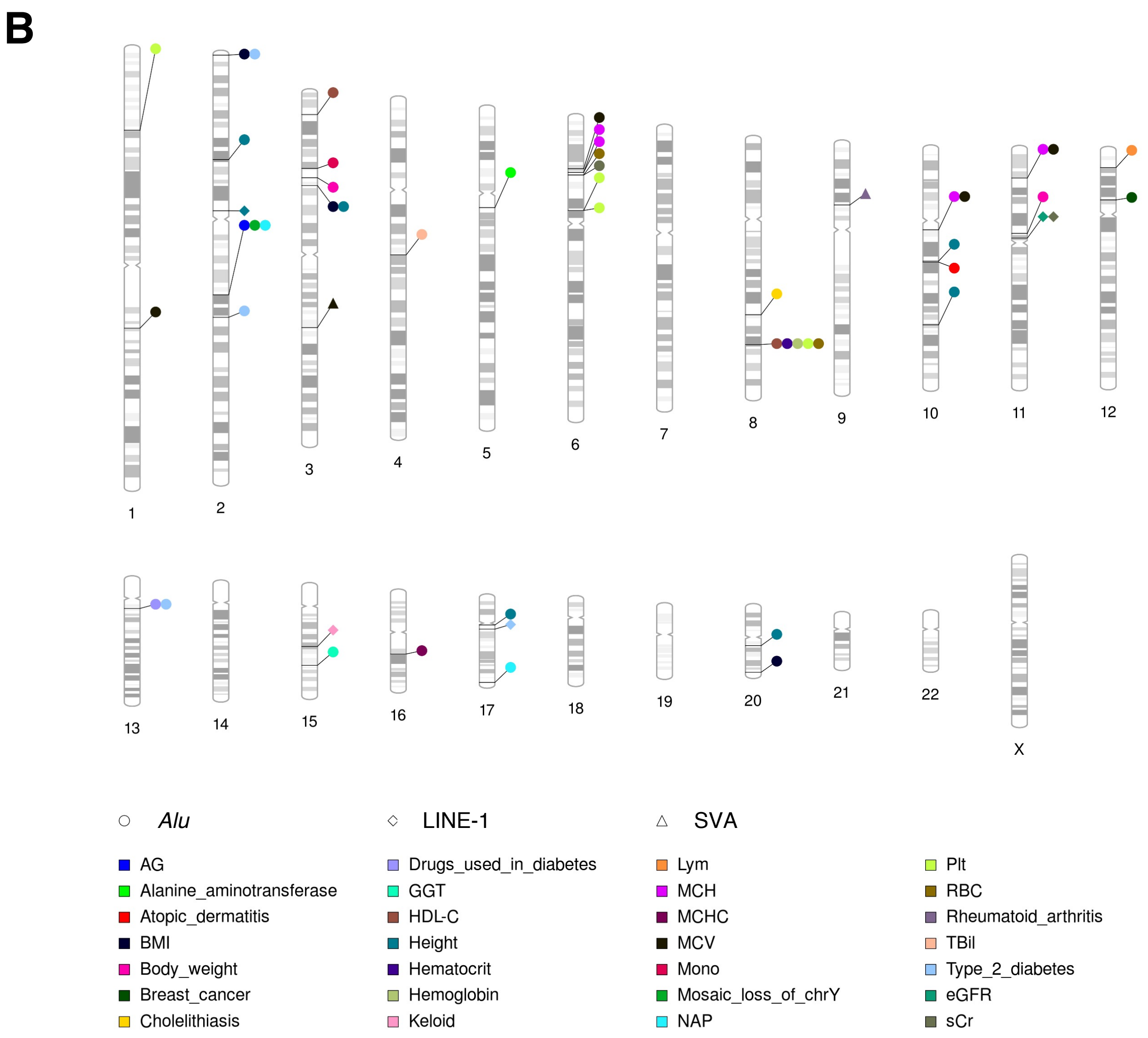
B
X
Alu
LINE-1
SVA
AG
Alanine_aminotransferase
Atopic_dermatitis
BMI
Body_weight
Breast_cancer
Cholelithiasis
Drugs_used_in_diabetes
GGT
HDL-C
Height
Hematocrit
Hemoglobin
Keloid
Lym
MCH
MCHC
MCV
Mono
Mosaic_loss_of_chrY
NAP
Plt
RBC
Rheumatoid_arthritis
TBil
Type_2_diabetes
eGFR
sCr

### Fig. S41

**A**

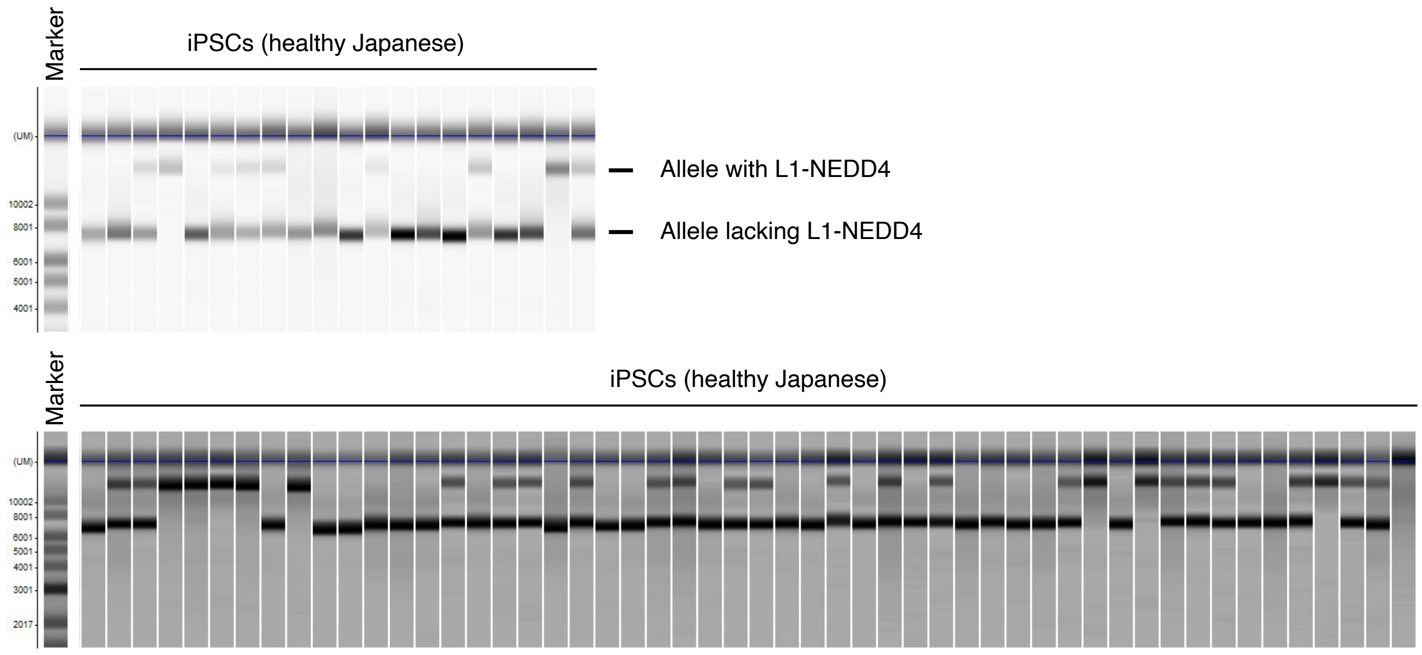
Marker
iPSCs (healthy Japanese)
(UM)
10002
8001
6001
5001
4001
Allele with L1-NEDD4
Allele lacking L1-NEDD4
Marker
iPSCs (healthy Japanese)
(UM)
10002
8001
6001
5001
4001
3001
2017

**B**

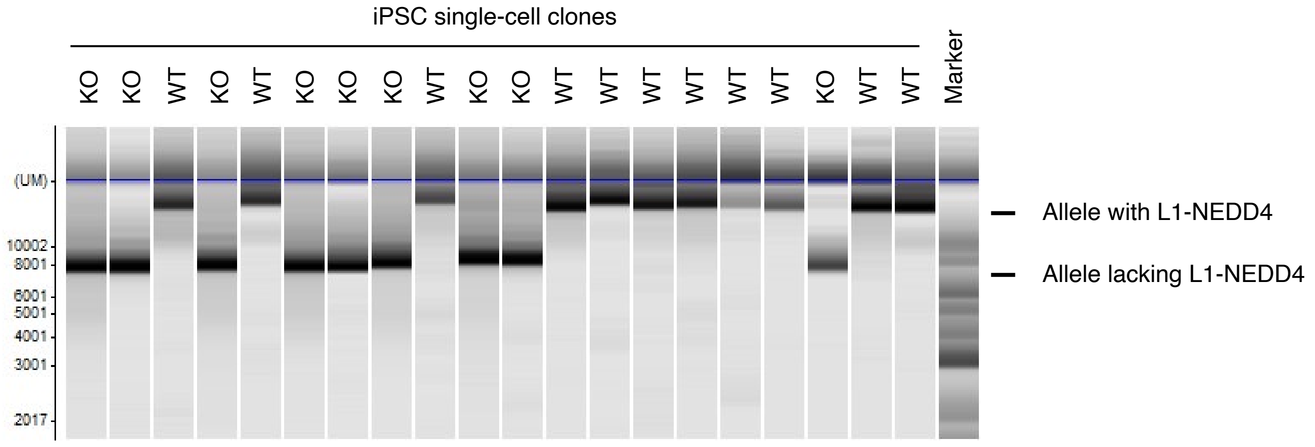
iPSC single-cell clones
KO KO WT KO WT KO KO KO WT KO KO WT WT WT WT WT WT KO WT WT
Marker
(UM)
10002
8001
6001
5001
4001
3001
2017
Allele with L1-NEDD4
Allele lacking L1-NEDD4

### Fig. S42

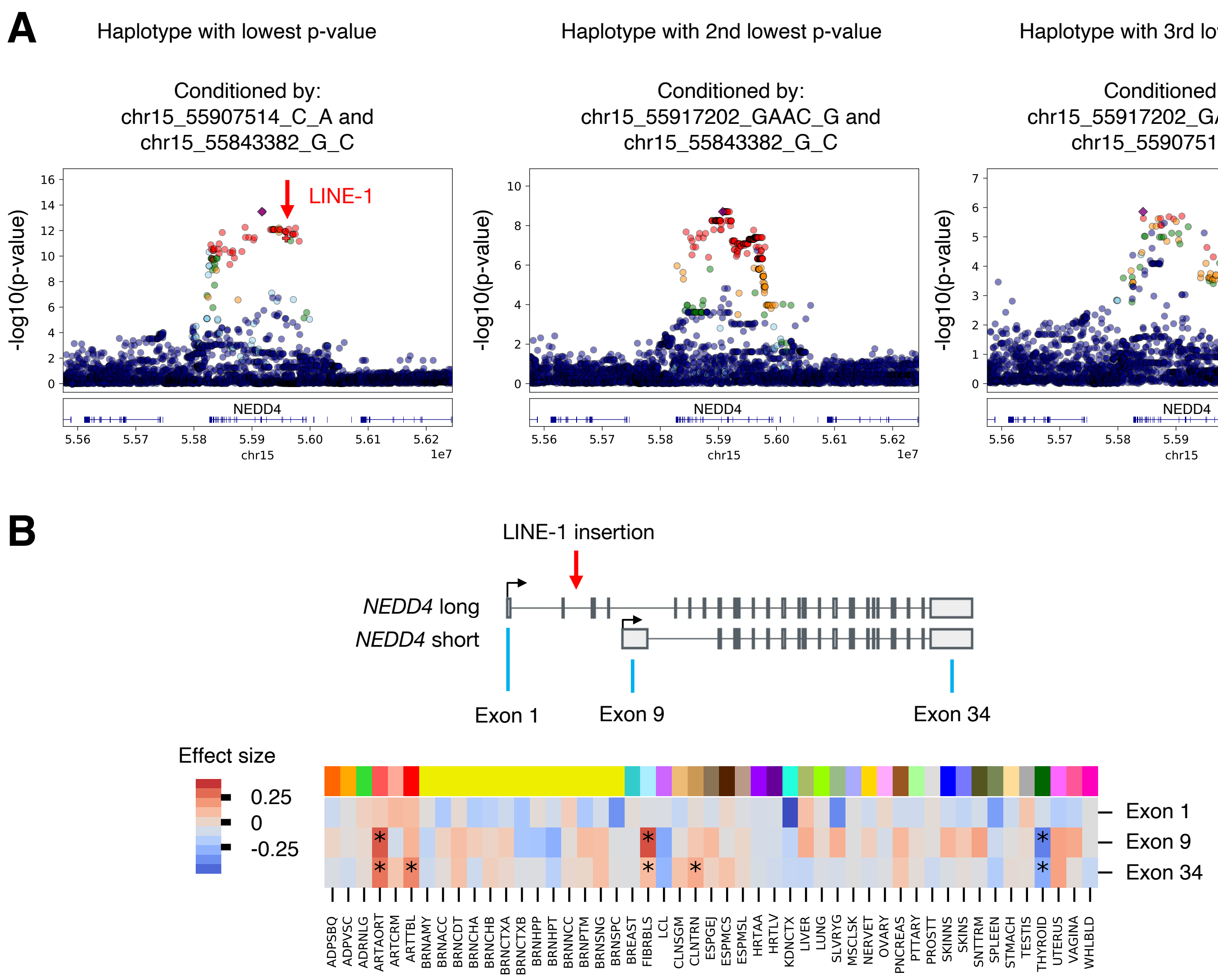
A
Haplotype with lowest p-value
Conditioned by:
chr15_55907514_C_A and
chr15_55843382_G_C
Haplotype with 2nd lowest p-value
Conditioned by:
chr15_55917202_GAAC_G and
chr15_55843382_G_C
Haplotype with 3rd lowest p-value
Conditioned by:
chr15_55917202_GAAC_G and
chr15_55907514_C_A
-log10(p-value)
LINE-1
NEDD4
chr15
1e7
B
LINE-1 insertion
NEDD4 long
NEDD4 short
Exon 1
Exon 9
Exon 34
Effect size
0.25
0
-0.25
Exon 1
Exon 9
Exon 34
ADPSBQ ADPVSC ADRNLG ARTAORT ARTCRM ARTTBL BRNAMY BRNACC BRNCDT BRNCHA BRNCHB BRNCTXA BRNCTXB BRNHPP BRNHPT BRNNCC BRNPTM BRNSNG BRNSPC BREAST FIBRBLS LCL CLNSGM CLNTRN ESPGEJ ESPMCS ESPMSL HRTAA HRTLV KDNCTX LIVER LUNG SLVRYG MSCLSK NERVET OVARY PNCREAS PTTARY PROSTT SKINNS SKINS SNTTRM SPLEEN STMACH TESTIS THYROID UTERUS VAGINA WHLBLD

### Fig. S43

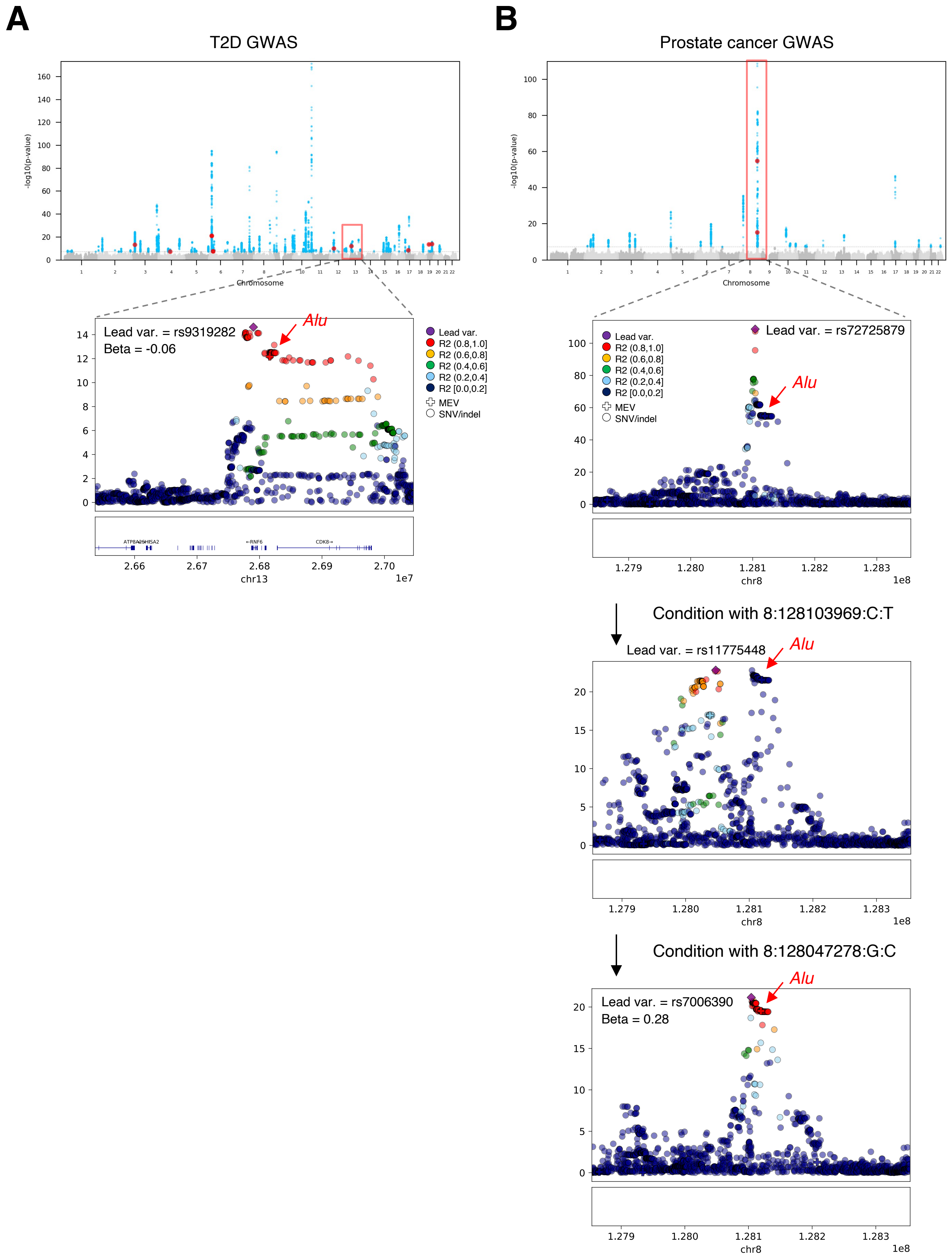
A
T2D GWAS
-log10(p-value)
Chromosome
Lead var. = rs9319282
Beta = -0.06
Alu
Lead var.
R2 (0.8,1.0]
R2 (0.6,0.8]
R2 (0.4,0.6]
R2 (0.2,0.4]
R2 [0.0,0.2]
MEV
SNV/indel
chr13
1e7
B
Prostate cancer GWAS
-log10(p-value)
Chromosome
Lead var. = rs72725879
Alu
chr8
1e8
Condition with 8:128103969:C:T
Lead var. = rs11775448
Alu
chr8
1e8
Condition with 8:128047278:G:C
Lead var. = rs7006390
Beta = 0.28
Alu
chr8
1e8

### Fig. S44

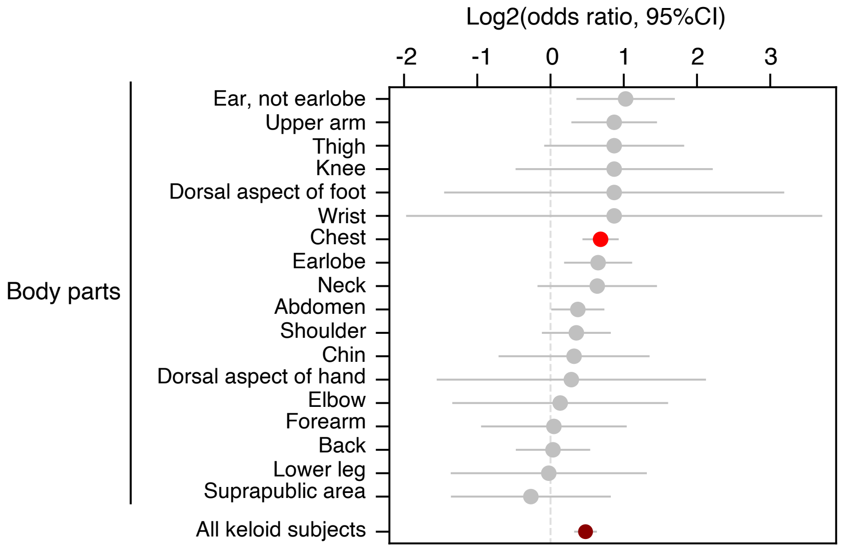

Log2(odds ratio, 95%CI)
-2
-1
0
1
2
3
Body parts
Ear, not earlobe
Upper arm
Thigh
Knee
Dorsal aspect of foot
Wrist
Chest
Earlobe
Neck
Abdomen
Shoulder
Chin
Dorsal aspect of hand
Elbow
Forearm
Back
Lower leg
Suprapublic area
All keloid subjects
